# Single-cell phenomics reveals behavioural and mechanical heterogeneities underpinning collective migration during mouse anterior patterning

**DOI:** 10.1101/2023.03.31.534937

**Authors:** Matthew Stower, Felix Zhou, Holly Hathrell, Jason Yeung, Shifaan Thowfeequ, Jonathan Godwin, Falk Schneider, Christoffer Lagerholm, Marco Fritzsche, Jeyan Thiyagalingam, Xin Lu, Jens Rittscher, Shankar Srinivas

## Abstract

Distal Visceral Endoderm (DVE) cells show a stereotypic unidirectional migration essential for correct orientation of the anterior-posterior axis. They migrate within a simple epithelium, the Visceral Endoderm (VE). It is unknown how DVE cells negotiate their way amongst the surrounding VE cells, what determines the bounds of DVE migration within the VE, and the relative contributions of different cell behaviours to this migration. To address these questions, we used lightsheet microscopy to generate a multi-embryo, singlecell resolution, longitudinal dataset of cell behaviour and morphology. We developed a machine learning based pipeline to segment cells and a data-informed systematic computational framework to extract and compare select morphological, behavioural and molecular parameters of all VE cells in a unified coordinate space. Unbiased clustering of this single-cell ‘phenomic’ dataset reveals considerable patterned phenotypic heterogeneity within the VE and a previously unknown sub-grouping within the DVE. While migrating, DVE cells retain regular morphology, do not exchange neighbours and are crowded, all hallmarks of the jammed state. In contrast, VE cells immediately ahead of them deform and undergo neighbour exchange. We show that DVE cells are characterised by higher levels of apical F-actin and elevated tension relative to the VE cells immediately ahead of them through which they migrate, but stop migrating upon reaching a region of the VE with matching elevated tension. *Lefty1* mutants, known to show abnormal over-migration of DVE cells, show disruption to this patterned tension in the VE. Our findings provide novel insights into the control of cell behaviour during the remodelling of curved epithelia, indicating that the collective migration of sub-sets of cells can be circumscribed by modulating the mechanical properties of surrounding cells and that migrating cells in this context remain as a jammed solid flock, with surrounding cells facilitating their movement by becoming unjammed.

**Graphical Abstract:** 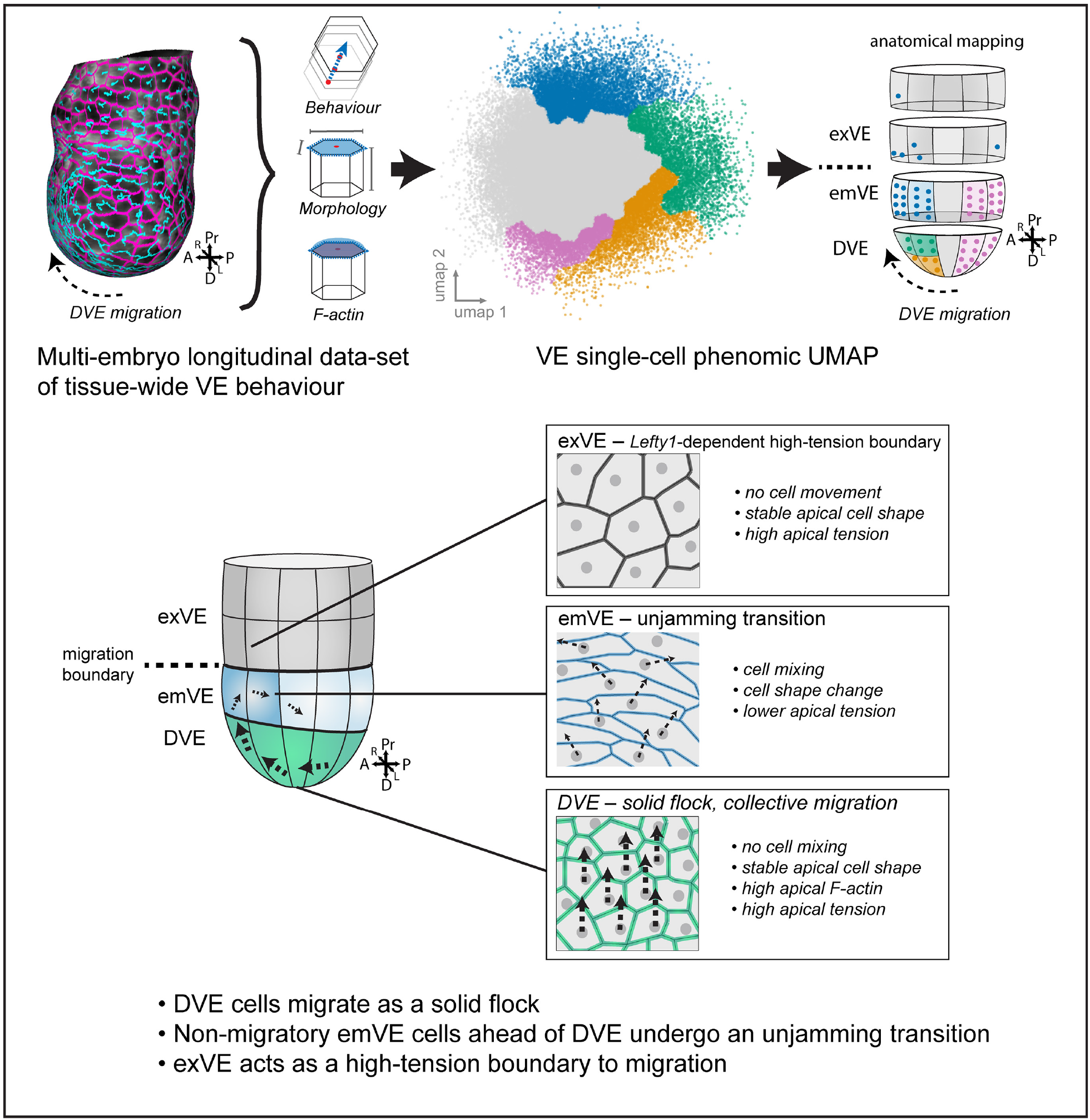

## INTRODUCTION

The dynamic remodelling of epithelial sheets requires the movement of interconnected cells through spatially localised, coordinated cell behaviours. Live imaging of epithelia during cell migration has revealed that cells can employ a variety of behaviours including T1 transitions (Bertet et al. 2004), or by forming transient multicellular ‘rosette’ structures that resolve in a directional manner to facilitate local neighbour exchange events (Blankenship et al. 2006). Cells in the tissue can also undergo changes in morphology, regulated patterns of divisions, and apoptosis, all of which must be coordinated to enable movements while retaining mechanical tissue integrity (Economou et al. 2013). In order to initiate movements in an epithelial context it has been suggested that the tissue undergoes a transition from a rigid, packed state, where cell movements are constrained (‘jammed’), to a fluid-like state where cell movements are permissible (‘unjammed’) (Angelini et al. 2011; Bi et al. 2011; Bi et al. 2015; Bi et al. 2016; Cates et al. 1998; Lawson-Keister and Manning 2021; Park et al. 2016; Trappe et al. 2001). In the classical formulation, particles in a jammed state can be forced to become unjammed by even small changes in the direction of stresses applied on them (Cates et al. 1998; Liu and Nagel 1998). In the biological context, cells can come to occupy a jammed state for a variety of reasons, with different mechanisms being responsible depending on biological context (Lawson-Keister and Manning 2021).

The onset of collective cell movements in a confluent cell layer has been described as a flocking transition, with parallels to the behaviour of birds flying together in large groups (Giavazzi et al. 2018) (Szabo et al. 2006; Vicsek and Zafeiris 2012). In such a model, rather than leader cells ‘pulling’ the cells behind them forward, it is the alignment of interactions between individual cells that promotes directional movement (Giavazzi et al. 2018; Malinverno et al. 2017), as cells far from the leading edge can also provide lamellipodial-based tractional force in cell culture experiments (Trepat et al. 2009). Modelling this process with Self-Propelled Voronoi models indicate the existence of liquid-flock and solid-flock states, where cells migrate with or without cell mixing, respectively (Giavazzi et al. 2018; Trepat and Sahai 2018). Such flocking behaviour can be induced in confluent human mammary epithelial cells by over expression of Rab5a, thought to be due to stimulating junctional remodelling (Malinverno et al. 2017).

Much of this work however has been carried out on relatively flat epithelia, *in vitro* two-dimensional (2D) flat culture or mathematical models of flat tissues. However *in vivo*, epithelial tissues exist in complex three-dimensional (3D) architectures, adding potentially different topological and mechanical settings across the varying spatiotemporal scales of the tissue that migrating cells need to negotiate (Li et al. 2021).

One such example is the Visceral Endoderm (VE), a monolayer epithelium in the developing mouse embryo arranged as a blunt-ended cylinder (often referred to as the *egg cylinder)*. A sub-set of VE cells, termed the Distal Visceral Endoderm (DVE) show a characteristic migratory behaviour within the plane of this curved epithelial tissue. DVE cells are induced at the distal-tip of the embryonic day 5.5 (E5.5) mouse embryo (Thomas et al. 1998) (Figure 1A) where they express a specific set of genes (Belo et al. 1997; Pfister et al. 2007; Thomas et al. 1998; Thowfeequ et al. 2021; Varlet et al. 1997; Yamamoto et al. 2004) and migrate proximally over the course of 3-5 hours in a unidirectional manner (Srinivas et al. 2004; Thomas et al. 1998). DVE cells stop at a boundary, defined by that of the underlying embryonic (Em) epiblast and extra-embryonic ectoderm (ExE) (the ‘Em-Ex boundary’), so that they are positioned asymmetrically along one side of the embryo, at which point they are termed Anterior Visceral Endoderm (AVE). DVE cells express *Lefty1, Cerl1* and *Dkk1*, inhibitors of the TGF-β/NODAL and WNT pathways (Belo et al. 1997; Yamamoto et al. 2004) that restrict the formation of the primitive-streak to the opposite (future ‘posterior’) side of the epiblast at E6.25. *Lefty1* mutant embryos show an abnormal over-migration of DVE cells into the region of VE overlying the ExE (Trichas et al. 2011), indicating that regulation of TGF-β signalling in this context is important for regulating migratory behaviour.

Live imaging has shown that DVE cell movement is an active process as cells produce polarised cellular projections from their basal aspect in the direction of migration (Migeotte et al. 2010; Srinivas et al. 2004) Furthermore, knock-outs of genes involved in actin regulation lead to loss of these cellular projections and disruption of migratory behaviour (Migeotte et al. 2010; Omelchenko et al. 2014; Rakeman and Anderson 2006).

Our understanding of the cellular basis of DVE migration is, nevertheless, incomplete. Studies typically focus only on the actively migrating cells, rather than considering the global epithelial context (i.e., cells positioned lateral and posterior to the DVE) within which migration occurs. Therefore we lack an integrated understanding of the regional cell behaviours (e.g., movement, changes in cell shape, oriented cells division events, cell-cell rearrangements) that underlie DVE migration – how, in short, the interactions at the level of individual cells lead to emergent tissue-level morphogenetic behaviour. Furthermore, while this migration is often described as a collective migration (Bloomekatz et al. 2012; Migeotte et al. 2011; Morris et al. 2012; Shioi et al. 2017), no study to date has tracked all DVE cells in a single embryo during their migration, so it remains unclear whether differences exist within the migratory cell population as they migrate through the plane of the epithelial monolayer.

Although relatively little is known about the cellular properties that distinguish migratory DVE cells from surrounding non-migratory VE cells, it is known that NODAL signalling from the epiblast is required for DVE induction (Brennan et al. 2001), and that this triggers cells to transition from squamous to columnar morphology prior to migration (Kimura et al. 2000; Rivera-Perez et al. 2003; Srinivas et al. 2004). It is understood that DVE cells are induced at the distal tip as the embryo grows and elongates, distancing the distal cells of the VE from repressive BMP signals emanating from the ExE (Mesnard et al. 2006; Rodriguez et al. 2005). Nevertheless, as DVE cells are positioned at the distal-tip of the cylinder-shaped VE, the region with the highest tissue curvature, these cells may also be subject to increased mechanical stress (Helfrich 1973; Khairy et al. 2018; Khmelinskii and Makarov 2020). As mechanical cues can affect cell fate decisions and cell behaviour (De Belly et al. 2022; Discher et al. 2009), it is unclear whether the columnar morphology and behaviour of the migratory DVE cells is due to biomechanical differences imposed by the curvature of the VE tissue, or autonomously controlled.

Consequently, identifying the key parameters underlying tissue morphogenesis requires analysis of the spatiotemporal heterogeneity in morphological and behavioural characteristics (phenotypes) of all the cells within the tissue at the level of the constituent single-cells. Importantly, for statistical robustness, these measurements require a framework to integrate such quantitative data from across multiple embryos, due to the nature of their heterogeneity. Quantitative time-lapse data in the mouse embryo can be obtained using lightsheet-based imaging and cell tracking (Ichikawa et al. 2013; McDole et al. 2018; Udan et al. 2014), however, such efforts so far have used only nuclear fluorescent reporters (e.g. H2B-GFP) that are relatively easy to segment and can be used for positional cell tracking, but do not provide information about cell-shape, surface-area or other cell morphology specific phenotypic parameters.

In order to address these limitations, we developed a longitudinal imaging based-approach for recording cell shape and movements at high spatial and temporal resolution during DVE migration, using lightsheet microscopy. To integrate data on the dynamics of cellular phenotypes from multiple embryos with varying numbers of cells and durations of DVE migration, we developed a method for the temporal staging and spatial alignment of embryos based on DVE migration. To enable systematic comparative analysis of phenotype between these large multi-dimensional image volumes, we developed innovative computational tools to map the apical surface of every VE cell at each time-point onto a single volumetric coordinate system, so that we could project it into 2D and use machine learning-aided approaches to segment individual cells for temporal tracking. To analyse these multi-dimensional data, we developed a multivariate single-cell manifold analysis that integrates cells from multiple embryos developing across time. Such approaches have recently been applied with good effect to fixed samples to study morphological transformations (Andrews et al. 2021). The extension of these approaches to longitudinal data enabled us to comprehensively profile and compare the morphological and behavioural phenotype of all VE cells during DVE migration, at single-cell resolution.

This single cell ‘phenomic’ (Davis 1949) characterisation revealed a previously unappreciated, patterned, heterogeneity within the VE. DVE cells can be categorised into distinct subgroups on the basis of their phenomes, while cells in the proximal emVE (Figure 1 A) also form distinct phenomic clusters. Tracking cells throughout the migration phase revealed that DVE cells remain relatively constant in morphology, do not exchange neighbours and are crowded, with a constant low apical surface area. In contrast, VE cells immediately ahead of them show key hallmarks of a phase transition to an unjammed state, changing cell shape to become irregular and undergoing cell mixing, enabling them to be displaced anteriorly and laterally. In addition to being distinguished from surrounding VE cells by their migratory behaviour and morphology, DVE cells also show elevated levels of apical and junctional F-actin, suggesting that they are also mechanically distinct from surrounding cells. We verified that DVE cells are characterised by a significantly elevated membrane tension, and that this tension is dependent on actomyosin contractile activity. DVE cells migrate through cells with significantly lower apical tension and halt their migration upon reaching the VE overlying the ExE, which has an apical tension comparable to the DVE. In contrast, *Lefty1* mutants show a significant reduction of tension in the VE overlying the ExE, into which DVE cells abnormally over-migrate.

**Figure 1.**
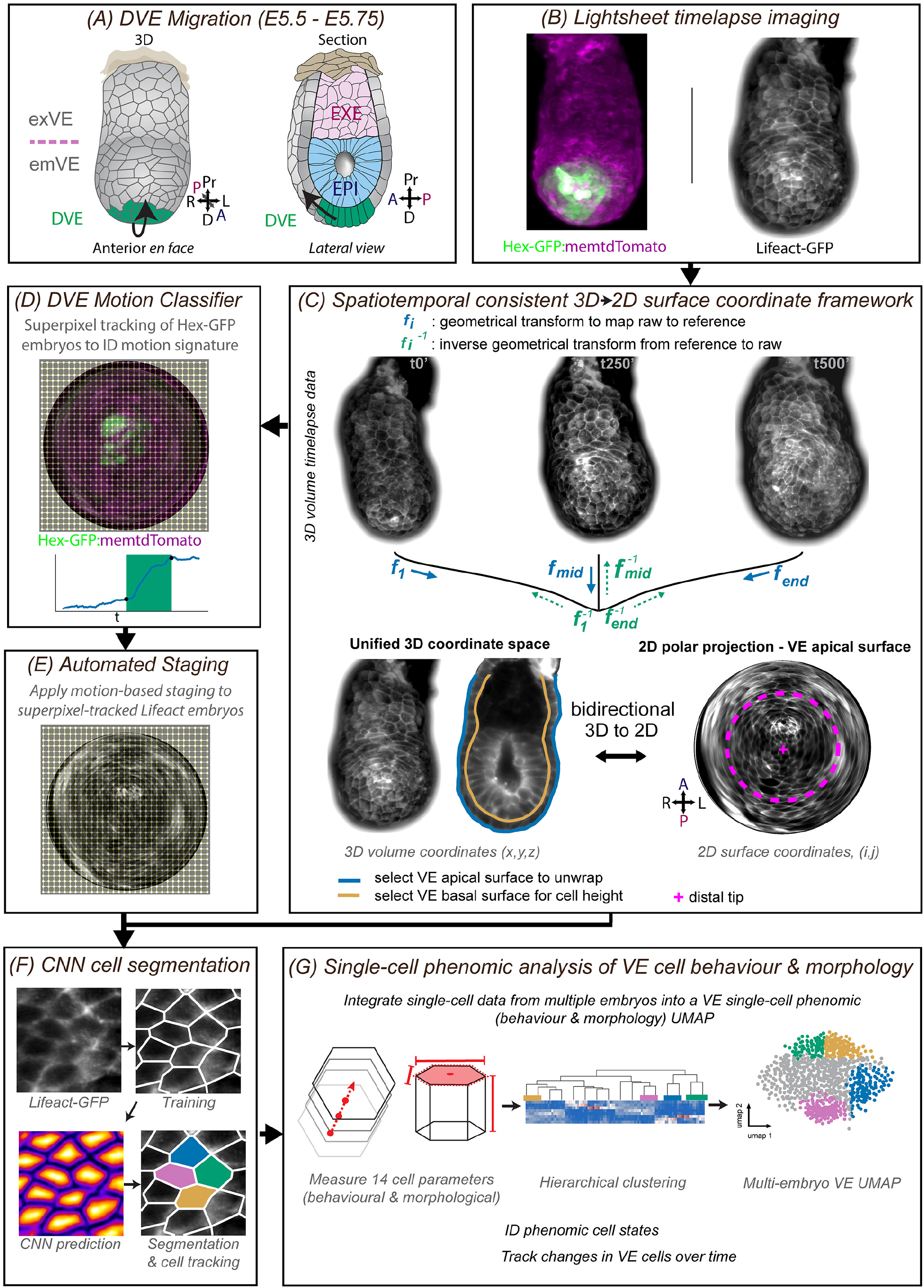
Data processing framework for single-cell, multi-embryo analysis of VE cell behaviour and morphology during DVE migration. **A.** En face and cut-away profile illustrations of the E5.5 mouse embryo showing the monolayer visceral endoderm (VE) surrounding the epiblast (EPI) and extra-embryonic ectoderm (ExE). The migratory distal visceral endoderm (DVE) cells are in green. Arrows indicate direction of migration to future anterior. **B.** Representative images of a single time point from lightsheet imaged Hex-GFP:membrane-tdTomato and Lifeact-GFP mouse embryos. **C-G.** Computational pipeline to extract and analyse quantitative single cell phenomic data from multidimensional image data. **C.** Spatiotemporal registration of the 3D time-lapse data to generate a consistent surface coordinate framework, to enable 2D geodesic surface projection of the apical VE surface. **D.** A DVE motion classifier trained using superpixel motion tracking of DVE cells labelled with Hex-GFP. **E.** The trained motion-based classifier is used to identify the migration phase of Lifeact-GFP embryos. **F.** Convolutional neural network to identify and segment individual cells from Lifeact-GFP time-lapse data. **G.** Single-cell analysis of the phenome (morphology and behaviour) of VE cells integrating time-lapse data from multiple embryos.

These results reveal a novel model for DVE migration, where they migrate as a solid flock within the epithelium of the VE, facilitated by the unjamming of cells immediately surrounding them, and delimited by patterned tension differentials within the epithelium.

## RESULTS

### E5.5 multi-embryo 5D lightsheet time-lapse dataset of DVE migration

To create a library of time-lapse datasets that capture at cellular resolution dynamic events in the VE during DVE migration and enable the extraction of parameters relating to cell morphology, and movements we optimised the culture and imaging of egg-cylinder stage embryos on the ZEISS Z.1 lightsheet microscope (Figure S1A-C).

To ensure that our culture conditions and imaging parameters permitted normal embryo development, we performed a number of tests. We used embryos carrying a ubiquitously expressed Lifeact-GFP transgene, which delineates every individual cell in the VE by labelling its cortical F-actin (Riedl et al. 2010). We acquired full z-stack image volumes of E5.5 embryos from two angles (0° and 180°)(Figure S1A) at 5-minute intervals, at diffraction-limited resolution, throughout DVE migration. First, we confirmed the correct patterning of key molecular markers of the three main cell types (DVE, epiblast and ExE) in embryos that had been imaged for 100 time-points at 5 minute intervals. Since each embryo is imaged from two angles, this constituted a total of 200 z-stack acquisitions at 2 μm z-interval. Imaged embryos showed the correct pattern of DVE markers OTX2 and AMOT, epiblast marker OCT4 and ExE marker CDX2, as assessed by whole-mount immunofluorescence and confocal microscopy (Figure S1C). Secondly, time-lapse imaged embryos had intact nuclear morphology and had maintained monolayer integrity, as seen by DAPI and phalloidin staining (Figure S1C). Third, we assessed the extent to which cells failed to divide, which is a hallmark of photo-toxicity (Laissue et al. 2017). Cell divisions could be observed in all tissues throughout the duration of the time-lapse (data not shown) and detailed analysis of all VE cell division events in Lifeact-GFP time-lapse embryos showed a similar rate of divisions among all imaged embryos (Figure S2A). Fourth, to further test if the behaviour observed in live embryos reflected *in utero* development, we imaged embryos expressing the *Hex-GFP* transgene that marks DVE cells and analysed the radial distribution and proximal extent of the Hex-GFP cells throughout the time-lapse. Comparison to non-cultured Hex-GFP control embryos fixed at early-, mid- and late-migration (total N=48) (Figure S2B,C) revealed that the Hex-GFP population in time-lapse imaged embryos progressed through a similar radial distribution and proximal position to that of the non-cultured control embryos (Figure S2D). Finally, we measured the duration of DVE migration in our dataset and found that it was comparable to that of previous studies (Migeotte et al. 2010; Srinivas et al. 2004) (average 4h hours 33 mins ± 1h 24 mins, N=18) (Table S1).

We next used these conditions to build a library of time-lapse image volumes of Lifeact-GFP (Riedl et al. 2010) embryos cultured during DVE migration, that captured the behaviour of every cell in the VE at high temporal and spatial resolution. To automatically stage embryos according to the extent of DVE migration, we developed a support vector machine classifier (see below and Methods) that we applied to time-lapse data from double reporter Hex-GFP:membrane-tdTomato embryos, so that DVE cells are labelled by Hex-GFP (Rodriguez et al. 2001) in a background where all cells express a membrane-targeted tdTomato (Muzumdar et al. 2007). We generated a comprehensive ~12 TB, 5D dataset (3 spatial, 1 temporal, 1 for multiple fluorescence channels) of DVE migration, capturing 7-10 hours of development in each embryo, at intervals of 5 minutes for Lifeact-GFP embryos (N=9), and 10 minutes for double reporter Hex-GFP:membrane-tdTomato embryos (N=9).

### Creation of a dataset of single-cell specific parameters of all VE cells during DVE migration

For the analysis of the multi-embryo dataset, we established a computational pipeline ‘STrEAMS’*(Spatio-Temporal Embryo Analysis at Multiple-Scales)* to automate the analysis of single-cell behaviours and tissue morphology changes throughout DVE cell migration. STrEAMS spatially and temporally registers time-lapse data and enables multi-scale feature extraction (Figure 1 & Figure S3A). For each time-lapse, STrEAMS builds a best quality 3D volume by fusing paired z-stacks from each imaging angle followed by spatiotemporal registration (Movie S1 and S2). Due to the computational complexity of cell segmentation and tracking in native 3D coordinate space, we developed a pipeline to extract the apical VE surface and re-project it as a series of 2D geodesic projections (Movie S3) that could be used for visualisation, augmentation, segmentation and cell tracking before re-projecting to 3D coordinate space for quantifications of cell statistics. In order to make a spatiotemporally consistent 3D to 2D coordinate framework, for each embryo, each time-point of the volume data was 3D shape-matched to a mid-timepoint ‘reference’ volume, while tracking measurements of embryo growth and embryo shape-change.

To enable the visualisation and annotation of cellular events across the entire radial circumference of the apical surface of the VE epithelium, the reference volume was then used to establish a consistent 1- to-1 mapping of selected 3D apical surface coordinates across time and to project the surface onto a flat, 2D geodesic map (Figure 1C). We also manually selected coordinates that corresponded to the basal surface of the VE, to enable local cell height to be measured. Critically, for all quantitative analyses, features annotated on 2D projections were transformed back to their corresponding original 3D coordinates before deriving values for the parameters being analysed (Figure 1C, Movie S6 and S7). This also enabled apical-surface VE measurements to be integrated with local 3D shape and behavioural measurements (see Methods for details).

Visual inspection of geodesic projections of Lifeact-GFP labeled embryos readily revealed DVE cells by their characteristic migratory behaviour. In order to generate binary cell outlines of VE cells, to track them and extract quantitative information about their apical cell-surface morphology and behaviour, we augmented the cortical F-actin signal in our Lifeact-GFP datasets by training a convolutional neural network (CNN) (see Methods and Figure S3E and Movie S4). We selected five of these Lifeact-GFP embryos for further extensive manual curation, enabling us to segment the apical outlines of individual cells across the entire duration of each time-lapse movie (Figure 2E, Movie S4). The centroid of each outlined cell was then tracked frame-to-frame using bipartite matching (see Methods, Figure 2F-F”, Movie S4 & S5). To integrate cell division events, we generated a graphical user interface ‘Cell Tracker’ that enabled the centroids of dividing cells and daughter cells to be easily annotated. The timing, position and division angle of each division was recorded and integrated with cell tracking (see Methods).

**Figure 2.**
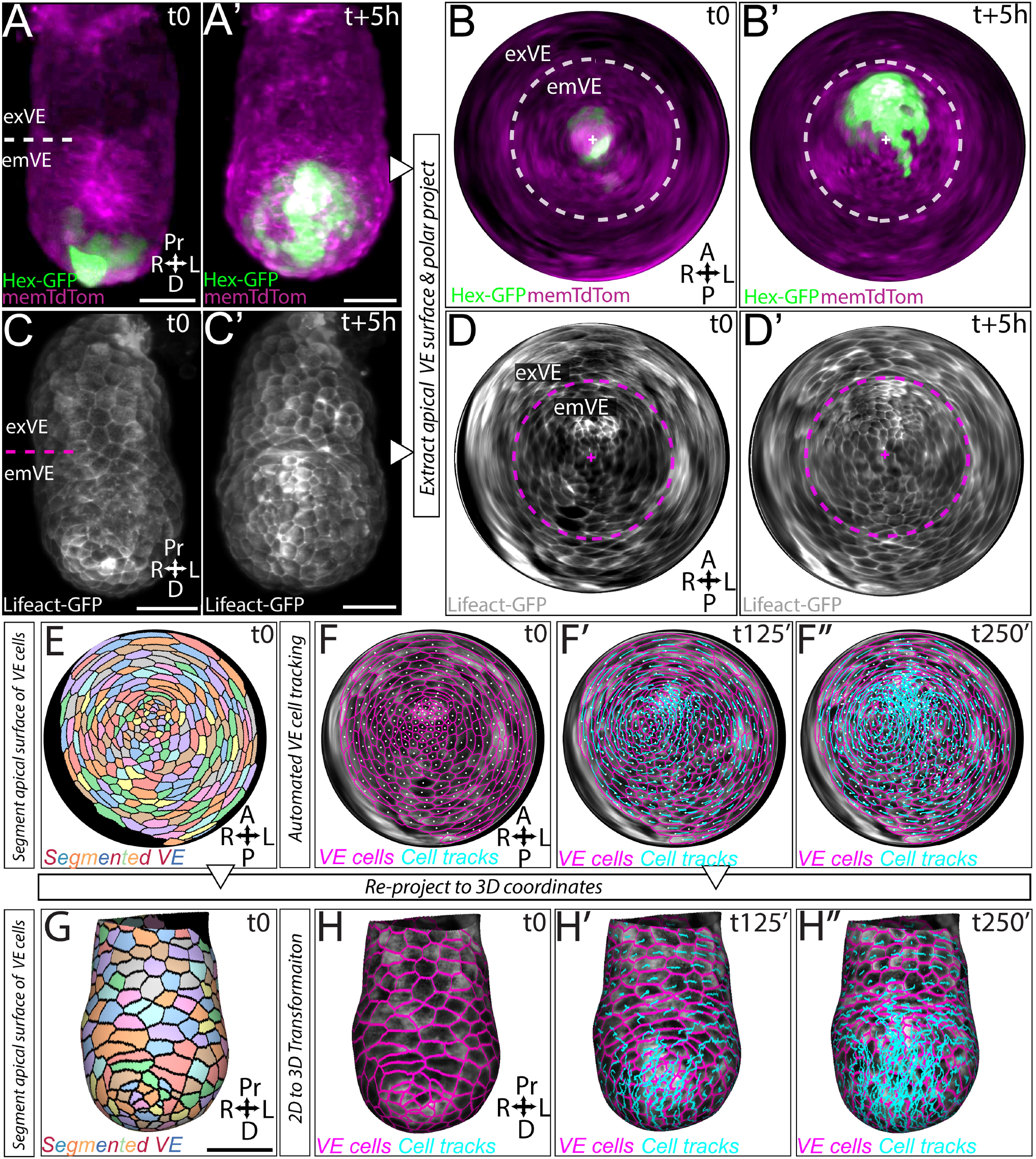
Embryo digitisation and VE cell tracking during DVE migration. **A, A’.** Example of a representative Hex-GFP:membrane-tdTomato embryo at two time-points during lightsheet imaging, used for generating a DVE motion classifier to automate staging of Lifeact-GFP data. **B, B’.** VE apical surface geodesic polar projections of the embryo in A, A’. Distal tip for the embryo indicated by white cross. Dotted line indicates approximate position of emVE - exVE boundary. **C, C’.** representative Lifeact-GFP embryo at two timepoints during lightsheet imaging. **D, D’.** VE apical surface geodesic polar projections of the embryo in C, C’. **E.** Segmentation of VE cells in polar projections after CNN membrane augmentation. **F-F’’.** Three time-points of a polar projection of the VE surface, showing tracks of segmented cells (also see Movie S5). **G**. Reprojection of segmented VE cells into a 3D coordinate space. **H-H’’.** Three time-points from reprojection of segmented VE cells into a 3D coordinate space (also see Movie S7). Scale bar = 50 μm. Note: reprojections have no scale bar as they are non-linear projections.

This generated a comprehensive single-cell longitudinal VE dataset consisting of 91,901 data-points (cell instances), tracking a total of 2221 unique VE cells from five Lifeact-GFP embryos over a cumulative total of 1,900 minutes of time-lapse data. Next, for each VE cell at each time-point, we measured 14 parameters (Table S2) including: two ‘dynamic’ parameters, the instantaneous speed in the anterior direction from the preceding time-point (*anterior speed*) and the cumulative distance a cell has moved in the anterior direction (*cumulative anterior distance*); twelve ‘static’ parameters, relating to the morphology and size of cells at each time-point (*apical cell surface area; apical cell perimeter; shape index; aspect ratio; number of cell neighbours; cell height; mean apical curvature; mean basal curvature; mean apical gaussian curvature; mean basal gaussian curvature; mean Factin levels along cell perimeter* and; *mean F-Actin levels on apical surface*) (see Methods and Table S2). All measurements were transformed to 3D coordinates to calculate corrected values. Together, these measurements describe the instantaneous ‘phenomic state’ of single VE cells (cell behaviour and cell morphology) in the time-lapse data, integrating both dynamic and static information. To visualise the segmented and tracked VE cells in the original context of the egg cylinder, we re-projected our 2D cell outlines (Figure 2F, Movie S4) and tracking back to 3D coordinates (Figure 2H-H”, Movie S7) using Mesh Lab (Cignoni et al. 2008) an open source 3D software interface. This allowed us to distill our multidimensional image data of embryonic development into ‘digital embryo’ representations, that we could interrogate for quantitative insights into DVE migration, and on which we could perform *in silico* labelling ‘experiments’ on cell fate.

Embryos at this stage show some natural variation not only in size but in the duration of DVE migration. To enable integration of data from across multiple embryos, we stage matched embryos in our dataset and spatially aligned them with respect to the major embryonic axes of symmetry. We did this using the motion-based superpixel tracking approach ‘MOSES’ (F. Y. Zhou et al. 2019) to train a support vector machine based DVE classifier. As training data, we used the Hex-GFP channel (that labels migrating DVE cells) from our lightsheet imaged double reporter Hex-GFP:membrane-tdTomato embryo dataset (Methods, Figure S3B). We first spatiotemporally registered and re-projected these time-lapse data as 2D geodesic projections (Movie S8), then used MOSES to extract motion features of the Hex-GFP labelled DVE cells, to train a motion feature classifier (Figure S3B, Movie S9). Finally, we used the trained classifier to categorise the timelapse data for each embryo (Figure S3C, Movie S10) into one of three phases: a first phase with DVE cells at the distal tip of the egg cylinder, with no directional persistence amongst any of the VE cells; a second phase capturing the majority of DVE migration, with consistent directional persistence and; a third phase, with a plateau in directional persistence, during which DVE cells show little or no anterior migration, having reached the embryonic-extraembryonic boundary (Srinivas et al. 2004; Trichas et al. 2011) (Figure S3C).

To understand the cellular basis by which DVE cells migrate directionally from the distal tip to this boundary, we focused on the first two phases, that we termed ‘Pre-Migration’ and ‘Migration’, respectively. To compare cells of corresponding regions from different embryos, we categorised cells according to their location at the start of the migration phase (Figure 3A). To do this, we demarcated eight different anatomical regions of the VE (Figure 3A and Figure S3F) based on four rings of VE spanning the girth of the egg-cylinder along the proximal-distal axis. These were in turn each divided into anterior and posterior halves. Though anterior and posterior become evident only upon DVE migration, the orientation of this axis could be back-propagated even to pre-migration stages because of the longitudinal nature of the data. Prior to migration, the DVE occupied Region ‘1’, the anterior side of the DVE and ‘2’, the posterior side of the DVE (see Methods and Figure S3F for description of how boundaries of these regions were determined). Regions ‘3’ and ‘4’ were proximal VE overlying the epiblast in the anterior (Anterior emVE) and posterior respectively (Posterior emVE). Regions ‘5’, ‘6’, ‘7’ and ‘8’ were the VE overlying the ExE (exVE)(Figure 3A).

**Figure 3.**
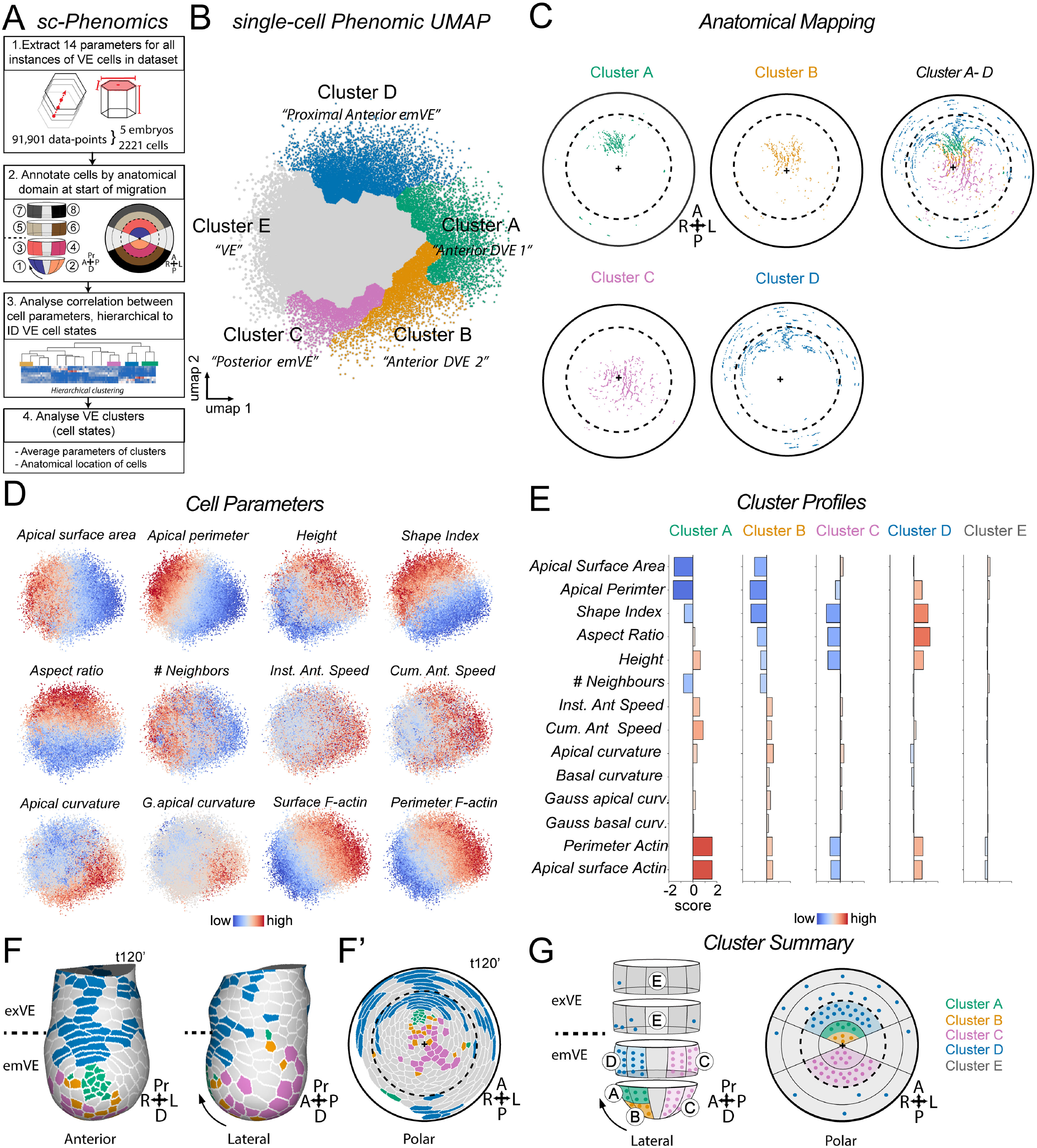
Single-cell phenomic analysis of multi-embryo longitudinal Lifeact-GFP data. **A.** Overview of the steps of the single-cell phenomic analysis method. **B.** 14 parameter single-cell phenomic UMAP integrating all instances (91,901) of all VE cells (2221 unique cells) from five Lifeact-GFP embryos. The five phenomic clusters that emerge are annotated. **C.** Anatomical mapping of all instances of clusters A to D from a representative embryo plotted as a polar projection to visualise both anterior and posterior regions (also see Figure S5 for other embryos). **D.** Single-cell phenomic UMAP with trends of each cell parameter shown as a heatmap. **E.** Mean profile of clusters A-E for each of the 14 cell parameters used in the UMAP analysis. **F, F’.** Single time-point from a representative digitised embryo showing the cluster identity of VE cells in that time point, in 3D and polar-geodesic views (also see Movie S11). **G.** Summary of anatomical origin of VE cells from each cluster. Cluster E was distributed throughout the VE.

Having spatiotemporally registered and defined these relative positional regions in each embryo, we could now integrate information from across multiple embryos to quantitatively investigate the behaviours of the VE during DVE migration, at the level of the component single cells.

### Single-cell phenomic analysis identifies behaviourally and spatially distinct VE cell populations

We analysed our data using two complementary approaches – on the basis of commonalities in cellular phenotype, and on the basis of shared anatomical position. Firstly, to identify in an unbiased manner cells with shared behaviours and morphological characteristics, we performed hierarchical clustering of all VE cells in the dataset based on similarities in the 14 cellular parameters (Methods, Figure S4). In order to incorporate longitudinal changes over time, each instance of a cell in the time-lapse data was included as a separate datapoint. This allowed us, in the first instance, to screen across our data for cells displaying similar behaviours irrespective of temporal or anatomical position, while still retaining this information for later analyses (Figure 3A, Figure S4C). This revealed five distinct phenomic cell clusters that we named A, B, C, D and E. We visualised these clustered data by generating a UMAP that captured the dominant behavioural and morphological phenomic variations over space-time in all VE cells – the phenomic space (Figure 3B). Cells that clustered closely shared a similar set of characteristics while those further apart had more divergent phenotypes. By plotting a heat-map of each parameter on the UMAP it was evident that many parameters were distributed in a graded pattern (Figure 3D). As each cluster was located in a different part of the UMAP, with clusters A-D positioned towards the edge of the UMAP and cluster E making up the remainder (Figure 3B), it suggested that cells in each cluster differ in their phenotype.

As our approach allowed us to retain information about the anatomical location of each cell at every time-point, we asked whether cells in each phenomic cluster were anatomically regionalised. To understand the precise locations of all instances of cells, we first performed a digital anatomical mapping, by extracting the cell IDs of all instances of cells from each phenomic cluster and plotted their position onto the corresponding 2D surface projection (Figure 3C & Figure S5B) and 3D re-projected ‘digital embryos’ (Figure 3F, Movie S11).

Secondly we analysed the anatomical region of origin of cells within each cluster (Figure 3G & Figure S5C’). Together these analyses revealed a striking pattern of anatomical segregation of cells from Clusters A to D. Anatomical digital mapping showed that each cluster was spatially ordered along the proximal-distal and future anterior-posterior axis of the emVE (Figure 3C, Figure S5B). Furthermore, with the exception of cluster E, all other clusters were comprised of a majority of cells that originated in one or two closely located anatomical regions (Figure 3G, Figure S5 B,C’). Clusters A and B consisted predominantly of cells from the anterior half of the DVE (Region 1), Cluster D of cells from anterior proximal emVE (Region 3) and Cluster C of cells from the posterior portion of the DVE and posterior proximal emVE (Regions 2 and 4) (Figure 3C, Figure S5C’). In contrast, Cluster E was comprised of cells distributed uniformly throughout the the emVE and exVE in both anatomical mapping (Figure S5B) and anatomical origin (Figure 3G, Figure S5C’). This indicated that cells that occupy different anatomical regions in the emVE at the onset of migration had characteristic and distinct phenomic signatures. Notably, even Clusters A and B, which consist predominantly of cells from the anterior part of the DVE (Region 1) map, within this relatively small region, to more proximal and distal positions respectively (Figure 3C and Figure 3G). Accordingly, we annotated Cluster A as *Anterior DVE 1*, Cluster B as *Anterior DVE 2*,Cluster C as *Posterior emVE*, Cluster D as *Anterior emVE* and Cluster E simply as *VE*. This extensive phenomic heterogeneity in the VE emerged primarily during DVE migration. When we plotted a UMAP of the subset of data only belonging to the premigration stage (Figure S6B), the vast majority of cell instances fell in the general VE category, with a minority belonging to two DVE clusters (Figure S6 B-D). Having identified this previously unappreciated extensive phenotypic heterogeneity within both DVE and non-DVE cells, we next explored the characteristics that define each of these phenomic clusters.

### Differences in both morphology and behaviour define VE cell clusters

We extracted the average statistics for each of the 14 measurements (Table S3) to calculate the ‘phenomic signature’ for each cluster (Figure 3E). We found that Cluster A (Anterior-DVE 1) cells had low apical surface area, with high values for cell height, anterior-ward speed and levels of F-actin (Figure 3E), as evident by their position on the parameter-highlighted UMAP (Figure 3B, E). Cluster B (Anterior-DVE 2) cells showed similar morphological characteristics, but were not as tall, had lower anterior-ward speed, moved a shorter distance and had a lower level of F-actin in comparison to cells in Cluster A (Figure 3F). Cluster C (posterior emVE) cells had the largest apical surface area and lowest cell height. They are also very regular in shape, showed relatively little anterior-ward motion and had a lower level of apical actin than cells in any other group (Figure 3E). In contrast, Cluster D (‘Anterior Proximal emVE’) cells had intermediate cell area, high cell perimeter, a high aspect ratio, high values for cell height and intermediate levels of F-actin (Figure 3E). Finally, Cluster E contained cells that had intermediate values for each parameter, confirming that these cells reflect ‘average’ VE cells (Figure 3E).

### VE cells undergo coordinated changes in phenomic profile during DVE migration

To understand how the phenotypic characteristics of cells from specific anatomical positions change over the course of DVE migration, we leveraged our longitudinal dataset and digitised representations of embryos. In order to determine the anatomical fate of cells from different parts of the embryo, we tracked them in anatomical space (Figure 4A, B, Figure S7B, Movie S12), and simultaneously, to determine the phenotypic changes they underwent during this process, we tracked them as they traversed phenomic space, to determine their ‘phenomic fate’ (Figure 4A, C, and Figure S7C).

**Figure 4.**
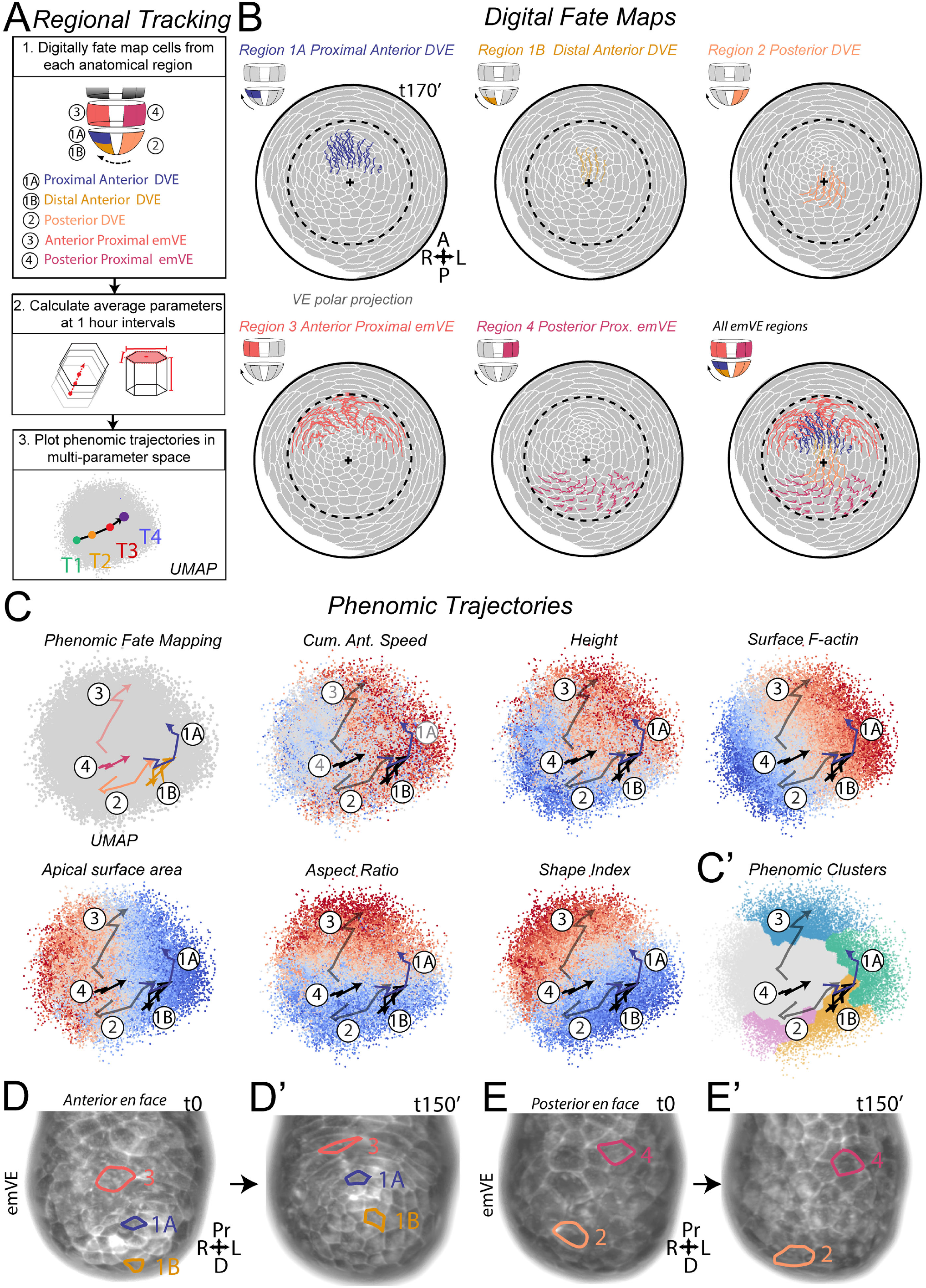
Tracking and phenomic trajectory analysis of emVE sub-regions during DVE migration. **A.** Overview of the steps in the analysis of anatomical sub-regions of the VE. **B.** Continuous tracking of cells from five sub-regions of the emVE from a representative embryo (also see Figure S7 for tracks from other embryos). **C, C’.** Phenomic trajectories of emVE sub-regions overlaid onto UMAPs showing selected cell parameters and the overview phenomic cluster map. **D, D’.** Representative examples of cells from sub-regions 1A, 1B and 3 at two time points. **E. E’.** Representative examples of cells from sub-regions 2 and 4 at two time points.

We used the eight anatomical regions described above (Figure 4A) and in addition, to provide added granularity to the analysis and reflect the heterogeneity revealed by our phenomic cluster analysis (Figure 3G), further sub-divided the anterior DVE into proximal and distal regions giving us a total of nine regions. We then categorised ‘digitally’ labelled cells based on their position at the start of the migration phase, and then tracked them forwards in time during migration as well as backwards in time into the pre-migration phase, to analyse how they change in multi-dimensional phenomic space over time. We generated phenomic trajectories (akin to ‘pseudo-time’ diffusion trajectories in single-cell transcriptomic analyses, but here having the added power of representing real temporal trajectories) by combining data from our stage-matched, spatially aligned embryos, to calculate the average UMAP phenomic coordinates at one-hour intervals for each region. This revealed clear differences in the average phenomic trajectories of cells originating in each anatomical region, with differences in both the direction and length of trajectories (Figure 4C, Figure S7C).

We first considered the anatomical and phenomic fate of DVE cells (subdivided into proximal anterior DVE, distal anterior DVE, and posterior DVE). Anatomical fate mapping showed that cells from all three of these anatomical regions moved unidirectionally towards the anterior (Figure 4B & Movie S11) and were similar across embryos (Figure S7B). However, phenomic fate mapping revealed that cells from each region showed distinct phenomic trajectories; proximal anterior DVE cells (Figure 4C, Region 1A) started in a UMAP region of low apical surface area, low aspect ratio and high apical curvature (Figure 4C, Region 1A). Upon migration, these cells moved towards a region of the UMAP characterised by higher apical F-actin, instantaneous anterior speed and height, but of low apical surface area and aspect ratio (Figure 4C). Cells in the distal anterior DVE (Figure 4A - Region 1B) started in a UMAP region of low apical surface area, low aspect ratio and high apical curvature (Figure 4C). While they increased in apical F-actin and instantaneous speed during migration, in contrast to proximal DVE cells, they did not increase significantly in height (Figure 4C - Region 1B). Furthermore their phenomic trajectory was shorter than that of proximal anterior DVE cells (Figure 4C) indicating that they changed less overall in their phenotype over the course of migration. Despite being migratory, we noted that cells in both of these regions retained low apical surface area and did not appear to significantly change shape prior to their movement (Figure 4C). In contrast, posterior DVE cells had a larger apical surface area at the start of migration (Figure 4C - Region 2), and also had a lower level of F-actin (Figure 4C, Region 2). However, over time they decreased in apical surface area, and increased in anterior speed, F-actin levels and height (Figure 4C) indicating that, over the course of their migration, they start to become more like cells in Region 1A (proximal anterior DVE).

Visualising these trajectories on the UMAP with the cluster boundaries highlighted revealed how tracked cell populations can shift across phenomic clusters (Figure 3C’). When we compared the phenomic trajectories of tracked DVE cell populations to the positions of each cluster, it could be seen that on average, as a cell population, they change cluster over time (Figure 4C). Plotting the trajectory of proximal anterior DVE revealed that its cells begin in cluster E but move through cluster B to cluster A (Figure 4C’ - Region 1A), whereas the distal anterior DVE remained largely within cluster B (Figure 4C’ - Region 1B). This difference in phenomic trajectories of cells in these two, relatively small, regions of the anterior DVE is consistent with them being differentiated into two different phenomic clusters (Figure 3G). Interestingly, the trajectory of posterior DVE cells started in cluster E but moved through cluster C and cluster B (Figure 4C’ - Region 2), consistent with them becoming progressively more like anterior DVE cells over time, as they move into the anatomical region vacated by anterior DVE cells during migration (Figure 4B).

We next considered the cells surrounding the DVE in the emVE (anterior proximal emVE and posterior proximal emVE) (Figure 4A - Regions 3 and 4). Anatomical fate mapping showed distinct behaviours in these regions with anterior proximal emVE showing anterior-lateral movements (Figure 4B - Region 3) while posterior proximal emVE stayed relatively static, with a slight movement towards the distal tip (Figure 4B - Region 4). Phenomic fate mapping revealed that cells in both regions start in a similar central part of the UMAP, i.e., Cluster E, ‘average’ VE cells (Figure 4C). However, over the course of DVE migration, they showed very different trajectories. Anterior proximal cells moved towards cluster D, characterised by higher aspect ratio, high cumulative anterior speed and increased cell height (Figure 4C, Figure 4D-D’). In contrast, posterior proximal emVE remained centrally in the UMAP, largely unchanged in behaviour and morphology (Figure 4C, Figure 4E-E’) showing that these two proximally located regions in the emVE have highly distinct phenomic fates during DVE migration.

Finally we performed fate mapping and phenomic trajectory analysis of exVE anatomical regions (Figure S7A-C). This showed that exVE cells remain relatively static in both anatomical fate mapping (Figure S7B) as well as phenomic trajectories (Figure S7C), in contrast to the dynamic emVE behaviour (Figure 4B, Figure S7B, C). Together, by incorporating data from across multiple embryos, these single-cell phenomic (scPhenomic) analyses revealed the salient temporal behaviour and morphological changes across the entire VE, during DVE migration.

### An unjamming transition occurs not in migrating DVE cells but in the cells ahead of them, the anterior proximal emVE

Our phenomic trajectory analysis showed that despite being highly migratory, anterior DVE (Regions 1A and 1B) cells remained largely unchanged in morphology during migration (Figure 4C-D’). Given that the initiation of movements in an epithelial context is often considered to involve cell shape changes to unjam the migratory tissue, this finding was unexpected. To investigate this in further detail, we took an integrative, tissue-level, approach by pooling phenomic data from across the multiple embryos in our spatially and stage-aligned library of embryos.

To do this, for each of the nine anatomical regions (Figure 5A), we calculated the mean UMAP phenomic coordinate at one-hour intervals by pooling the cells from each region (see Methods). This enabled us to plot the phenomic dynamics for each region over the course of DVE migration (Figure 5A-E, Figure S7D). To visualise these dynamic changes in the context of the embryo, we also overlaid heat-maps for these parameters onto polar projections of the digitised VE cells (Figure 5 B-E, Movie S13).

**Figure 5.**
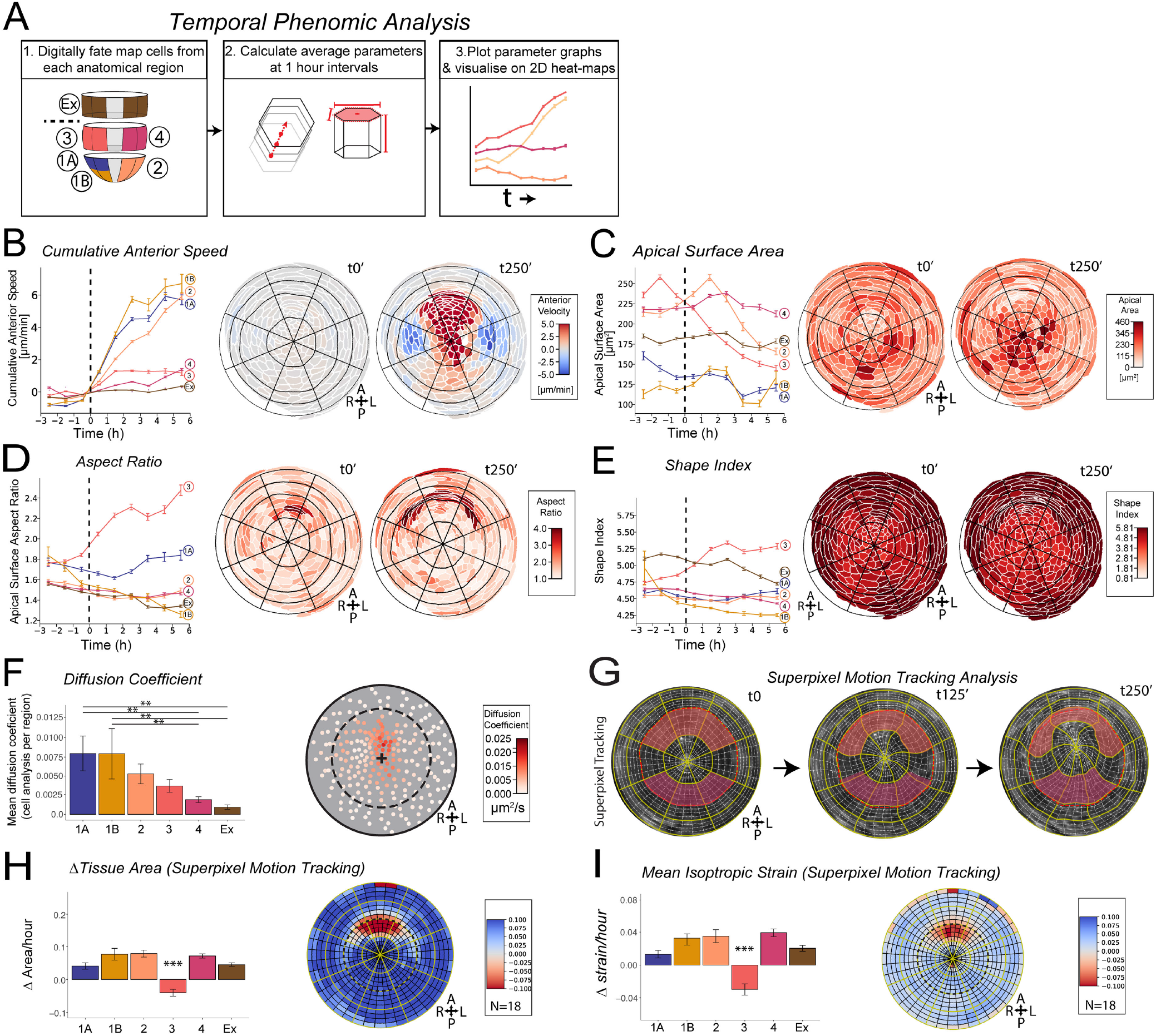
Temporal analysis VE cells in anatomical sub-region. **A.** Overview of the steps for temporal analysis. **B, C, D, E.** Plots of cumulative anterior speed, apical surface area, aspect ratio and shape index (mean±s.e.m) at one hour intervals for cells from each of the five emVE and one exVE sub-regions shown in A. Beside each plot are two time points from a polar projection of a representative embryo with the parameter values overlaid on individual cells (also see Movie S13 for entire animation). **F.** Diffusion coefficient (mean±s.e.m) of cells originating in each VE sub-region, over the course of migration. **G.** Superpixel tracking of tissue behaviour in the polar projection of a representative embryo. A grid is overlaid on the embryo and deformed by motion tracking. Equivalent regions in the anterior (orange) and posterior (magenta) proximal emVE are shown prior to migration (t0) and after motion deformation during migration (t125’, and t250’) showing tissue-level behavioural differences in shape and size. **H.** Fractional area change rate measured by super-pixel analysis. Region 3, the anterior proximal emVE, is the only region to shows a significant difference compared to other regions and is the only region to decrease in surface area during migration. **I.** Mean isotropic change rate measured by super-pixel analysis. Region 3, the anterior proximal emVE, is the only region to show significant difference compare to all other regions, and is the only region to show mean negative isotropic strain.

These pooled data confirmed our scPhenomic analysis, that anterior DVE cells (Regions 1A and 1B), had the highest anterior speed (Figure 5B), smallest apical surface area (Figure 5C) and remained relatively regular in shape throughout migration (Figure 5D, E). To understand how cell movements vary across regions, we next calculated the diffusion coefficient, a measure of the ease of mobility of cells (see Methods). We found that regions 1A, 1B and 2 had significantly elevated diffusion coefficients compared to the other regions, consistent with DVE cells being more mobile (Figure 5F).

Interestingly, this region-anchored analysis highlighted that, corresponding to the onset of migration, cells in region 3 (the anterior proximal VE, comprising cells ahead of the migratory DVE) showed a marked, sustained increase in anisotropy of cell shape (Figure 5D). This suggested that it was cells proximal to the migratory DVE, not cells of the DVE itself, that undergo a phase transition and become unjammed. A hallmark of the jamming-unjamming transition is an increase in cell shape index, a measure of the perimeter vs. areal cortical tension of a cell’s apical surface (Park et al. 2015). While most regions, including the DVE, remained relatively unchanged in their shape index throughout migration, the anterior proximal emVE (Region 3) increased significantly in shape index from the onset of migration and remained high throughout the migration phase (Figure 5E), further supporting the notion that these cells are undergoing an unjamming transition.

The jamming-unjamming transition is a tissue-level event. To complement our single-cell based characterisation of VE cell behaviour with an independent, integrative, characterisation of net tissue-level behaviour, we performed motion-sensing superpixel analysis (Zhou et al., 2019). We performed this on all nine Lifeact-GFP embryos in our time-lapse library as well as our nine Hex-GFP:membrane-tdTomato embryos, as the power of this approach lies in it not requiring the segmentation and tracking of individual cells. We first confirmed that this approach was sensitive enough to track the rotational movement of single-cell tracking, by plotting the average superpixel motion during migration of Lifeact-GFP embryos (Figure S8). Next we seeded a 640-sector grid across the VE tissue by subdividing the area within each of the 32 original regions into smaller 5×4 grids, and allowed the motion of super pixels to deform the finer grid (Figure 5G, Movie S9, MovieS10, Methods). The degree of change over time reflects the cumulative effect of all cellular events including cell movements, cell shape-change, cell division events and cell mixing. We found that the anterior proximal emVE tissue region specifically underwent a deformation at the onset of migration, becoming stretched along the radial axis and compressed along the proximo-distal axis at the Em-Ex boundary (Figure 5G). No other region in the VE showed such a high degree of deformation. Consistent with the single-cell based findings, this analysis verified that Region 3 specifically experienced a net reduction in tissue area (Figure 5H, Movie S14). Importantly, it revealed that this region, uniquely within the VE, is subject to compression, as indicated by a significant negative mean isotropic strain rate (Figure 5I, Movie S14). These tissue-level analyses support the idea that the anterior proximal emVE specifically is undergoing an unjamming phase transition during DVE migration.

### Cell intercalation events are restricted to the region ahead of the DVE

Potential triggers for unjamming include cell divisions (Petridou et al. 2021; Ranft et al. 2010), and cell-cell rearrangements (Merkel and Manning 2017). We therefore next tested whether either of these cellular behaviours differed specifically in the anterior proximal emVE.

We compared the number, timing and angle of divisions (Figure 6A - C’’, Figure S9A - F) within the anterior proximal emVE and the contralateral region of the embryo, the posterior proximal emVE. We found no significant difference in the number of cell divisions events (Figure S9C), consistent with previous studies (Stuckey et al. 2011). We also found no difference in the timing of cells division events in these two regions (Figure S9D,E). Furthermore, there was no burst of division events prior to-, or at the onset of anterior directional DVE migration (Figure S9B). In contrast to the DVE (Regions 1A, 1B and 2) where cell division angles were uniformly distributed (Figure 6C), division angles in the anterior proximal emVE were biased along the radial axis (Figure 6C-C’’) (along the girth of the embryo - see diagram in Figure 6A). However, a similar bias was also found in the posterior emVE (Fig 6C-C’’) suggesting that a difference in the distribution of division angles is not responsible for the specific unjamming in the anterior proximal emVE.

**Figure 6.**
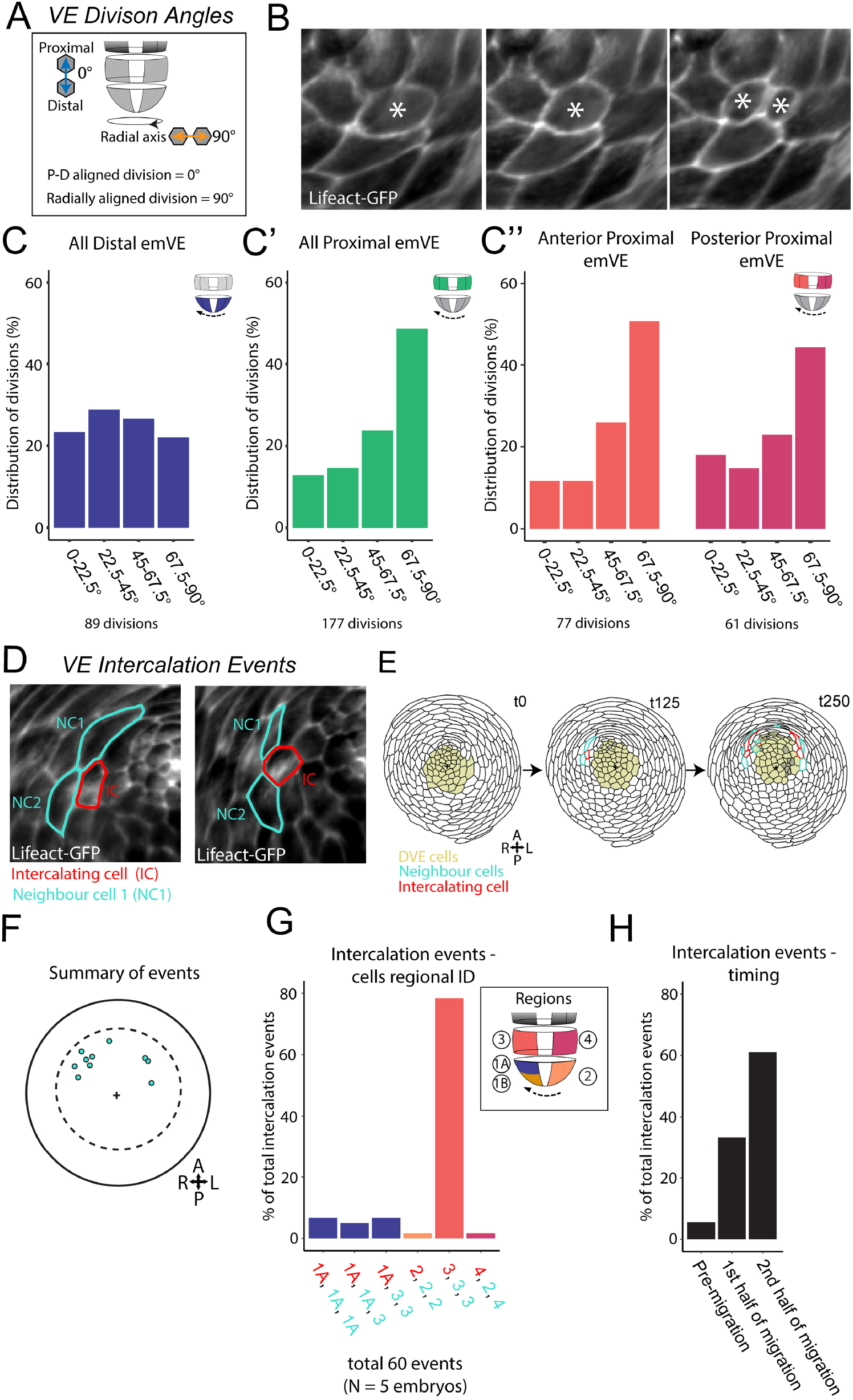
VE cell division angles and cell-cell intercalation events. **A.** Schematic of cell division angle measurement. **B.** Example of a cell division event from a Lifeact-GFP time-lapse dataset. **C.** Cell divisions within the distal emVE binned into four angle ranges. The distribution was not significantly different from random. *χ*^2^ test for expected probabilities, p=>0.05. **C’.** Cell divisions within the proximal emVE binned into four angle ranges. The distribution was significantly different from random. *χ*^2^ test for expected probabilities p=<0.001. **C’’.** Cell divisions within the anterior and posterior proximal emVE, binned into four angle ranges. Both regions showed a distribution that was significantly different from random. Anterior: *χ*^2^ test for expected probabilities p=<0.001, Posterior: *χ*^2^ test for expected probabilities p=<0.005. **D.** Example of VE cell-cell intercalation event in Lifeact-GFP time-lapse data showing two neighbouring cells (NC1 and NC2, cyan) and an intercalating cell (IC, red) moving in between them. **E.** Selected time-points from the polar projection of a representative embryo showing the position of DVE cells (yellow), neighbouring cells (cyan) and intercalating cells (red). See Movie S15 for all frames. **F.** Position of all intercalation events (cyan) across time, in the embryo in E. **G.** Regional distribution of 60 intercalation events from 5 embryos. X-axis show the region of origin of the intercalating cell (red), and neighbouring cells (cyan). The majority of intercalation events (78.3%) occur amongst cells in the anterior-proximal emVE. **H.** Temporal distribution of 60 intercalation events from 5 embryos, across the pre-migration phase (5.5%), first half of the migration phase (33.3%) and the second half of the migration phase (61.1%).

We next analysed our time-lapse data for cell-cell intercalation events, where the contact between a pair of neighbouring cells is broken by a third cell moving in-between them (Figure 6D). We recorded the frequency, timing and anatomical region of intercalation events across the VE in our time-lapse data. In addition to quantifying intercalation events, we also visualised them on the polar projections of digitised embryos (Figure 6E, Movie S15). This revealed a striking localisation within the VE of cell intercalation events, that occur almost exclusively in the anterior proximal emVE (Figure 6F, G). Intercalation events also increased over the course of migration, with the majority occurring in the latter half of the migration phase (Figure 6H). Cell intercalation events within the migrating DVE (1A, 1B and 2) were exceedingly rare (Figure 6G). As cell mixing is a hallmark of unjamming, this further supports a model where the anterior DVE remain jammed during migration while anterior proximal emVE cells undergo unjamming.

### DVE cells have higher apical membrane tension than the surrounding emVE

The primary characteristic of jammed tissues is mechanical constraint (Lawson-Keister and Manning 2021; Mitchel et al. 2020). Our single-cell analysis showed that migratory DVE cells had higher levels of F-actin compared to the surrounding VE, suggesting they may be under distinct mechanical stresses (Figure 3F - Cluster A & B, Figure 4C). To confirm that F-actin levels are indeed higher in DVE than surrounding cells, we calculated the relative levels of F-actin in each cell, on its apical surface and along junctions with surrounding cells. We did this using the normalised intensity of the Lifeact-GFP signal at one-hour intervals prior to and during migration. This revealed that even prior to migration, DVE cells show the highest level of F-actin (both junctional and apical surface) and maintain this difference throughout migration (Figure 7A, B), indicating that they might be mechanically distinct from surrounding VE cells.

**Figure 7.**
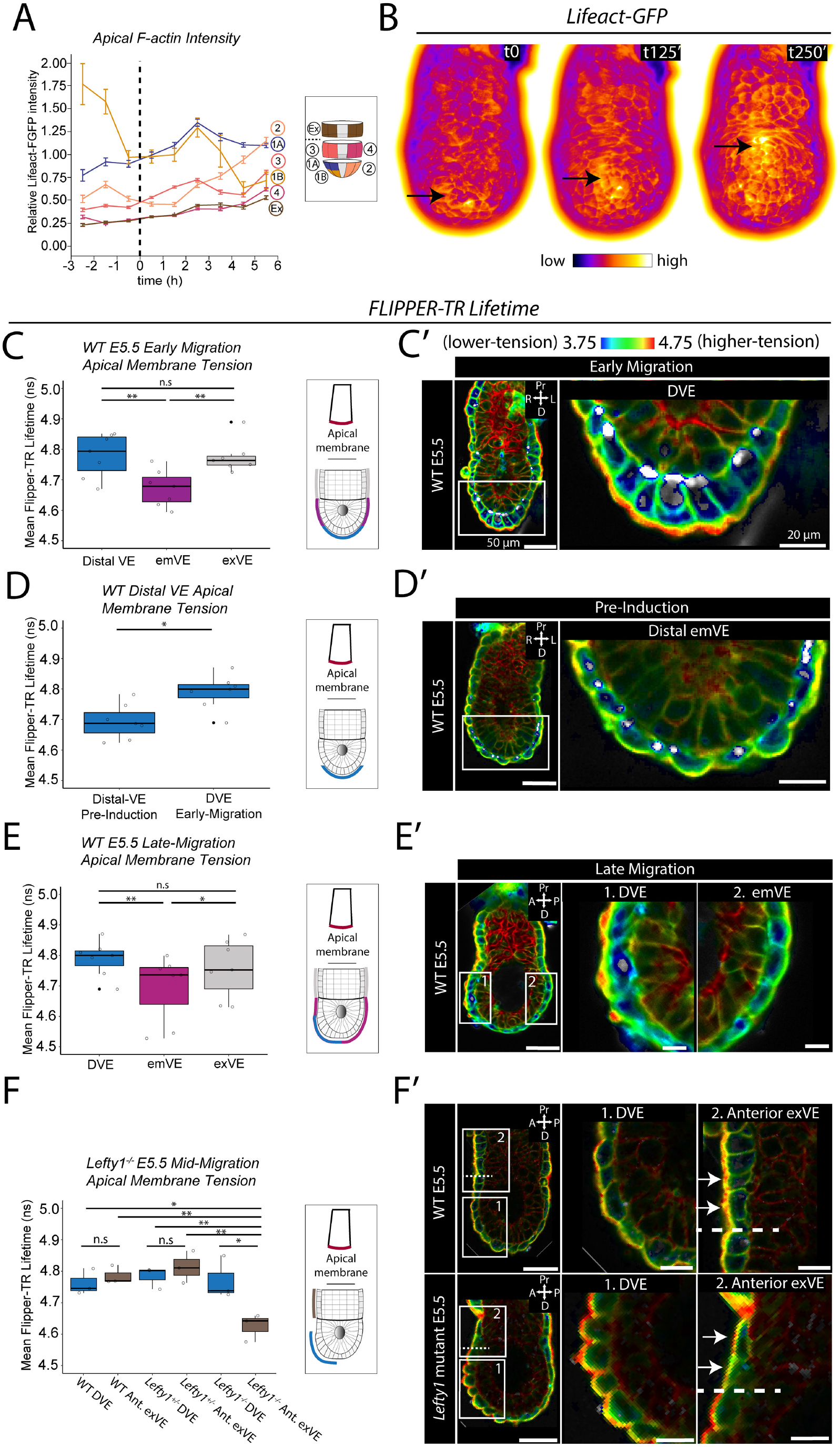
Migratory DVE cells show elevated apical F-actin and higher membrane tension. **A.** Plot of relative apical Lifeact-GFP intensity (mean±s.e.m) at one hour intervals for cells in each of the five emVE and one exVE sub-regions shown in the inset. **B.** Extended focus surface projection of Lifeact-GFP intensity from multiple time points of a representative embryo. The arrow shows the position of the same DVE cell at each time point. **C, D, E, F.** Fluorescence lifetime imaging microscopy (FLIM) and quantitation of FLIPPER–TR membrane tension reporter in mid-sagittal optical sections. **C.** Apical membrane lifetime of DVE cells, emVE cells not belonging to the DVE, and exVE in early migration embryos (N=7). Apical lifetime was significantly higher in DVE and exVE compared to emVE, reflecting a higher tension in the former two regions (one-way ANOVA, p=<0.01, followed by Tukey’s HSD Test on DVE vs. emVE (p =<0.01) and exVE vs. emVE (p=<0.05)). There was no significant difference between DVE and exVE (Tukey’s HSD Test: p=>0.05). **D.** Apical membrane lifetime of cells at the distal tip of E5.5 embryos prior to (N=7) and after (N=7) they had acquired DVE identity. Apical lifetimes were significantly higher in DVE cells, compared to cells from pre-induction embryos (Student’s t-test, p=<0.05). **E.** Apical membrane lifetime of late-migration DVE cells, emVE and exVE from E5.5 embryos (N=7). Lifetimes were significantly higher in migrated DVE than in the remaining emVE (one-way ANOVA, p=<0.001, followed by Tukey’s HSD Test, p =< 0.01) and in exVE compared with emVE (Tukey’s HSD Test, p=<0.01). **F.** Apical membrane lifetime of mid-migration *Lefty1*^−/−^ null (N=3), *Lefty1*^+/−^ heterozygous (N=3) and wild type (N=3) embryos. There were significant differences in tension based on genotype and region of the embryo, one-way ANOVA, p=<0.001, followed by Tukey’s HSD Test for specific comparisons. There was no difference in lifetime between anterior exVE and migrated DVE in WT or *Lefty1*+^/−^ heterozygous embryos (Tukey’s HSD Test, p=>0.05 for both comparisons). However in *Lefty1*^−/−^ mutant embryos, anterior exVE had a significantly lower lifetime than the DVE (Tukey’s HSD Test, p=<0.01). Furthermore, the anterior exVE from *Lefty1*^−/−^ mutant embryos had significantly reduced tension when compared with the DVE and anterior exVE of wild type and *Lefty1*+^/−^ heterozygous embryos (Tukey’s HSD Test on *Lefty1*^−/−^ anterior exVE vs. wild type DVE (p=<0.05), wild type anterior exVE (p=<0.05), *Lefty1*^+/−^ heterozygous DVE (p=<0.01) and *Lefty1*^+/−^ heterozygous anterior exVE (p=<0.001)). **F’.** In inset images; dotted line marks the emVE - exVE boundary and arrows highlight cells in the anterior exVE region.

To test this prediction, we made use of a live-cell tension-sensitive dye, Flipper-TR, whose fluorescence lifetime is a readout of tension (Colom et al. 2018) and has previously been used to characterise mechanical tension in E6.5 mouse embryos (Royer et al. 2022). We labelled live wild-type E5.5 embryos and used the columnar morphology of DVE cells (Kimura et al. 2000; Rivera-Perez et al. 2003; Srinivas et al. 2004) to identify the position of the DVE and categorise embryos as ‘early-migration’ stage based on whether the DVE was at, or just off-set from, the distal tip (Figure 7C and Figure S10A). We then performed fluorescence lifetime imaging microscopy (FLIM) to capture a mid-sagittal z-section of each embryo, so as to compare the apical membrane tension of DVE cells to that of cells in the surrounding emVE and exVE. This revealed that apical tension in DVE cells was significantly higher than that in surrounding emVE, but not significantly different from exVE (N=7) (Figure 7C-C’).

The elevated tension of the DVE could be a property intrinsic to DVE cells, or could be the result of their being positioned at the distal end of the egg cylinder, which is the site of highest curvature. To distinguish between these two possibilities, we examined embryos prior to DVE induction (Figure 7D-D’) a stage at which VE cells positioned at the distal tip of the egg cylinder have not yet acquired the molecular (expression of genetic markers) (Belo et al. 1997; Pfister et al. 2007; Thowfeequ et al. 2021; Yamamoto et al. 2004), or morphological hallmarks (columnar morphology) of the DVE (Kimura et al. 2000; Rivera-Perez et al. 2003; Srinivas et al. 2004). We found no significant difference in apical tension between the pre-DVE cells at the distal tip and the surrounding cells in E5.5 pre-induction embryos (Figure S10B). Furthermore, the apical tension was significantly lower in the cells at the distal tip of preinduction embryos than in DVE cells of E5.5 early-migration embryos (Figure 7D), suggesting that elevated membrane tension is acquired by distal cells only upon induction to the DVE state and is not simply a property of cells by virtue of their position at the distal tip of the egg cylinder.

To further test this finding, we examined late-migration embryos in which DVE cells had moved away from the distal tip and towards the Em-Ex boundary (Figure 7E-E’). We found that DVE cells retained elevated apical tension compared to surrounding emVE cells even after they had migrated away from the distal tip of the egg cylinder (N=7) (Figure 7E-E’). These data further indicate that DVE cells remain mechanically distinct from surrounding emVE cells even during migration, consistent with them remaining in a jammed state, as a solid flock.

Given the high levels of F-actin in DVE cells, we tested whether elevated apical tension in DVE cells might be the result of increased actomyosin based contractility, using the Myosin inhibitor blebbistatin (Straight et al. 2003). While control DMSO treated embryos showed no change in tension after treatment (N=9) (Figure S10C-C’), blebbistatin treated embryos showed a significant and rapid decrease in tension (N=9) (Figure S10C-C’). This points to an actomyosin dependent mechanism for the elevated apical tension of DVE cells.

### Lefty1 dependent modulation of mechanical heterogeneity in the VE delimits DVE migration

DVE cells do not normally migrate into the exVE region, despite it being part of the contiguous monolayer epithelium of the VE. Instead, DVE cells abruptly stop migrating proximally at the exVE and start to be displaced laterally (Srinivas et al. 2004; Takaoka et al. 2011; Trichas et al. 2011). In both E5.5 early-migration and E5.5 late-migration embryos, we observed that while the proximal emVE cells had a significantly lower tension relative to the DVE, exVE cells had a high tension, comparable to that of the DVE (Figure 7C, E). We hypothesised that this matched high-tension may prevent DVE migration into the exVE. To test this hypothesis, we measured VE cell tension in *Lefty1* null mutants (Meno et al. 1998) (Figure 7F-F’, and Figure S10D-D’) in which DVE cells abnormally over-migrate into the exVE (Trichas et al. 2011). While E5.5 mid-migration WT (N=3) and heterozygous *Lefty1*^−/+^ embryos (N=3) showed no difference in tension between DVE and exVE (Figure 7F), mid-migration *Lefty1*^−/−^ null mutants (N=3) had significantly lower anterior exVE tension compared to DVE cells (Figure 7F). This was due to a significantly reduced tension in the anterior ExVE of mutants in comparison to WT and heterozygous embryo, while the tension of mutant DVE cells was unaffected (Figure 7F). Furthermore this lower tension was found only in the anterior exVE of mutants embryos, but not their posterior exVE (Figure S10D’). This lowering in apical membrane tension in the anterior exVE could help explain the over migration phenotype observed in *Lefty1* null embryos.

## DISCUSSION

### DVE cells remain as a solid flock during collective migration

DVE migration takes place in an intact monolayer epithelium requiring extensive coordination of cell behaviour, so as to retain tissue integrity while enabling dynamic movements. By generating a multi-embryo, single-cell resolution longitudinal dataset through lightsheet microscopy, and developing a computational pipeline to extract and analyse fundamental VE cell morphological and behavioural parameters, we have been able to decompose global tissue-scale events to their component cellular behaviours. We find that, in contrast to other epithelial tissues where the onset of cell movement often involves a transition of cells to a fluidised state, the DVE remains in a jammed state throughout migration. It is the non-migratory VE cells ahead of the DVE that show hallmarks of an unjamming transition (cell shape changes and mixing), leading to fluidisation of the tissue. Migrating DVE cells in contrast show hallmarks of the jammed state (Lawson-Keister and Manning 2021) such as crowding and elevated tension with little change in the shape or size of cells. By remaining in a jammed state, DVE cells are bound together as a collective, enabling them to migrate via basally located projections (Migeotte et al. 2010; Srinivas et al. 2004), as a coherent group displacing the anterior emVE cells ahead of them.

This finding suggests that DVE cells migrate as a solid flock, and offers the first *in vivo* example of this. It supports the more nuanced way of considering the role of the unjamming transition in the context of epithelial migration, where unjamming need not be an obligate characteristic of migrating cells, but can be a property of cells being deformed by an actively migrating collective of cells moving as a solid flock (Lawson-Keister and Manning 2021). In the case of the VE, the solid flock of DVE cells might even be responsible for triggering the unjamming liquefaction of the cells ahead of them, by changing the directional stress they are subject to (Cates et al. 1998; Liu and Nagel 1998). One characteristic of solid flocks is that they are ‘self-propelled’. Migrating DVE cells show hallmarks of active migration, such as cell projections polarised in the direction of motion (Srinivas et al. 2004), and being dependent on the molecular regulators of active cell migration NAP1 (Rakeman and Anderson 2006) and RAC1 (Migeotte et al. 2010).

Flocking behaviour can be described as either ‘solid’ or ‘liquid’ depending on the extent of local cell mixing superimposed on the long-range correlated movement of the flock (Giavazzi et al. 2018; Trepat and Sahai 2018). Given that DVE cells have an important signalling role in pattering the epiblast, migration as a solid flock might be necessary for it to remain as a coherent cell population so that the signals emanating from it can be properly localised, rather than being diluted or misplaced by dispersal of DVE cells were they to migrate through unjamming or as a liquid flock. Interestingly, embryos mutant for Nap1 (Rakeman and Anderson 2006), or in which PCP signalling is disrupted (Trichas et al. 2011), show an abnormal dispersal of AVE cells within the visceral endoderm similar to what a migrating liquid flock might be expected to resemble, pointing to potential molecular modulators of flocking properties of DVE cells. Flocking behaviour can be induced in confluent human mammary epithelial cells by over expression of Rab5a, a regulator of endocytosis that is thought to promote junctional remodelling in this context (Malinverno et al. 2017). It is unknown what happens in the VE if one perturbs Rab5 function, but embryos genetically null for another endocytic regulator, Rab7, show patterning defects consistent with defective DVE induction or migration (Kawamura et al. 2012). Though the precise cause of the defect in these mutants is unclear, this presents the possibility that endocytosis might be modulating junctional remodelling in the the mouse embryo as well.

We had previously reported that neighbour exchange was restricted to the emVE and not seen in the exVE but due to limitations in imaging, could not tell whether neighbour exchange occurred in the posterior VE (Trichas et al. 2011). Lightsheet image volumes and our digitised embryos allow us to examine the entire surface of the egg cylinder and reveal the striking extent to which neighbour exchange events are restricted to only the anterior emVE, supporting the idea that it is this region specifically that is undergoing unjamming.

Studies of cell migration in other models generally consider relatively flat expanses of tissue composed of many cells. The difference we observe in the mouse VE might be because it is only a distinct subset of the VE, the DVE, that shows migratory movement. It might also be due to the cylindrical arrangement of the VE, with a relatively small circumference and high curvature. The range of shape index values we recorded for VE cells differed from the critical shape index of 3.81 reported for jamming in 2D epithelia (Park et al. 2015), presumably due to the intrinsic 3D curvature of the VE. This highlights the potential influence of overall tissue topology on the properties and collective behaviour of component cells. It will be interesting in future studies to examine other cylindrical epithelia such as the ureteric branches of the developing kidney or pulmonary branches of the developing lung to determine to what extent behaviours in cylindrical epithelia differ from those in relatively flat epithelia.

### Regional heterogeneity of behaviour and mechanical state circumscribes DVE migration

DVE cells become more columnar than neighbouring VE cells upon induction and remain that way throughout their movement (Kimura et al. 2000; Rivera-Perez et al. 2003; Srinivas et al. 2004). Here we show that distal VE cells prior to induction are squamous and have a similar membrane tension to the surrounding emVE. When they are induced to form columnar DVE cells, apical tension increases. If columnarity is linked to the cells occupying a different mechanical regime, it might explain why DVE cells first undergo this change, perhaps to increase their contact area, allowing them to adhere more effectively to each other and facilitating their migration as a collective through the surrounding VE cells. We also note from other studies that DVE cells express elevated levels of specific cytoskeletal modulators, Keratin19 and Drebrin molecules (Thowfeequ et al. 2021), and Keratin8 (Despin-Guitard et al. 2022) that could provide additional mechanical differences.

The significantly higher tension of DVE cells relative to surrounding emVE cells suggests a model whereby the high-tension collective of DVE cells migrate through a ‘permissive’ lower tension emVE. DVE does not normally migrate into the exVE despite this being part of the continuous monolayer tissue of the VE. We find that in wild-type embryos, the emVE shows a similar tension to that of the DVE, both higher than that of the emVE. These findings could explain why DVE migration results in a rotational, flow-like pattern in the emVE as shown by us and others (Shioi et al. 2017; Takaoka et al. 2011; Trichas et al. 2011), that may emerge as a consequence of the jammed DVE population deforming the unjammed emVE against the relatively high-tension exVE, that defines the limits of DVE migration. Interestingly, these tissue-wide movements appear similar to the ‘polonaise’ movements in the pre-gastrulation chick epiblast (Cui et al. 2005; Graper 1929; Voiculescu et al. 2007; Wetzel 1929) which has also been shown to be surrounded by a high-tension boundary (Saadaoui et al. 2020), though in this case, the cellular mechanisms (mediolateral intercalation events, oriented cell divisions)(Firmino et al. 2016; Voiculescu et al. 2007) appear to be different.

In *Lefty1* null mutants, the DVE over-migrates into the exVE region (Trichas et al. 2011). Our finding that in these mutants, the tension is lower in anterior exVE but not the DVE, suggests that in the wild-type embryo, exVE is a barrier to migration because its mechanical properties are matched to those of the DVE and that a differential in tension is required for migration. This opens up the interesting possibility that patterned differentials in tension across epithelia might be a general mechanism through which cell movements is controlled in other epithelial contexts.

As *Lefty1* is expressed in DVE cells, how its loss leads to reduced tension in the exVE is unclear, though as a secreted ligand, it could be expected to have non-cell-autonomous effects on nearby tissues. LEFTY1 is an inhibitor of the TGF-β family member NODAL, that has recently been shown to mediate an unjamming transition during zebrafish gastrulation (Pinheiro et al. 2022). In this system, a gradient of NODAL leads to unjamming mediated by changes in cell motility (Pinheiro et al. 2022). Though the cellular mechanism of unjamming in the mouse egg cylinder appears to be independent of cell motility, it is possible that it is still ultimately mediated by LEFTY1 acting via NODAL signalling. *Nodal* is expressed in the DVE (Brennan et al. 2001; Thowfeequ et al. 2021) and its downstream effectors, SMAD2/3 are nuclear localised in VE cells (Yamamoto et al. 2004; Yamamoto et al. 2009), indicative of active NODAL signalling. Furthermore, we have previously shown that *Nodal* and *Lefty1* mutant embryos mislocalise the Planar Cell Polarity protein DVL2 (Trichas et al. 2011), suggesting a possible mechanism by which LEFTY1 might modulate the behaviour and mechanical properties of the VE.

### Molecular regulation of patterned cell behaviours in the Visceral Endoderm

What are the molecular players regulating regionalised behaviours in the VE? Our recent transcriptomic analysis of the E5.5 VE (Thowfeequ et al. 2021) has identified two distinct transcriptional sub-clusters within the DVE that spatially correspond roughly to the phenomic clusters A and B (Anterior DVE1 and 2 respectively), allowing one to postulate a transcriptional underpinning to the phenomic grouping. Our transcriptomic characterisation also identified a novel role for Ephrin and Semaphorin signalling in the VE (Thowfeequ et al. 2021), that might be involved in establishing the behavioural and mechanical differences we observe between the DVE, surrounding emVE and exVE.

We also note that the anatomically restricted cell-cell intercalation events that we observed in the anterior proximal emVE corresponds to a region that expresses DKK1 (Kimura et al. 2001; Thowfeequ et al. 2021). In addition to acting as an inhibitor of the canonical WNT pathway, DKK1 has been shown in zebrafish to also interact in the WNT-PCP pathway (Caneparo et al. 2007), which can regulate cell-cell rearrangements in other developmental contexts (Voiculescu et al. 2007; Wallingford et al. 2000; Yang and Mlodzik 2015). It is possible that DKK1 might be facilitating the neighbour exchange and cell shape changes that lead to the unjamming of anterior proximal emVE cells.

### Single-cell phenomic analysis to identify localised behaviours and morphologies

Analyses of lightsheet time-lapse datasets during development have predominantly focused on cells tracked using nuclear markers, or involved the analysis of pre-selected sub-groups of cells, or the averaging of behaviour over time. By using lightsheet microscopy to image entire volumes of Lifeact-GFP expressing embryos, we were able to generate data on the cell outlines of all the VE cells of the cylindrical embryo. In order to leverage our rich, longitudinal dataset, we developed a machine-learning based approach to segment cells and a quantitative methodology to unbiasedly study the changing phenotypic characteristic of cells – a single cell ‘phenomics’ approach. Using this novel methodology, we were able to integrate data from across multiple embryos and leverage the large numbers of cells and time points to identify distinct morphological and behavioural sub-clusters which are spatially ordered in the embryo, leading to the insights on DVE migration presented here. This approach is applicable to a variety of contexts and we anticipate that it will be a valuable method for other researchers generating and analysing similarly high temporally and spatially resolved data. Similarly, our extensive dataset of curated longitudinal cell phenotypes represents a unique resource to colleagues for developing and testing theoretical models.

## ACKNOWLEDGEMENTS

We thank Dr Antonio Scialdone, Dr Ben Simons and Dr Berta Verd for valuable discussions and helpful comments on the manuscript. We thank Dr Satish Arcot Jayaram for help with mouse breeding, Dr Tristan Rodriguez for the Hex-GFP line, Dr Roland Wedlich-Söldner for the Lifeact-GFP line and Dr Liqun Luo for the ROSA26^mTmG^ line. This work was funded by BBSRC Research Grant BB/J00989X/1, BBSRC ALERT13 Award BB/L014750/1, Wellcome Strategic Award 108438/Z/15/Z and Senior Investigator Award 103788/Z/14/Z to SS. FZ and XL were funded by the Ludwig Institute for Cancer Research. HH was funded by Wellcome Trust Chromosome and Developmental Biology Doctoral Training Programme 109100/Z/15/Z. JT and SY were supported by EPSRC grant EP/T10001569/1 via the Alan Turing Institute and EP/V001310/1. FS and MF would like to acknowledge generous funding from the Rosalind Franklin Institute and the Kennedy Trust for Rheumatology Research, as well as Wellcome (212343/Z/18/Z) and EPSRC (EP/ S004459/1). FS is grateful for the support by EMBO (ALTF 849-2020) and HFSP (LT000404/2021-L) fellowships. We thank the Micron Advanced Bioimaging Facility, supported by Wellcome Strategic Awards 091911/B/10/Z and 107457/Z/15/Z, and the Wolfson Imaging Centre for microscopy support. We thank Pathology Services Building and Biomedical Services staff for excellent animal support.

## METHODS

### Mouse Strains, Husbandry, and Embryo Collection

Genetically modified mice were maintained on a mixed C57Bl/6 CBA/J background. The Hex-GFP line (Rodriguez et al. 2001) was bred into the ROSA26^mTmG^ (Muzumdar et al. 2007) background to create a double homozygous line. Hex-GFP:membrane-tdTomato stud males were crossed with CD1 females (Charles River) for live imaging experiments. The Lifeact-GFP line (Riedl et al. 2010) was maintained as a heterozygous line and crossed with CD1 females (Charles River) for live imaging experiments. For FLIM experiments C57Bl/6 studs were crossed with CD1 females for all wild-type analysis. *Lefty1* (Meno et al. 1998) mice were crossed for FLIM experiments and mutants identified post-imaging though PCR genotyping as previously reported (Trichas et al. 2011). All mice were maintained on a 12 hour light, 12 hour dark cycle. Noon on the day of finding a vaginal plug was designated 0.5 days *post coitum*. Embryonic day 5.5 (E5.5) embryos were dissected in M2 medium (Sigma) with fine forceps and tungsten needles and transferred into pre-heated culture medium (as per (Trichas et al. 2011)) supplemented with antibiotics, and placed in an incubator at 37°C, 5% CO2 prior to imaging.

### Embryo mounting for lightsheet imaging

Low melting point agarose (2%) (Sigma) in 1 × phosphate buffered saline was drawn up into a 20 μl glass capillary (Brand, 701904) using a teflon coated plunger (transferpettor piston rod) (Brand, 701934). A lumen was created within the agarose cylinder using a 150 μm diameter copper wire. Once solidified, the wire was removed, the end of the cylinder was sealed with agarose and a window to the lumen was cut with a razor blade. Two embryos, with their proximal ends opposing one another were transferred into the lumen of the agarose cylinder in a 35 mm petri-dish filled with culture medium. The agarose supporting the embryos was withdrawn into the glass capillary, the capillary was then transferred to the imaging chamber of a ZEISS Z1 lightsheet microscope. Once positioned in the imaging chamber, the agarose supporting the embryos was extruded proud of the glass capillary prior to imaging.

### Lightsheet time-lapse imaging

Embryos were imaged in a ZEISS Z.1 lightsheet microscope using a plan-apochromat 63X/1.0 NA water immersion lens. Full volume z-stacks 1920 × 1920 pixels, 16bit resolution of each embryo were obtained at a 2 μm interval, recorded sequentially from 2 imaging angles (0° and 180°), with a 135 μW ± 228 nW 488 nm (Coherent) or 106.9 μW ± 248 nW 561 nm laser (Coherent) 106.9 μW ± 248 nW for eGFP and tdTomato, respectively. For Hex-GFP:membrane-tdTomato, eGFP and tdTomato channels were obtained in parallel using a 561 LP secondary beam splitter. Each z-plane was illuminated sequentially with right and left illumination pivot scans oscillating at 23 kHz. The laser lightsheet was focused through a pair of 10X/0.2 NA lenses and the paired lateral illuminations were fused using ZEN Black (ZEISS) “online dual-fusion” setting. For Lifeact-GFP embryos a pair of z-stack volumes were acquired every 5 minutes. For Hex–GFP:membrane-tdTomato experiments a pair of 2-channel z-stack volumes were acquired every 10 minutes. All equipment was sterilised prior to each experiment by autoclaving or UV and Ozone treatment in a cool-CLAVE steriliser (AMSBIO).

### Immunofluorescence

Embryos were fixed in 4% PFA at room temperature for 20 min, washed at room temperature three times for 5 min each in 0.1% Triton-X100 in PBS; incubated in 0.25% Triton-X100 in PBS for 25 min; washed three times in 0.1% Tween-20 in 1xPBS; blocked with 2.5% donkey serum, 2.5% goat serum, and 3% Bovine Serum Albumin (BSA) in 0.1% Triton-X100 in PBS overnight; then incubated overnight at 4°C in primary antibodies diluted in 1:100 in blocking solution. Embryos were washed three times in 0.1% Tween-20 in PBS (PBT) for 5 min each, with a final additional wash for 15 min; incubated overnight at 4°C with appropriate secondary antibody 1:100 in 0.1% PBT; embryos were incubated with phalloidin at 1nM concentration in PBT overnight at 4°C, washed four times for 5 min in PBT at room temperature; and finally mounted with Vectashield mounting media containing 4’,6-diamidino-2-phenylindole (DAPI) (Vector Labs H-1200).

### Antibodies and Phalloidin

Primary antibodies used were 1:100 goat anti-AMOT (Santa Cruz, 82491), 1:100 rabbit anti-CDX2 (Cell Signalling, 9775), 1:100 rabbit OCT-4 (Abcam, ab200834), 1:100 rabbit anti-OTX2 (Cell Signalling, 11943S). Secondary antibodies used were 1:100 Alex-Fluor (AF)-555 donkey anti-rabbit (Invitrogen, A31570), 1:100 AlexFluor (AF)-633 donkey anti-goat (Invitrogen, A21082). For F-actin staining phalloidin-atto 647N (Sigma, 65906) or Alexa Fluor 488 phalloidin (Invitrogen, 49409) were used at a 1nM final concentration in PBT.

### Confocal Microscopy of fixed embryos

Fixed embryos were imaged on a ZEISS LSM 880 confocal microscope using a 40x oil (1.36NA) objective. Z-stacks of embryos were acquired at 1 μm interval using non-saturating parameters. 3D opacity rendering images were made using Velocity Software (Improvision). Figures were prepared with Adobe Photoshop and Adobe Illustrator (Adobe Inc.).

### FLIM experiments

Fluorescence lifetime imaging (FLIM) experiments were carried out as previously reported (Royer et al. 2022). For wild-type C57Bl/6 (Charles River) studs were mated with CD1 (Charles River) females to generate E5.5 embryos that were dissected in phenol red-free M2. Embryos were then transferred to 8-well imaging plates (No. 1.5 glass. ThermoFisher Scientific) and incubated at 37°C in 250 μls of 1 μM FLIPPER-TR probe (Spirochrome, SC020) membrane tension reporter (Colom et al. 2018), and diluted in phenol-free M2. The 8-well imaging chambers were mounted on the pre-heated stage of an Leica SP8 with a Fast Lifetime Contrast (FALCON) module allowing for acquisitions at high photon counts using LAS-X (Leica Microsystems) software for acquisition and pre-processing. Embryos were imaged at 37°C using a 20× water immersion objective (Leica C PL APO CS2 20×/0.75 IMM). FLIPPER-TR was excited at 488 nm with a tuned white light laser (WLL: NKT Photonics) pulsing at 20 MHz. Zoom was at 2.2 × yielding an 264.4 um2 field of view covered by 1024 × 1024 pixels. The pinhole was set to 1.2 AU, scan speed 200 Hz and 25 repeats were acquired. Fluorescence was collected from 499 - 701 nm on an HyD-SMD detector (Leica Microsystems). Pixels were binned by a factor 4 to increase signal to noise and confidence in photon arrival times. Pixels containing only background photons (less than 50 counts) were removed. Lifetime images were generated using the Phasor-FLIM workflow in LAS-X. We used the phasor plots and a rainbowfalse-colouring from 3.75 - 4.75 nm to aid in visualisation of membrane tension differences. For quantification, Phasor-FLIM images were exported to .tiff (using 0.01 lifetimes per grey level) and were further processed using the macro previously developed (Royer et al. 2022). ROI’s were drawn in FIJI using the intensity tiff image to unambiguously identify the apical membrane of each VE cell. The ROI was then applied to the lifetime channel alone and the lifetime vales per pixel within each region saved to a .csv file. Embryos were staged through morphological assessment of the position of columnar DVE cells, and the apical membrane lifetime vales from multiple embryos from each stage were combined for further analysis in R to calculate mean lifetimes of the apical membrane of VE cells using the tension-sensitive lifetime range of 2.8 - 7 ns (Colom et al. 2018). Lefty1 (Meno et al. 1998) FLIM experiments embryo were carried out as above. Post-imaging, all embryos were recovered and processed for PCR genotyping. For pharmaceutical inhibitor culture FLIM experiments, embryos were prepared as above in 8-well imaging plates incubated in 1 μM FLIPPER-TR probe and a baseline FLIM acquisition of a single mid-sagittal optical section captured of each embryo. Each well was then further incubated with either 2 μM blebbistatin (Sigma), or 1:1000 DMSO as a vector control, for 30 minutes before a second FLIM acquisition was taken. For analysis a total of 9 embryos incubated with DMSO, and 9 with blebbistatin, were analysed and the baseline compared with the 30 minutes treatment across each group.

### STrEAMS (Spatio-Temporal Embryo Analysis at Multiple-Scales) data processing framework

Figure S3 illustrates the key steps to process raw .czi (Carl Zeiss Imaging Format) two-angle, z-stack time-lapse data of live E5.5 embryos imaged using a ZEISS Z.1 lightsheet microscope. 3D z-stack time-lapse data were processed to obtain a 2D “unwrapped” view of the apical surface of the visceral endoderm (VE) for quantitative analysis. As the visceral endoderm epithelium remains as a monolayer throughout DVE migration (Trichas et al. 2011), projection from 3D-to-2D significantly simplifies the segmentation and tracking of all VE cells. By retaining each cells’ original coordinate information, calculation of 3D corrected values is enabled, allowing for the efficient integration of cell behaviour with cell morphology analysis. Processing can be grouped into three distinct modules; pre-processing, spatiotemporal registration and 3D-to-2D VE apical surface projection (“unwrapping”). The input to the framework is a volumetric time-lapse of an embryo from two angles. The output is the unwrapping coordinate mapping from 2D-to-3D and the unwrapped 2D time-lapse of the apical surface of the VE monolayer.

### Data preprocessing

The raw image data, stored in the ZEISS CZI image file format, of two angle (0° and 180°) z-stack volume acquisitions and all time-points from the ZEISS Z.1 lightsheet microscope were processed to make voxels spatially isotropic by image interpolation and to make the full volumetric time-lapse of an embryo computationally tractable for registration on a single PC; by 16bit to 8bit conversion, volumetric downsizing by a factor of 2 and cropping out empty voxels using a bounding box. The resulting volumes are saved as .tif per angle, per time-point and has an isotropic xyz voxel resolution of 0.363 μm; a size of ~200MB per 8bit .tif file, 30-45GB for the whole time-lapse with two acquisition angles per time-point.

### Intensity-based spatiotemporal registration

We spatiotemporally register both angles and all time-points in a multi-step procedure to obtain a single fused volumetric time-lapse capturing only cell movement independent from motion artefacts and embryo growth. Importantly all transformations are reversible, enabling accurate single-cell quantification data to be calculated, reincorporating growth and 3D shape information.

#### Rigid registration

Using time-point 1 (t1) as the reference volume, we temporally registered all volumes captured from angle 1 in sequential manner using Matte’s mutual information (Mattes et al. 2001; Raghunathan et al. 2005) and similarity transformation; permitting only translation, rotation and scaling (Fig S3A, step i). The typical default parameters for similarity registration were (750, 750, 100) iterations at downsampled scales of [16,8,4] using Matlab’s imregtform. Translation compensates for the drift of the embryo and rotation for global rotations of the embryo around and towards the embryonic long axis. Scaling removes the components of cell movement due to embryo growth over the imaged duration. The rigid registered volumes isolate only the components of relative cell movement caused by cell migration and cell shape change.

#### Angle alignment

To combine angle 1 with angle 2, we rotate angle 2 by the known angle (180°) and refine the alignment by translation-only registration (and Matte’s mutual information metric) to the matched pre-temporally registered angle 1 as reference (Figure S3A, step ii). The typical default parameters for translation registration were (1250, 1250, 100) iterations at downsampled scales of [16,8,4] using Matlab’s imregtform. This step reasonably assumes that any embryo rotation and growth is negligible in the ~ 30s between the imaging angles (20s to image the volume, 10s to reset). The angle 1 temporal transforms are applied to the angle 1 spatially aligned angle 2 volumes to obtain an equivalent temporally registered angle 2 time-lapse at all time-points for fusion (Figure S3A, step iii).

#### Tilt correction

To enable coherent 2D surface projections of 3D volume data, the long (proximal-distal) axis of each embryo was manually aligned using morphological landmarks (ectoplacental cone, columnar DVE cells, epiblast tissue) to correct for any tilt in their mounting. Due to the prior temporal registration, the tilt correction angle had only to be found for t1 and could then applied to all other time-points to obtain spatiotemporally registered and proximo-distally aligned volumes for both angles.

#### Angle fusion

The two-angle acquisitions at each time-point were fused using a custom sigmoidal blending scheme, with a specified depth parameter marking roughly half way through the embryo. The sigmoidal blending allows a smooth transition of pixel intensity between the two angles at the fusion depth parameter. The depth parameter was manually specified per embryo due to the asymmetric exposure mentioned above.

#### Non-rigid registration

We lastly use non-rigid registration (Demons (Thirion 1998; Vercauteren et al. 2009), Matlab imregdemons) to match the outer embryo shape over time to the shape at a reference time-point half-way through the timelapse. This step is essential to compensate for embryo shape variations and create a static reference volume for surface unwrapping and tracking single cells. To ensure warping of the outer shape with minimal impact to individual cells, multi-scale non-rigid registration was applied to downsampled volumetric images at three different resolutions, 4×, 8× and 16× such that the embryo-level detail of the outer edge of the embryo and the inner cavity were apparent, but not the cellular detail. The regularisation parameter in the optimisation objective of Demons was manually determined per embryo to find a balance between visualisation of cell shapes and the embryonic surface.

### 3D-to-2D VE apical surface unwrapping

The 3D apical VE surface was re-projected (‘unwrapped’) to 2D to enable the visualisation of the entire circumferential surface of each embryo in a single flat image for subsequent analysis. This process encompasses three steps; semi-automatic binary embryo segmentation to generate the VE surface to unwrap as a collection of (*x, y, z*) surface coordinates; specifying the 3D-to-2D unwrapping transformation and projecting the 3D image to 2D image intensities by interpolation. Due to the spatial-temporal registration (above) the unwrapping transform needs to only be generated for t1, then iterated across all subsequent time-points in the time-lapse.

#### Semi-automatic binary embryo segmentation

An initial binary segmentation capturing the VE surface was automatically found either by binary Otsu thresholding or by using a fixed intensity threshold followed by a series of opening and closing 3D morphological operations. The automatic result was then manually refined slice-by-slice per embryo, from proximal to distal. The corrected binary was downsampled, smoothed, and re-upsampled to generate the final surface. The concatenation of all slice-by-slice contour points along the proximal-distal axis generates the (*x, y, z*) surface coordinates for re-projection (unwrapping).

#### Find 3D-to-2D unwrap coordinates

In order to unwrap, we need to parameterise; resample and find a new set of (*x,y, z*) surface points where each (*x, y, z*) also has a unique (*s, θ*) coordinate thereby generating unique 5-tuples, (*x,y, z, s, θ*). This is guaranteed when we choose *s* to be the geodesic distance of (*x, y, z*) along the curved embryo surface relative to a fixed specified reference coordinate, (*x*_0_, *y*_0_, *z*_0_) (here the distal most surface point along the long embryo axis through the embryo’s centroid), and *θ* ∈ [0,2*π*] is the radial rotation angle. We developed a two-step approach to generate (*x, y, z, s, θ*) from the input surface coordinates from the previous binary segmentation which we denote (*x*^0^, *y*^0^, *z*^0^) in this section. The first step generates (*x*′, *y*′, *z*′ *s*′) by binning the data into angle bins and applying spline fitting, smoothing and resampling in each bin. The second step generates the desired (*x*″, *y*″, *z*″, *s*″, *θ*″) from the first step, binning the points into *s* bins and applying spline fitting, smoothing and resampling in each bin. In particular, the resampling is a crucial step to ensure the embryo surface has been uniformly sampled. We let ′ and ″ denote the output from the first and second step respectively.

##### Step 1

*From* (*x*^0^, *y*^0^, *z*^0^) *to* (*x*′, *y*′, *z*′, *s*′). For input (*x*^0^, *y*^0^, *z*^0^) surface coordinates from binary segmentation, compute the radial angle, *θ*^0^ = tan^−1^((*z*^0^ – *z*_0_)/(*y*^0^ – *y*_0_)) relative to the fixed distal reference coordinate,(*x*_0_, *y*_0_, *z*_0_) and let *x* denote the proximal-distal axis. This is appended to each coordinate to produce 4-tuples, (*x*^0^, *y*^0^, *z*^0^, *θ*^0^). Discretising the angle space, *θ* ∈ 0-2*π* radians into 480 bins we assign each (*x*^0^, *y*^0^, *z*^0^) surface points into one of the bins according to *θ*^0^. In each angle bin, the coordinates represent unordered sampling of a line segment cross-section of the embryo surface. To obtain a uniform sampling and to measure exactly the geodesic distance we use splines; fitting an independent spline for *x*-, *y*-, *z*-coordinate. To fit the spline, we must first order the (*x*^0^, *y*^0^, *z*^0^) in the bin according to their geodesic distance from the reference (*x*_0_, *y*_0_, *z*_0_). For embryos, which are smooth and pseudo-cylindrical and therefore largely convex, we use the Euclidean chordal distance as a good surrogate to the unknown geodesic distance that preserves the relative distance ordering from (*x*_0_, *y*_0_, *z*_0_). A cubic spline was then used to parameterise and fit a smooth line through each of the sorted *x*-, *y*-, *z* - coordinate with a function, *f* of a single variable *t* with value from 0-1 such that (*x*^0^,*y*^0^, *z*^0^) =(*f_x_*(*t*), *f_y_*(*t*), *f_z_*(*t*)), *t* ∈ [0,1]. The variable *t* corresponds to a normalised geodesic distance, that is when we evaluate *t* in 1000 equal increments, the output (*x*′, *y*′, *z*′) = (*f_x_*(*t*), *f_y_*(*t*), *f_z_*(*t*)), *t* = 0,0.001,0.002,…, 1 coordinates are ordered, lie on the embryo surface cross-section and the distance between them are the same. We can compute *s*’ for each generated (*x*′, *y*′, *z*′) as the cumulative sum of all pairwise differences preceding the point in the order. For example, *s*’ of the 5^th^ point is the sum of the distances of the first point to (*x*_0_, *y*_0_, *z*_0_), the second to the first, the third to the second, the fourth to the third and the fifth to the fourth. Applying the described spline parameterisation to points in every angle bin, resampling 1000 points per bin gives the full set of (*x*′, *y*′, *z*′, *s*′) coordinates.

##### Step 2

*From* (*x*′, *y*′, *z*′, *s*′) *to* (*x*″, *y*″ *z*″, *s*″, *θ*″). Step 2 smooths the obtained coordinates angularly before recomputing the associated angle, *θ*. This helps reduce discontinuous artefacts that may arise from processing the embryo with independent 1d splines in step 1. Similar to step 1, we discretise the range of the geodesic distance, *s*′ from 0 to 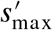, the maximum distance into bins of increments 1 voxel i.e. 0,1,2,3…., *s*’_max_ where [·] is the ceiling function. Similarly we assign each (*x*′, *y*′, *z*′, *s*′) point into one of the bins according to *s*′. These points represent a concentric cross-section of the embryo. For each bin, we thus sort the points angularly according to computed *θ* = tan^−1^((*z*′ –*z*′_0_) /(*y*′ – *y*Ȳ_0_)), where (*x*′_0_, *y*′_0_, *z*′_0_) is the centroid of the coordinates in the bin. We then fit a periodic smooth cubic spline for each *x*′, *y*′, *z*′, *s*′ to obtain individual functions, *f* of a single variable *t* with value from 0-1 such that (*x*′, *y*′, *z*′, *s*′) = (*f_x_*(*t*), *f_y_*(*t*), *f_z_*(*t*), *f_s_*(*t*)), *t* ∈ [0,1]. Again, the variable *t* corresponds to a normalised angular distance, such that the cumulative angular differences is the cross-section circumference and we can resample 1000 equi-angularly spaced points with refined coordinates, (*x*′, *y*′, *z*′, *s*′) per bin. Doing so for all bins we finally produce the final set of 5-tuple (*x*″,*y*″, *z*″, *s*″, *θ*″) with *θ*′ = tan^−1^((*z*″ – *z*_0_) /(*y*″ – *y*_0_)) computed relative to the fixed distal reference coordinate, (*x*_0_, *y*_0_, *z*_0_).

#### Project image 3D-to-2D

To obtain a 2D unwrapped projection image of the embryo surface we populate an *M* × *N* × 2 image grid of desired (*s*, *θ*) coordinates and use *k*-nearest neighbours (*k* = 1) trained on (*x*″, *y*″, *z*″, *s*″, *θ*″) to get the matching *M* × *N* × 3 image grid of (*x*, *y*, *z*) coordinates. This is often called pullback. Trilinear interpolation of the volumetric pixel intensity at the *M* × *N* × 3 (*x*, *y*, *z*) coordinates for every time-point of the spatiotemporally registered embryo produces the final unwrapped 2D time-lapse videos for each embryo. We used two different *M* × *N* × 2 (*s*, *θ*) coordinate grids to compute two different projections; the Cartesian and Polar geodesic projections.

##### Cartesian geodesic projection

This pseudo-cylindrical projection creates the *M* × *N* × 2 grid with the geodesic distance *s* as the row coordinate and radial angle *θ* the column coordinate. *s* equally samples 0 to [*s*′_max_]+1 where [·] is the ceiling function i.e. we let *M* = [*s*″_max_]+1. *θ* equally samples 0 to 0-2*π* radians with *N* equal bins. *N* is chosen so that the ratio *M/N* preserves *s*″_max_/*c*″_max_ where *c*″_max_ is the maximum cross-section circumference of a *s*-bin from Step 2 above.

##### Polar geodesic projection

This projection creates the *M* × *M* × 2 grid (i.e. *N* = *M*) such that the geodesic distance *s* and the radial angle *θ* are the polar coordinate of the image grid i.e. if (*i, j*) denotes the image row and column coordinate respectively then: (*s, θ*) = (*i* – *M*/2)^2^ + (*j* – *N*/2)^2^, tan^−1^((*i* – *M*/2) /(*j* – *N*/2))). We choose *M* to be [1.2 [*s*″_max_] / 2 ] where [·] is the ceiling and [·] the floor functions respectively. This means that the radial distance of *i, j* from the image centre, (*M*/2,*N*/2) equally samples 0 to 1.2 [*s*″_max_] with *M*/2 bins for *θ* = 0,*π*/2,*π*, 3*π*/2 radians.

3D-to-2D unwrapping inevitably distorts the true Cartesian surface distances. By construction the Cartesian geodesic projection best preserves distances of the cylindrical embryo surface whilst the Polar geodesic projection best preserves distances around the distal tip where DVE cells originate and migrate from. Note since there is bijection between the *M* × *N* × 2 image grid of (*s, θ*) and the *M* × *N* × 3 grid of (*x, y, z*) we can always remap any processing in the subsequent analyses done in 2D back into Cartesian 3D to obtain true geometric measurements.

### Tissue motion extraction for alignment and staging

To extract the tissue motion prior to cell segmentation we used motion sensing superpixels (F. Y. Zhou et al. 2019). Instead of using the Farneback optical flow (Farnebäck 2003) as per the original publication, we used DeepFlow (Weinzaepfel et al. 2013) that captures the collective tissue dynamics and produced motion fields with close agreement to manually annotated single cell tracking (below). We extracted superpixel motion tracks at 1000 superpixels (larger region-of-interest) and 5000 superpixels (smaller region-of-interest) on both 2D Cartesian and polar geodesic projection time-lapses of each embryo. We used the tracks from 1000 superpixels to train a classifier to identify DVE-associated superpixel tracks for migration staging and tracks from 5000 superpixels to compute DVE migration stage and the DVE migration angle.

### Automated DVE migration staging

As the duration of DVE migration varies between individual embryos, staging the dataset was necessary to allow consistent comparison between embryos. To enable objective staging, we developed an automatic pipeline using the motion characteristics of migrating cells from MOSES extracted superpixel tracks (1000 superpixels). The pipeline comprises two modules; the first module involves training a classifier using a novel time-lapse dataset of lightsheet imaged Hex-GFP:membrane-tdTomato embryos in which DVE cells are labeled by Hex-GFP to classify each superpixel track of a ubiquitously Lifeact-GFP labelled embryo as “DVE” or “non-DVE” motion associated. The second module computes the difference in cumulative persistent distance moved between the mean DVE track vs the mean non-DVE track to classify each time-point of an embryo time-lapse into one of 3 phases: no directional persistence, consistent directional persistence (reflecting the movement of Hex-GFP labeled DVE cells from distal tip to Em–Ex boundary), or a plateau in persistence (after the Hex-GFP labeled cells have reached the boundary).

#### Manual staging of embryos

To train a motion-based DVE classifier, two researchers independently staged 2D surface projections of registered time-lapse data from 9 Hex–GFP:membrane-tdTomato embryos into 3 stages (as above) by following the behaviour of the Hex-GFP labeled DVE cells. The annotation of both researchers were combined to produce a single consensus staging for each embryo, the middle frames were used as consensus when not precisely matched. Lifeact-GFP datasets were also manually staged as a baseline to compare with automated results.

#### Module 1, DVE-associated track classifier

Hex-GFP:membrane-tdTomato (where Hex–GFP expression labels DVE cells) time-lapse datasets were processed through the pipeline to 2D surface projections and used to guide the training of a classifier to assign each MOSES extracted superpixel track computed over the manually annotated DVE migration phase as either DVE (*Hex*+ve) or non-DVE (*Hex-ve*) -associated (Figure S3). The trained classifier could then operate on MOSES superpixel tracks of 2D surface projections of Lifeact-GFP embryos extracted over the entire video. We use the Cartesian geodesic projection and found that a 1000 superpixel coverage worked best as higher superpixels produced a non-continuous classification (data not shown). As *Hex* exhibits salt and pepper expression and not all cells in the DVE region express *Hex-GFP* equally (Srinivas *et al.*, 2004; Migeotte *et al.*, 2010; Takaoka, Yamamoto and Hamada, 2011), we applied Otsu binary thresholding to the maximum projection *Hex-GFP* image (computed over time) to generate the binary superpixel class for supervised training. In total 9 *Hex-GFP* embryos were used to train the classifier. As the Hex-GFP:membrane-tdTomato embryos are imaged at 10 minute intervals compared to 5 minutes for Lifeact-GFP embryos we encoded each superpixel track as a velocity feature vector of length 18 by concatenating the mean velocity of the superpixel track of interest (2 values, one each for x- and y-directions, red node, Figure S3B) and the mean velocity of the surrounding 8 superpixel tracks (black nodes, Figure S3B). The classifier takes the 18-vector of velocities as input to predict a binary variable where DVE=1 and non-DVE=0. Inclusion of neighbouring superpixel velocities helped promote greater spatial continuity in the final classification. As the number of embryos was small, we used a support vector machine (SVM)(Boser et al. 1992) classifier which is conservative and performs well on small datasets due to its maximum-margin property equipped with an RBF kernel to handle nonlinearity. To ensure further robustness, we trained a separate RBF-SVM classifier for *k*=*10* random 5/4 partitionings of the 9 *Hex-GFP* embryos (i.e. 5 embryos for training, 4 embryos for testing). Each embryo further comprises 2 Cartesian geodesic projections unwrapped with respect to reference radial angle *θ* = 0 and *θ* = *π* radians respectively. This can be seen as a data augmentation to promote robustness to the unknown DVE migration angle. This gives a total 18 trained classifiers. We use all trained classifiers to produce an ensemble prediction more robust to any individual training partition. Specifically a superpixel is only classified as DVE=1 if at least 0.85 × 18 > 15 or 18 classifiers predicted the superpixel was DVE=1. The largest graph connected component of the DVE=1 superpixels yields the final DVE classification.

#### Module 2, Staging DVE migration based on the persistence of migration

We developed an unsupervised method to use the trained DVE classifier on the Cartesian geodesic projection to stage individual Lifeact-GFP embryos from MOSES superpixels tracks extracted in the Polar geodesic projection. The polar projection best captures in a single spatiotemporally continuous manner the initiation of DVE migration at the distal tip and its migration to the Em-Ex boundary (5000 superpixels). We first apply the Cartesian trained DVE classifier to MOSES superpixel tracks extracted from only the manually specified migration to boundary stage of Lifeact-GFP embryos, where available, or the MOSES superpixel tracks extracted over the full duration, if not available. The classified superpixels designate the region of the image that comprise DVE cells at the time-point (TP) at the start frame of (manually annotated) migration, *TP*_mig_ if classifying using migration stage or the first frame, *TP*_0_ if using the full duration. We stress the use of *TP*_mig_ is not necessary for computing the staging. We use it here to show how to incorporate human guidance in the DVE classification. As any motion prior to migration, was minimal we found there was almost no deviation between the final results. To transfer the Cartesian projection classification result into the polar projection, we remapped the DVE classified superpixel points. The mapping is learnt by nearest neighbour matching between the respective unwrapping coordinates. We then compute the concave hull of the polar-mapped DVE points to get a spatially contiguous binary mask. The binary mask is applied to the MOSES superpixel tracks (5000 superpixels) extracted directly from the polar projection over the full video duration by direct lookup at time-point *TP*_mig_ or *TP*_0_ respectively to classify tracks into DVE and non-DVE. If the (*i, j*) row, column coordinate position of superpixel track at *TP*_mig_ lies inside the binary then the whole superpixel track is designated DVE, else it is non-DVE. DVE migration is characterised by continual persistent directional motion. We use this *prior* knowledge to stage. As we do not know the persistent angle of migration and this can differ per track, we define a cumulative persistence measure, *d_direct_*, the cumulative directional distance moved for a track *SPi* at time *t* using the mean velocity as the persistent direction, 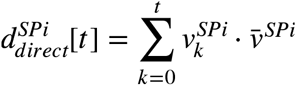 where *ν* is the displacement between time *t* and *t* + 1, 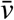 the unit normalised mean velocity, representing the persistent direction and · the dot product. We evaluate *d_direct_* after mapping back to Cartesian 3D. We compute the instantaneous 3D velocity 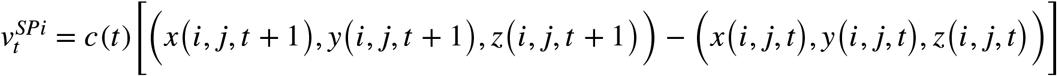 using a linear piecewise approximation that assumes 3D displacement is small and where *c*(*t*) is the constant scaling correction factor from the rigid temporal registration at time-point *t*. We use (*i, j, t*) as shorthand to denote the (*i, j*) row, column coordinate position of the track at time *t*. The unit normalised mean velocity vector is determined from the mean of the 2D superpixel velocities in the unwrapped polar projection. In 3D on a curved surface, the 3D equivalent directional vector will change depending on the surface position. Denoting the fixed 2D vector as 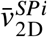, we compute the 3D version at a position (*x*(*i, j, t*), *y*(*i, j, t*), *z(i, j, t*)) at time *t* using the linear piecewise approximation above with the next time-point a 2D displacement of *k* pixels from (*i, j, t*) in the direction of 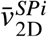; 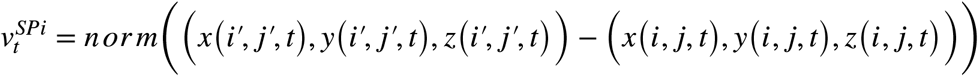 where 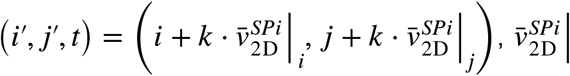 and 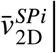 denote the *i*-, *j*-direction component of 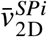, and *norm* denotes unit vector normalisation such that magnitude of the 3D vector is 1. The displacement *k* pixels is used to avoid getting the zero 3D vector. *k* should therefore be chosen small. We empirically find *k* = 10 pixels is good. We compute directional persistence for all superpixel tracks, and average within the DVE and non-DVE tracks separately to obtain two separate curves, 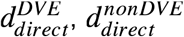 respectively. We subtract 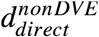 from 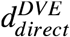 to normalise 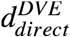. We further scale normalise to obtain a dimensionless measure by dividing 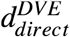 by a characteristic length, *l*, the cube root of the embryo volume at time-point 0. We don’t compute the real embryo volume since we have bounding box cropped during spatiotemporal registration. Instead we use the spatiotemporally registered volume image dimensions of *X* × *Y* × *Z* pixels so that 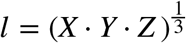. We then stage DVE migration by identifying distinct stepwise changes in the temporal rate of change, 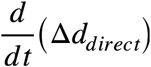 in the normalised 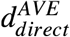 denoted Δ*d_direct_*. Stepwise changes are detected by computing a breakpoint score [0-1], 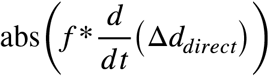 given by the absolute magnitude of the ‘same’ padding convolution of a step function, 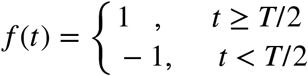 where *T* is the number of frames with 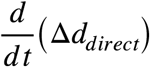, and detecting the time-points of the local peaks of height > 0.15 and separated by at least 5 frames (25 min). Any valid time-points must further be in the time interval [3,*T* – 3] frames. All detected valid time-points is sorted chronologically, [*t*_1_, *t*_2_,…, *t_N_*]. We construct all contiguous time intervals formed by the time-points, [0, *t*_1_], [*t*_1_, + *t*_2_, … [*t_N_* + 1, *T*] and compute for each interval the mean Δ*d_direct_*. We then stage the intervals chronologically based on the observation Δ*d_direct_* should be large and non-zero in migration to boundary and close to 0 in pre-migration and post-migration with these distinguished by whether they occur before or after migration. The observation is hard-coded as computational rules as follows. We first detect if there exists a time interval exhibiting a migration phase, defined as having a mean Δ*d_direct_* > a fixed threshold of 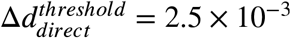 for all embryos. If multiple time intervals satisfy this criteria, the interval with the highest Δ*d_direct_* is designated as the migration anchor interval, *anchor_mig_*. If an interval is found, we set all intervals starting at times before *anchor_mig_* and have mean 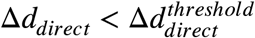 as ‘pre-migration’. We then go through the remainder unclassified time intervals in temporal order, comparing if their mean Δ*d_direct_* is closer to the mean Δ*d_direct_* of all ‘pre-migration’ or the ‘migration’-classified intervals. If an interval is closer to ‘pre-migration’ and precedes the earliest interval of ‘migration’ interval, then it is designated ‘pre-migration’. If the interval occurs after the latest ‘migration’ intervals, then it is ‘post-migration’ and all remainder intervals are ‘post-migration’.

The final algorithm automatically stages a given embryo video into three behavioural/motion phases: Phase I: no directional persistence, Phase II: consistent directional persistence, Phase III: a plateau in directional persistence.

### A-P Axis Alignment (DVE migration-angle determination)

We determine the DVE migration angle to enable alignment of each embryo along their future anterior-posterior axis and enable consistent mapping (sub-regionalisation) across embryos. Angle determination proceeds from migration staging (above) where the DVE classifier has been applied to classify polar extracted MOSES superpixel tracks (5000 superpixels) as; DVE or non-DVE. We then find the most directionally persistent subset within the DVE-classified tracks to compute mean direction as the DVE migration angle. We compute the migration angle from both 2D unwrapped polar and Cartesian geodesic projections to produce a final ‘consensus’ migration angle. We define the subset in polar geodesic projection by computing as the directional persistence score, the magnitude of the mean 3D geodesic superpixel track velocity of superpixel track *SP_i_*, 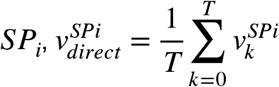 where the instantaneous 3D velocity: 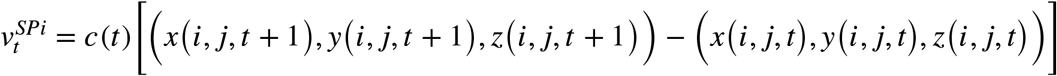 is defined as previously, after mapping back the 2D superpixel track positions back to Cartesian 3D. *c*(*t*) is the constant scaling correction factor from the rigid temporal registration at time-point *t*. The most directionally persistent subset of DVE classified tracks are those with a magnitude of 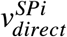 greater than 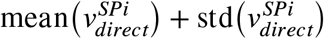 where mean (·) and std (·) are the mean and standard deviation operations. We post-process these tracks, keeping those within the largest connected component as the final subset, 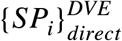. We find an equivalent subset in the Cartesian geodesic projection after remapping, in a manner similar to transferring the Cartesian DVE classifier results into the polar projection as described in the migration staging above. Given 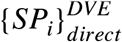 in polar and Cartesian projections we compute the angle as follows.

#### DVE Migration angle from polar geodesic projection

In the polar projection the migration angle is the equivalent 2D angle of the mean 3D velocity vector (Figure S3D). We compute the mean 3D velocity vector, 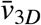 of the most directionally persistent subset of DVE superpixel tracks, 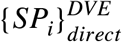 by mapping superpixel positions to Cartesian 3D. The mean 3D velocity vector, 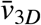 is then converted to a 2D velocity vector, 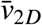 in the unwrapped polar projection. The migration angle, 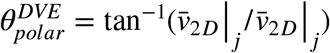 where 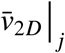 and 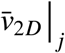 are the *i*-, *j*-direction component of 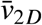. Conversion of 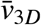 to 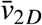 uses coordinate lookup and interpolation. We use the 2D superpixel tracks to compute the mean (*i, j*) coordinate position in polar projection to infer the mean (*x, y, z*) coordinate position on the Cartesian 3D surface. We then compute the position on the surface if we take a small displacement 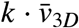 from the mean (*x, y, z*) coordinate where *k* = 10. This is mapped by nearest neighbour matching to an equivalent 2D (*i*′, *j*′) coordinate using the polar geodesic unwrapping coordinates, then 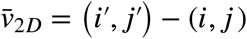.

#### DVE Migration angle from Cartesian geodesic projection

In the Cartesian projection, every column *j*-coordinate, corresponds to a unique radial angle and the row *i*-coordinate the proximal-distal embryo axis. Therefore finding the migration angle is equivalent to finding the *j*-coordinate with the highest velocity component in the ‘upward’ *i*- or proximal direction. However superpixels only sparsely sample the *j*-coordinate. Thus we divide the *j*-coordinate into 20 equal sized bins. We allocate each Cartesian extracted superpixel track in 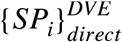 to one of the bins. For each bin, we then compute the mean 3D velocity component in the proximal direction,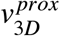 averaged over all superpixel tracks in the bin. This results in a plot of *j* vs 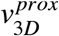. To get a more accurate approximation of the *j*-coordinate corresponding to maximum 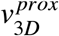 we fit a smooth univariate cubic spline and evaluate the spline at all integer increments of *j* ∈ [0, *N*] where *N* corresponds to the dimensions of the *M* × *N* pixel Cartesian projection. The migration angle is then obtained by converting the *j* coordinate of maximum 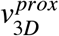 into a radial angle, 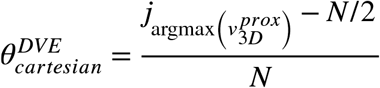 radians where the offset *N*/2 corresponds to a radial angle of 0 radians in the polar projection. The mean 3D velocity component in the proximal direction, 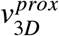 for a superpixel track *SP_i_* is computed as 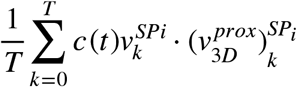 where 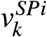 is computed as previously described, · is the dot product and *c*(*t*) the growth scale correct factor from registration. The directional vector in the proximal direction is computed pixelwise, 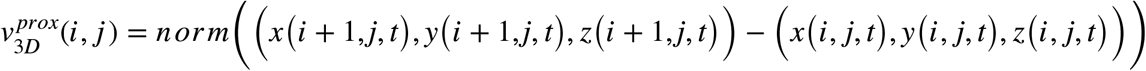 and 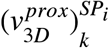 is then found by interpolation at the required (*i, j*) position at time *t* on the 2D superpixel track.

#### Consensus migration angle from polar and Cartesian projections

The angle mean of 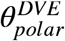 and 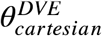 was taken as the consensus angle of DVE migration, *θ^DVE^*.

### Labelling and propagating the Em–Ex boundary throughout time-lapses

To ensure the Em-Ex boundary was correctly annotated we used VGG Image Annotator (VIA) (Dutta and Zisserman 2019) was to manually annotate the boundary between emVE- and exVE at the first time-point as a closed polygon in the unwrapped polar geodesic projection using the clear morphological difference between the epiblast and extraembryonic ectoderm tissues. The manually specified polygon coordinates were resampled using a linear spline to be 50 points. We then used the frame-by-frame computed DeepFlow optical flow field to propagate the 50 point polygon at time-point 0 to all later time-points. The final result is a *T* × 50 × 2 matrix for an embryo of total *T* frames.

### Partitioning the VE into sector region-of-interests (ROIs)

Individual embryos are heterogeneous in size, shape and rate of growth. We use the DVE migration stage to temporally align and DVE migration angle and the Em–Ex boundary to spatially partition each embryo in the polar geodesic projection (of size *M* × *M* pixels) into 32 sector regions-of-interest (8 radial angle bins and 4 geodesic distance bins) to allow consistent inter-comparison of DVE migration across embryos, Spatial partitioning for each embryo corresponds to the first frame of the migration phase. To spatial partition, we first use the inferred consensus migration angle, *θ^DVE^* to rotate the unwrapped polar projection such that the DVE cells migrate upwards in the vertical axis direction of the image and corresponds to *θ* = 0°^0^. We then partition the radial angle space *θ* ∈ [0, 2*π*] equally into 8 angular bins such that *θ* = 0° is the centre of the first bin; 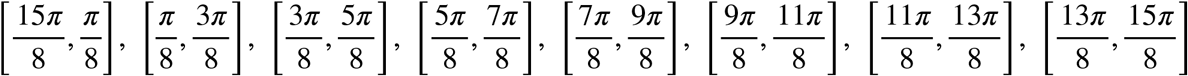. We partition the geodesic distance 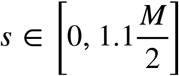, corresponding to the area between the central point and an outer concentric ring of distance 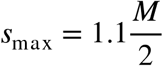 respectively equally into 4 intervals, guided by the Em-Ex-VE boundary; 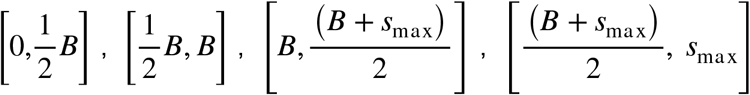, where the boundary is denoted as a function of geodesic distance and angle, *B* = *B*(*s, θ*). This further partitions both the emVE and exVE into 2 regions, along the proximal-distal (long) axis.

### Tissue motion deformation analysis

We measure the local tissue deformation as a consequence of DVE migration in each migration stage as the mean fractional rate of change in surface area, 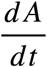 of a local tissue patch previously reported (Rozbicki et al. 2015) where 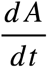 at time *t* is defined 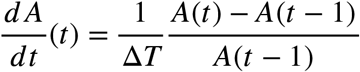 where Δ*T* = 5 min, the time elapsed between individual frames and 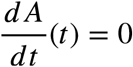 for time *t* =0. For visualization, we smoothed the 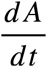 for each tissue surface patch temporally, running asymmetric least squares smoothing (Eilers and Boelens 2005)with parameters *p* = 0.5, *λ* = 100 for 10 iterations after edge-mode padding of signals by the maximum of *T*/10 or 3 frames. For each migration stage, we compute the 32 sector ROI as described in the previous section for the starting frame. The 32 sector is further subdivided equally by 5 angular and 4 distance intervals so that each quadrant is tessellated without overlap by 5×4=20 smaller quadrant ROIs. The subdivision parameters were chosen so that the small quadrant ROIs is a good quadrilateral approximation (same surface area) of the continuous corresponding curvilinear surface patch in Cartesian 3D. Each ROI region is modelled as a quadrilateral in 2D by linearly joining its 4 corner points; (*i*_1_, *j*_1_), (*i*_2_, *j*_2_), (*i*_3_, *j*_3_), (*i*_4_, *j*_4_) and similarly in 3D where (*i*_1_, *j*_1_) ↔ (*x*(*i*_1_, *j*_1_), *y*(*i*_1_, *j*_1_), *z*(*i*_1_, *j*_1_)) etc. The corner points of each of the 32 × 20 = 640 ROIs are propagated frame to frame until the end of the respective migration stage by the MOSES extracted DeepFlow optical flow. The 3D quadrilateral surface area at time 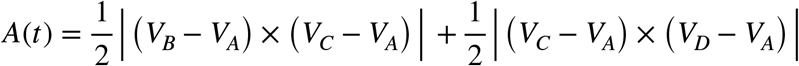, is the sum of the area of the two constituent planar triangles Δ*ABC* and Δ*ACD* where *V_A_, V_B_, V_C_, V_D_* are the vertex (*x, y, z*) coordinates of the quadrilateral *A BCD*.

### Single-cell segmentation

We used a semi-automatic scheme to segment the outline and temporally track the centroids of the apical surface of individual cells in the VE monolayer of 5 Lifeact-GFP embryos throughout pre-migration and migration phases in the 2D polar geodesic projection of time-lapse data imaged at five-minute intervals. This consisted of the development of a custom convolutional neural network (CNN) to segment cells in the densely-packed epithelium over-time, manual refinement and curation of the automatically segmented cells and semi-automated single cell tracking.

#### Spatiotemporal convolutional neural network (CNN) for single-cell segmentation

We trained a deep neural network on the Cartesian geodesic projections to segment the apical surface of individual VE cells for single-cell morphometric analysis. VGG Image Annotator (VIA) (Dutta and Zisserman 2019) was used to generate training data through manually annotation all cells in 75 time-points of Cartesian projected Lifeact-GFP data (7 embryos), comprising a training dataset of ~14,000 individual cell annotations. A custom UNet convolutional bidirectional LSTM neural network was used to predict the single-cell segmentation at time *t*, given the 3 sequential time slices at *t* – 1, *t, t* + 1.The neural network outputs three images to describe the predicted instance segmentation for time *t* in manner similar to Cellpose (Stringer et al. 2021). The first is a cell probability map that assigns a score of 0-1 to each (*i, j*) pixel of it containing a cell and such that cell centre pixels have the highest score of 1 and pixels lying on cell membranes having the lowest score of 0. The second and third images are the ‘*i*’-direction and ‘*j*’-direction displacements of each (*i, j*) pixel from the unique cell centre that it is predicted to belong to. Pixels located at cell centres by definition have ‘*i*’-direction and ‘*j*’-direction displacements of 0. We construct the supervised training for the three outputs from the manually annotated instance segmentation labels. The first output, the probability map is the composition of the normalised (0-1) distance transform of each unique cell (segmentation label). Similarly, the second and third outputs, the *x*- and *y*-direction displacements is the composition of the image gradient of the distance transform of each unique cell (segmentation label). The neural network effectively learns a deep watershed, direct from the data itself (Bai and Urtasun 2017; Stringer et al. 2021). The network is trained with a multi-task loss function; a softmax loss function to enforce mutual exclusivity of background and foreground regions in the cell probability map, the first output and L1 loss for the *x*-, *y*-direction displacements, with an uncertainty weighting scheme to automatically improve training (Kendall et al. 2018). The dataset was split into annotated frames from 4 unique embryos for training and annotated images from 2 unique embryos for testing. Frames from the last embryo were equally shared, frames 18-22 for training and frames 3-7 for testing. This gives n=30 (from 50 unique frames) and n=15 (from 25 unique frames) 3-frame images for training and testing the CNN respectively. Images were resized to a pixel size of 512 × 640 and were augmented in real time during training and validation. The set of augmentations include random left-right, up-down flipping, piecewise affine deformation, Gaussian blur, average blur, median blur, additive Gaussian noise, pixel and coarse dropout and intensity and contrast manipulations. We parse the individual cell segmentations from the predicted outputs by Euler integration similar to that used in Cellpose (Stringer et al. 2021). An *M* × *N* image corresponds to a uniformly seeded grid of points located at (*i, j*) row, column coordinates. Likewise, the predicted ‘*x*′-, ‘*y*′-direction displacements corresponds to the (Δ_*j*_(*i, j*), Δ_*i*_(*i, j*)) displacements respectively that we should displace the point at (*i, j*) to a new location (*i*′, *j*′) ← (*i* + *α*Δ_*i*_(*i, j*), *j* + *α*Δ*j*(*i, j*)) where *α* is a constant step size in pixels that we take in the predicted direction. Iteratively updating the location such that (*i*′, *j*′) is the new starting position for the next update, (*i, j*) ← (*i*′, *j*′) for each point will eventually enable all points to converge to the unique cell centre it is predicted to belong to by the neural network. Individual cell centroids correspond to non-spatially connected regions of high point density which were found by thresholding and connected components analysis. Tracing back to the starting position of individual pixels which voted for a particular cell centroid retrieves the individual instance cell segmentations over the whole image. We iteratively advected (*i, j*) pixels for 30 iterations to generate the cell segmentations in the Cartesian or 50 iterations for the polar geodesic projection.

#### Manual curation of CNN-predicted single-cell segmentation

The unwrapped 2D Cartesian and Polar geodesic projection enables visualization of every cell in the VE surface in a single image at each time-point. As each projection (Cartesian and polar) necessarily distorts cell appearances in different regions of the VE cell sheet (greater at the distal-tip and proximal regions, respectively), it can result in over-segmentation. To edit over-segmentations resulting from the automated outlining manual proofreading and correction of CNN segmentations was necessary. This was done in 3 steps to obtain the final corrected CNN segmentation in the polar projection. Instance segmentations were ‘inverted’ to obtain binary membrane outlines for easier manual correction when comparing to the Lifeact-GFP signal. Step 1: Check and correct individual cell outlines for all cells in the Cartesian projection except those around the distal tip and all cells at the distal tip in the polar projection. Step 2: remap Cartesian manually corrected cell outlines into polar view and combine with polar manually corrected cell outline. Step 3: check for any inconsistencies in the combined polar projection. Step 4. Convert membrane cell outlines to instance segmentation by binary pixel inversion and connected component labelling.

### Automatic single cell tracking and manual correction for cell division

The single cell segmentations in individual frames need to be linked temporally such that all occurrences of the same unique cell in different frames are captured in a single cell track. We regard mothers and daughters as unique individuals such that the mother and two daughters constitute 3 separate cell tracks. The cell segmentations were first linked automatically using by nearest neighbour tracking of cell centroids. Every cell centroid at time-point 0, the first time-point of the time-lapse and of the pre-migration stage is initialised as a unique cell track. We then compute the pairing of all cell centroids in the next frame to the last cell centroid in each unique cell track that results in the minimum total 3D Euclidean distance using bipartite matching (Kuhn 1955). Centroids that could not be matched begin new tracks. Tracks that cannot be matched are terminated. The process is repeated until the end of the migration stage for each of the 5 Lifeact-GFP embryos. The automatic linking does not explicitly account for cell divisions. Manual proofreading was subsequently undertaken to enforce cell division and correct for erroneous linkage using a custom graphical user interface, “Cell Tracker”. Cell Tracker enables visualization of the history of all or a subset of cell tracks overlaid on the 2D projections and the ability to delete or create new tracks and to create associations between cell track IDs. Cell divisions were annotated as single time-point cell tracks (see cell division annotation below) and used to break up the automatic tracks after manual refinement: daughter tracks are assigned to a mother track, and combined to form continuous track in the dataset.

#### Single-cell phenomic characterisation

For each segmented and tracked cell, we extracted fourteen measurements to describe the instantaneous morphodynamics of its instance at time *t*; two dynamic (VE Anterior speed, cumulative VE anterior distance), five planar/surface (surface area, cell perimeter, shape index, aspect ratio, number of cell neighbours), one depth (cell height, four relating to global embryo context (VE apical, apical Gauss, basal and basal Gauss surface curvatures), and two related to Lifeact-GFP/F-actin signal (area and perimeter apical Lifeact-GFP/F-actin intensity) along with their lineage annotation and embryonic-spatial position within the radial grid schema. All dynamic measurements were measured for time *t* if an instance exists at time *t* + 1. This means all cell segmentation instances in the last frame of pre-migration and migration stages do not have dynamic measurements. Dynamic measurements were computed after transformation back to 3D coordinates and multiplicative scale factor correction between time-points to account for growth. All geometrical measurements were transformed to 3D coordinates and multiplicative scale factor correction with additional with reversal of non-rigid registration parameters to calculate real absolute values (minus motion artifacts from rotation and translation). We use Lifeact-GFP pixel intensity to quantify actin. Area actin is the mean Lifeact-GFP intensity within the cell area. Perimeter actin is the mean Lifeact-GFP intensity along the cell perimeter. The raw Lifeact-GFP intensity varies across embryos and increases over time. We therefore use the z-score instead of the raw intensity with the per frame mean and standard deviation. Together, the fourteen measurements, summarised in Table S2 describe the instantaneous phenotypic-state of single VE cells in the time-lapse during pre-Migration and migration phases, integrating both planar and 3D information.

To compute average cell height, *K*_Basal_ and *H*_Basal_ we unwrapped for each Lifeact-GFP embryo an additional manually annotated basal VE binary volume, taking into account the shape of the underlying epiblast using the same method described for apical VE above. As the apical and basal binary volumes are similar in shape, we put the two unwrapped coordinates in alignment by resizing the basal projections and unwrapping coordinates to match that of the corresponding apical VE. We then used the same apical VE single cell segmentations to extract single cell statistics of the apical-basal height of the cells. As each cell is not 3D segmented, this computed cell height metric provides an approximation of cell height as it measures the apical-basal height of the local VE tissue.

### Single-cell UMAP phenomic space

Analysis of single-cell parameters is a multi-step process involving the initial extraction of data from all datapoints followed by a filtering step to remove invalid or missing values. From five Lifeact-GFP embryos we initially extracted a total 102,872 cell instances (data-points) with a complete set of fourteen measurements (phenomic signature), tracking a total of 2358 unique cells and incorporating 447 division events. Cell instances with any invalid or missing values i.e., *inf* or *nan* value in any measure were removed leaving a total of 91,901 cell instances tracking a total of 2221 unique cells, with a total of 447 divisions events across the data The phenomic signature of these remaining cell instances was preprocessed to normalise each measure and corrected for potential individual embryo-specific effects. Each measure is power-normalised and z-scored relative to all instances from the embryo it belongs to. The mean is computed from the data for non-signed measures: area, perimeter, shape-index, aspect ratio, cell height and cell neighbours and is set to 0 for signed measures: anterior speed, cumulative anterior distance, Gaussian apical surface curvature, apical surface curvature, Gaussian basal surface curvature, basal surface curvature, apical areal and apical perimeter actin intensities. The zscored measures reflect the extremity of each measure relative to that of a mean cell instance from the same embryo as a multiplicative factor of standard deviation. We remove all cell instances which has any measure with an absolute zscore value > 3. For the remainder cell instances we compute for each of the eleven measures, a value corrected for embryo (batch) - the residual *r_i_* of linearly regressing measure *i*, *F_i_* as the dependent variable with embryo, *Emb* encoded as an independent categorical variable, *F_i_* ~ *C*(*Emb*) + *r_i_*. We apply UMAP (Becht et al. 2018; McInnes et al. 2018) to the normalised and embryo corrected phenomic signatures to plot all remaining 91,901 cell instances, as one datapoint per instance, across all embryos, all time-points into a jointly shared 2D coordinate space - the morphodynamic phenomic space for comparative analysis. This 2D space captures the multidimensional phenomic differences between all individual cell instances over space-time during pre-migration and migration stages according to the eleven-dimensional morphodynamic signature.

### Automatic UMAP phenomic clustering

To automatically determine the number of unique phenotype clusters we first grouped all cell instances into 100 clusters using k-means clustering on the 2D umap coordinates as the input features to cluster on. Hierarchical clustering with Ward linkage and Euclidean metric was then applied to group the 100 k-means clusters into phenotype clusters using automatic cluster determination. The input feature vector per k-means cluster for hierarchical clustering is the histogram of the quadrant ID of all data points. The histogram describes the fraction of all data points assigned to a k-means cluster belonging to all possible quadrant IDs. The quadrant ID uses the reduced 8 sector ROI scheme which aggregates the anterior and posterior regions of the 32 sector ROI. Cell instances were assigned to an 8 sector ID based on cell origin according to the position of the unique cell it belong to at the first frame of migration stage – lineage quadrant ID. Cell instances were also assigned a quadrant ID according to their instantaneous position in the embryo – instantaneous quadrant ID. The k-means cluster summary features is sixteen-valued; the concatenation of the histogram based on lineage and instantaneous quadrant IDs. Data points corresponding to lateral VE quadrants are therefore not included and do not influence the hierarchical clustering. Hierarchical clustering generates a dendrogram such that the root node corresponds to all k-means cluster grouped as one label, and progressive branch splitting corresponds to subsplitting the larger parental grouping into unique smaller groupings. The bottommost leaves of the dendrogram are the 100 individual k-means clusters. Automatic cluster determination is equivalent to finding where to cut the dendrogram. This was based on finding the minimum number of groups that grouped k-means clusters in a stable and spatially homogeneous way. We operationalise this definition by finding all potential groupings of the k-means clusters when placing cuts up to a maximum linkage distance (to constrain the search). For each grouping, we compute a homogeneity score to summarise the tendency to group k-means clusters coming from the same regional quadrant ID together. Specifically, the consistency of a k-means cluster is quantified by the fractional dominance of any single quadrant ID given by the maximum value in the sixteen-valued histogram vector. We plot the homogeneity score as a function of the linkage distance. A grouping is stable if over a range of linkage distance the homogeneity score does not change. We detect and use the first linkage distance to generate the first possible stable grouping. This produces a total of 6 phenotype groups. We retain groups as the final confident groups if the homogeneity score > 0.3. The groups that do not satisfy this criteria are identified as a single large miscellaneous group. We find in total 5 distinct phenomic clusters, designated A-E which we could relate to spatial regions of the embryo; cluster A: anterior DVE 1, cluster B: anterior DVE 2, cluster C: posterior emVE, cluster D: proximal anterior emVE, and cluster E: grouping representing the mean emVE and exVE. For visualization in polar geodesic projection, we use label spreading (D. Zhou et al. 2004), a semi-supervised machine learning technique to impute the phenomic cluster of cell instances not included in the original UMAP by its multidimensional similarity to the nearest 15 cell instances (15 nearest neighbours). The algorithm is iterative so that the first iteration labels unknown cell instances which are closest to cells used in the UMAP and successive iterations gradually propagates the labels to cells of unknown labels transductively.

### Sector-based UMAP phenomic fate trajectories

Phenomic space is a two-dimensional representation of a continuum of morphodynamic states. Construction of a temporal phenomic fate trajectory representing the evolution of the average behaviour of cells starting in each of the 8 spatial sectors overtime was carried out as follows: 1) bin all datapoints starting in sector *i* into 6 equal temporal intervals from −2h to +4h of migration, 2) for each temporal interval, derive a spatial heatmap of the number of datapoints (density) mapping to any local region in phenomic space, 3) find the spatial regions of highest datapoint density by thresholding on mean + 1 standard deviation of density, 4) compute the mean UMAP coordinate of the high density spatial regions, 5) repeat steps 1-4 for all temporal intervals and join together the UMAP coordinates in chronological order to form the phenomic fate trajectory.

### Phenomic correlation between sectors across migration stages

We compute the Lasso regression coefficients (partial correlation) treating each of the fourteen statistics in turn as the independent variable and the other thirteen statistics as the dependent variable. We use the Python *statsmodels.regression.linear_model.OLS.fit_regularized* function with L1 penalty = 1, and the normalized and embryo corrected phenomic signatures to do the Lasso regression. The result is a 14 × 14 coefficient matrix. We compute such 14 × 14 matrices for each of the 8 sectors labelled according to cell lineage and separating out pre-migration and migration stages. This gives 16 matrices in total. Each 14×14 matrix, *C_s_* summarises the interdependency between single cell statistics as captured by the cell instances in each sector *s*, in each migration stage. To assess the phenomic correlation between sectors across migration stages, we applied hierarchical clustering, Euclidean metric, complete linkage to the 16×16 pairwise difference matrix between the 16 matrices. We define the pairwise difference between two coefficient matrices, *C_i_* and *C_j_* is the total sum of the absolute difference in matrix entries, ∑|*C_i_* – *C_j_*|. Ia similar manner we assessed the phenomic correlation between sectors and migration stage when cell instances were instead labelled by their instantaneous position.

### Single measure time-series

The time-series for a single chosen measure in Figure S5B-E, and Figure S7D and quadrant ID origin is the mean and standard error of the mean (s.e.m) of all the datapoints within the given time interval, *t_start_* < *t* < *t_end_*. The mean and s.e.m is computed on the raw data, not the batch embryo-corrected and preprocessed data used for defining the UMAP.

### Single-cell track diffusion coefficient

The mean squared displacement (MSD) of individual single cell tracks were computed using the *imsd* function in the Python *Trackpy* package using a maximum lag time of one quarter the maximum video length. The mean squared displacement is related to the diffusion coefficient *D* in *n* dimensions of freedom by the formula MSD = 2*nD t* where *t* is the time lag. *D* is estimated for each cell track using *n* = 2 (the cell track migrates effectively constrained to the 2D surface) using linear regression.

### Cell division annotation and analysis

Cell divisions were manually annotated using custom designed software “Cell Tracker” Due to Lifeact-GFP condensation at the site of cytokinesis, cell division events could be clearly seen as bright puncta in the time-lapse data. Using the Cell Tracker software the 2D surface projections could be used to efficiently screen through the time-lapse data for divisions. The centroid of the mother and corresponding daughter cells of the post-processed time-lapse data were manually inputted along with the timing of the division event, anatomical position and linked to its cell track. Division angles were computed after remapping the individual 2D daughter cell centroids to 3D. For analysis we computed the angles relative to the proximal-distal axis as follows; a division resulting in a line drawn between daughter cell centroids when aligned perpendicular to the proximal-distal axis is given as 0° , if aligned orthogonal to the proximal-distal axis (i.e., perpendicular to the radial axis) the division angle is given as 90°. Negative division angles were inverted (i.e., −45 to +45) for comparison, so that all angles were converted to a range of 0°-90°. For analysis all cell division events from 9 Lifeact-GFP embryos per selected anatomical region were combined, and binned into 4 groups (0-22.5, 22.6-45, 45.1-67.0, 67.1-90). A Chi-squared test for given probabilities was then performed. To analyse the frequency of cell division events we combined all cell division events occurring in the anterior proximal emVE and separately, the posterior proximal exVE during DVE migration. We performed a Chi-squared test for significance from expected probabilities. To analyse the timing of cell division events with respect to the onset and duration of DVE migration, we binned cell division events from anterior proximal emVE and posterior proximal exVE into 10 bins according to the time of migration (both prior to and during DVE migration). A Chi-squared test was performed to test for significance from expected probabilities. We note that the random distribution of angles on curved surfaces such as a sphere does not follow a uniform distribution (Cai et al. 2013), it does not apply here as the radius of curvature is much larger relative to the cell size at the distal tip of embryos and therefore the surface is locally flat - enabling us to use the uniform distribution.

### Cell intercalation events

Cell intercalation events whereby a pair of neighbouring VE cells were separated by a third cell moving between them, and not associated with a cell division event, were manually annotated using the Cell Tracker GUI (above). 2D surface projections of post-processed time-lapse data were screened through for incidences of intercalation events and centroids of the cell trio; neighbouring cells 1 and 2, and the intercalating cell, were inputted, along with their timing, anatomical position and cell tracking IDs. The cell tracking ID could then be use to cross-reference with the instantaneous cell status (e.g., phenomic cluster).

### Visualization of cell statistics in 2D projections

Each segmented cell instance has a unique cell ID. To visualize the desired scalar statistics of a segmented cell instance, the scalar value was linearly mapped to a Python *Matplotlib* colorscheme to colour the entire cell area in the 2D projection; ‘Reds’ colorscheme for non-signed measures: area, perimeter, shape-index, aspect ratio and cell neighbours and ‘coolwarm’ colorscheme for signed measures: anterior speed, cumulative anterior distance, Gaussian surface curvature, mean surface curvature, apical area-, and apical perimeter actin intensities.

### Visualization of cell statistics on 3D meshes

We visualize the desired scalar statistics of a segmented cell instance in 3D using MeshLab. (Cignoni et al. 2008). Meshlab requires as input a triangle mesh (trimesh). We use Python *Trimesh* library to write a .obj trimesh which requires a list of 3D (*x, y, z*) vertex coordinates, a list of faces, 3-tuple specifying how vertex indices are connected together in triangles and a list of vertex colours, 3-tuple specifying the RGB colour at each vertex. Cell statistics were first visualised in the 2D projections as an image as described above with cell tracks drawn as 3-pixel wide lines using Python *Scikit-Image.* A 2D image is equivalent to a quadrilateral mesh specified by a list of 2D (*i, j*) vertex coordinates, a list of quadrilateral faces, 4-tuples specifying how individual pixels are connected to neighbours in squares and the 2D image pixel colour as the RGB vertex colour. To obtain the desired 3D trimesh, we triangulate the quadrilateral mesh to a trimesh using the Python *Trimesh* library (*trimesh.geometry.triangulate_quads* function); remove all (*i, j*) vertex coordinates and faces not part covering the VE in the polar geodesic projection; and remap the 2D (*i, j*) coordinate to 3D (*x, y, z*) vertex coordinates using the 2D-to-3D unwrapping coordinates (see above).

---

**Figure S1.**
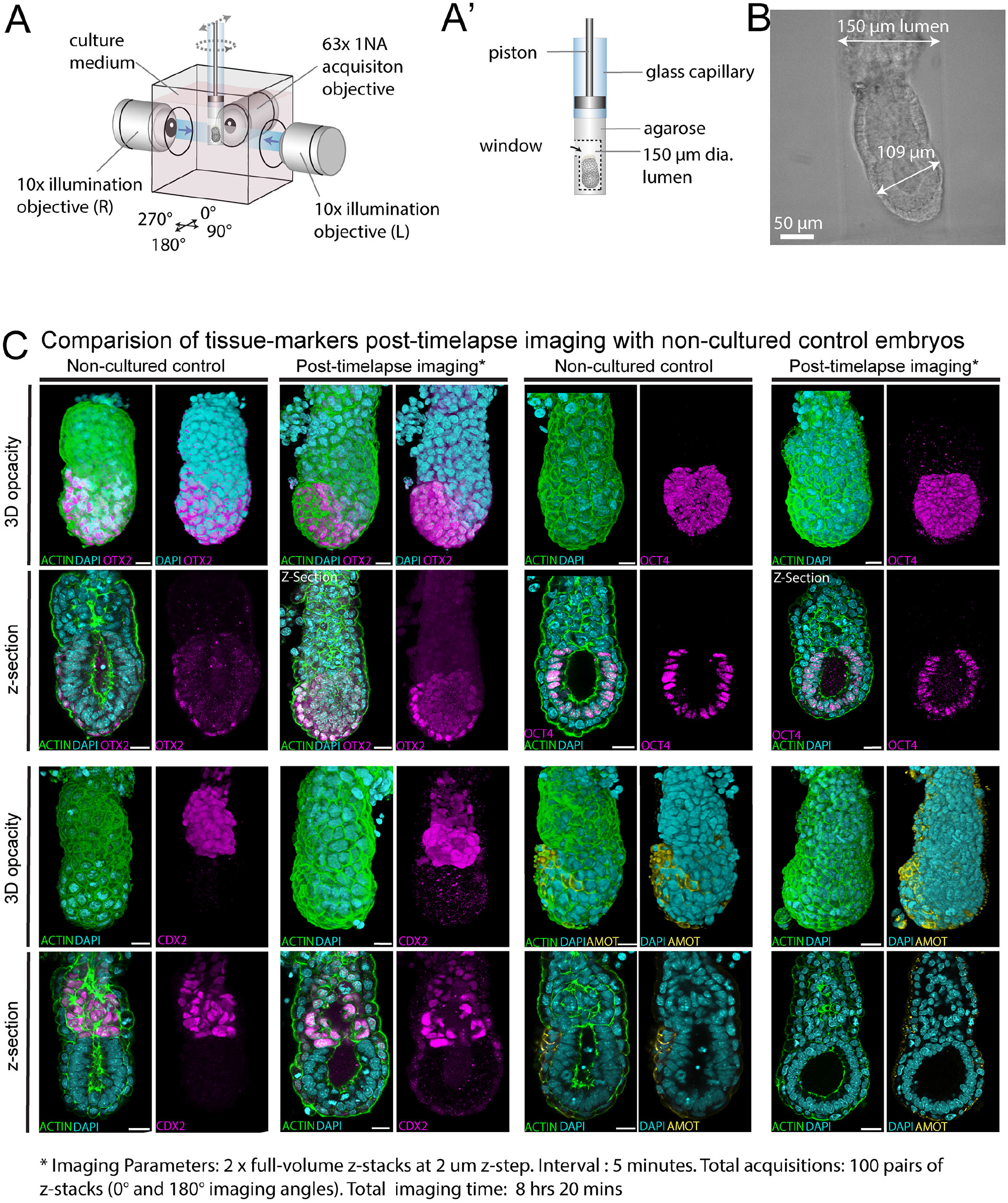
Lightsheet imaging set-up and expression of tissue markers in imaged embryos. **A.** Diagram of the imaging chamber of the ZEISS Z.1 lightsheet microscope. **A’.** Diagram of an E5.5 embryo mounted for imaging. **B.** Brightfield image of an E5.5 embryo mounted for imaging in the lumen of an agarose cylinder. Note the embryo is placed within a lumen wide enough so that it does not constrict growth. **C.** Whole-mount immunofluorescence of control non-cultured E5.75 embryos and Lifeact-GFP E5.5 embryos imaged for 100 time points (>8 hours) at five-minute interval in a ZEISS Z.1 microscope. Imaged embryos show similar expression patterns of DVE (OTX2, AMOT), epiblast (OCT-4) and extraembryonic ectoderm (CDX2) markers to controls. Phalloidin (green) and DAPI (cyan) staining show that the VE remains as a monolayer epithelium. All scale bars = 25 μm.

**Figure S2.**
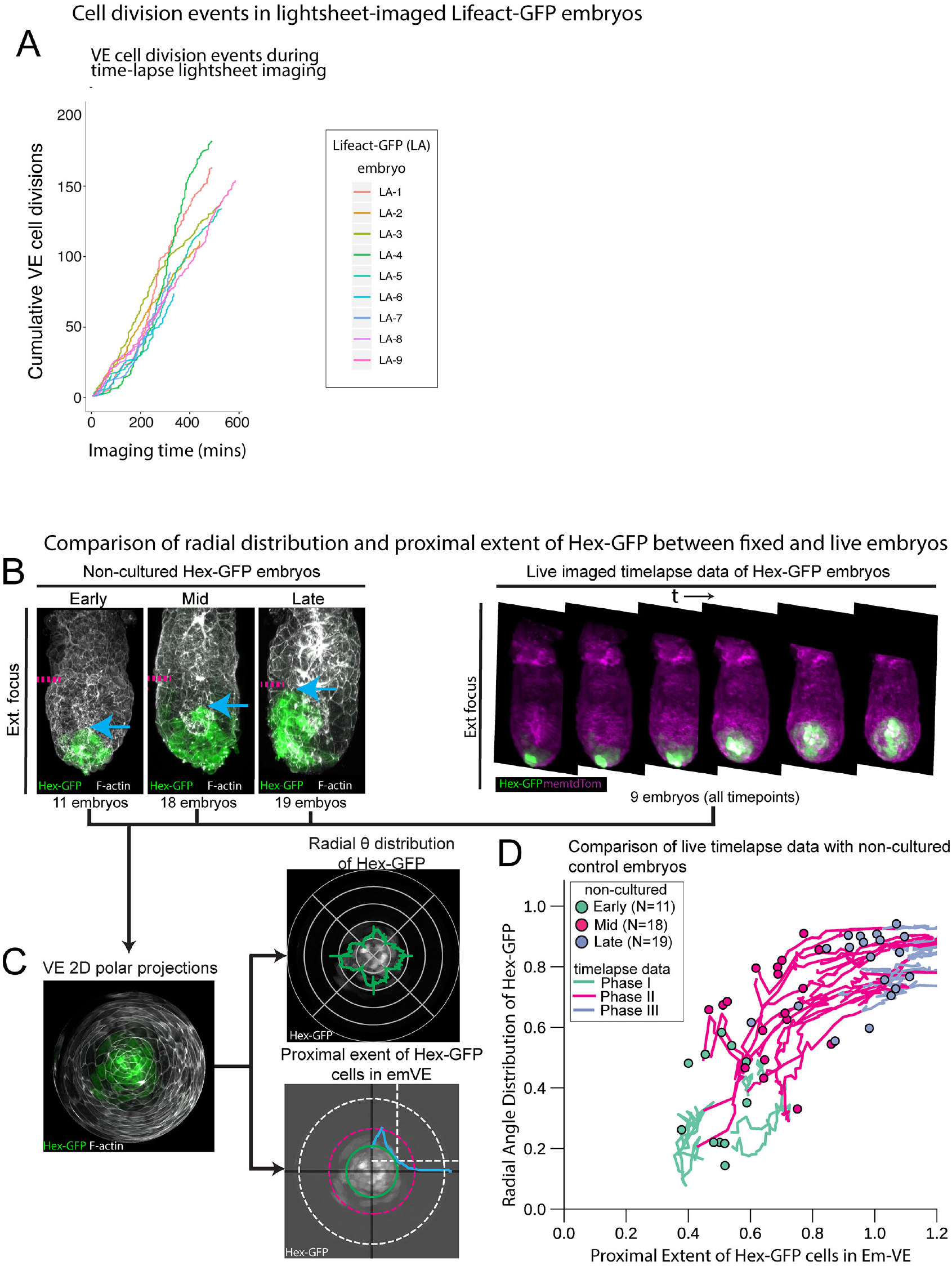
Controls to assess health of imaged embryos – cell division events and comparison with non-cultured stage-series reference embryos. **A.** Cumulative VE cell division events for each of 9 Lifeact-GFP embryos imaged in a ZEISS Z.1 ligthsheet microscope from 2 view angles every five-minutes with a 2 μm z-step interval show that cell divisions continue throughout imaging. **B to D.** Lightsheet imaged embryos were compared to a stage-series of fixed embryos with respect to the radial distribution and proximal extent of the Hex-GFP cells during the course of the time-lapse. **B.** At left, representative examples of embryos fixed at early, mid-, and late stages of migration, with migrating DVE cells marked by Hex-GFP expression (green) and all cell outlines visualised with Phalloidin labelled F-actin (white). At right, full volume confocal z-stacks of select time-points of a representative, live imaged Hex-GFP:membrane-tdTomato embryo, capturing different stages of DVE migration. **C.** 3D confocal data of fixed embryos, as well as time-lapse live data were reprojected as 2D polar projections. At left, a representative example of a non-cultured, fixed embryo is shown. To enable a quantitative comparison of DVE migration in embryos, the radial distribution of the Hex-GFP expressing DVE cell population was calculated for each non-cultured embryo, and for each time-point of live imaged embryos, with; 0 = radially symmetric, 1 = polarised (top panel, green line). To assess how far DVE cells had migrated in each embryo, the proximal extent of the Hex-GFP cell population in the emVE was calculated for each non-cultured embryo, and for each time-point of live imaged embryos. As embryos differ in size, the proximal extent of migration was expressed as a fraction of the distance from the distal-tip to emVE-exVE boundary for each embryo (lower panel, magenta line = emVE-exVE boundary, green line = proximal extent of DVE, blue line = Hex-GFP intensity). **D.** The radial distribution of DVE cells was plotted against their proximal extent for fixed and cultured embryos. The stage series of fixed embryos is represented by coloured dots (green = early migration, magenta = mid-migration, purple = late migration). Time-lapse imaged cultured embryos are represented by solid lines, coloured by phase of migration (phase I = pre-migration, phase II = migration and phase III = late migration). The Hex-GFP cell population in live imaged embryos progress through a radial distribution and proximal extent within the range of non-imaged, control embryos showing that imaged embryos reflect normal development.

**Figure S3.**
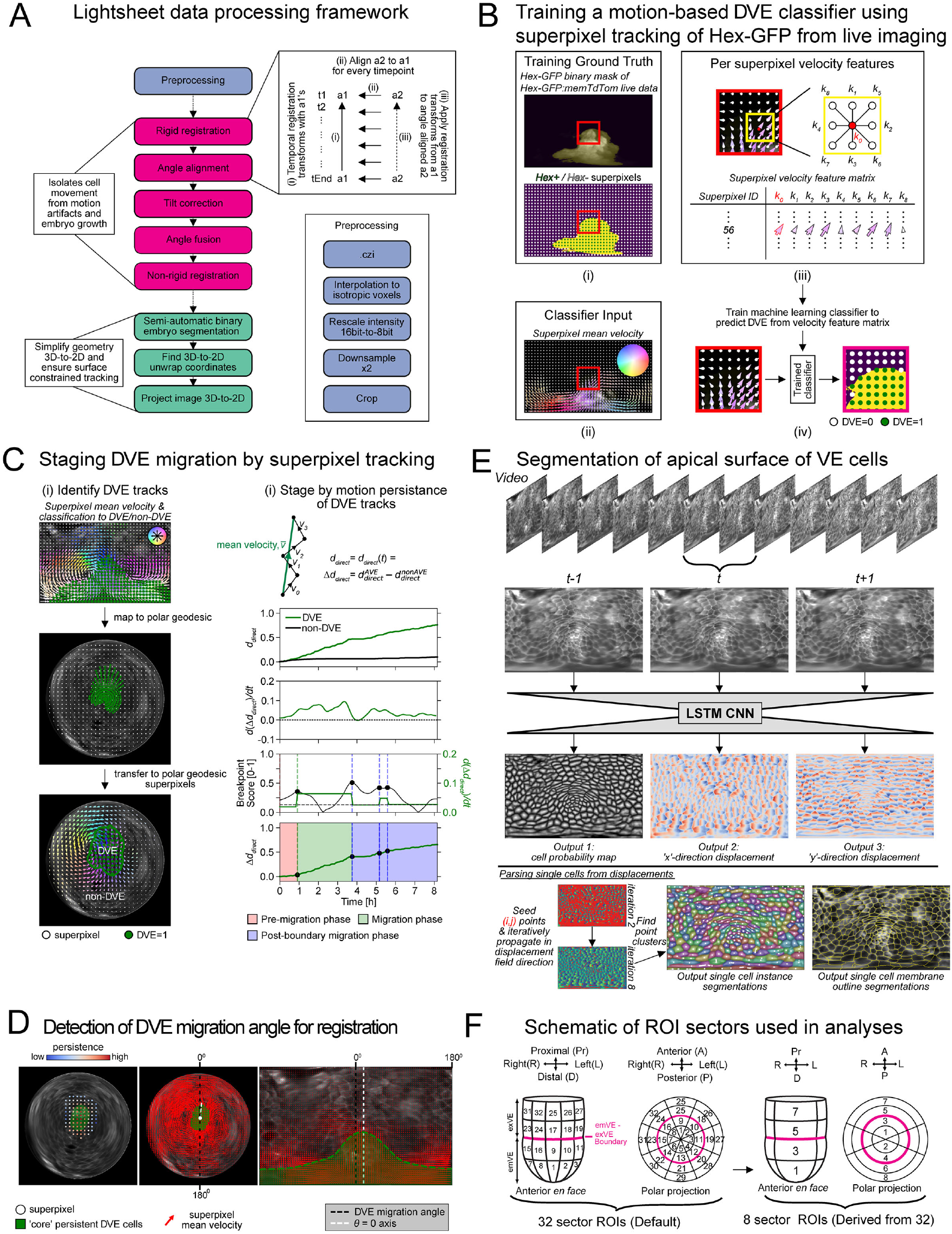
Lightsheet data processing framework. **A.** Overview of the analytical steps to generate a consistent surface coordinate framework for a two-angle volumetric embryo time-lapse, and to project the apical surface of the VE to a 2D geodesic surface projection in a reversible manner. **B.** Overview of the key steps to train a superpixel-based motion classifier based on Hex-GFP embryos, to be used to identify DVE cells in Lifeact-GFP data. This includes: (i) initial binary segmentation of the Hex-GFP signal to label Hex-GFP+ve / Hex-GFP -ve associated superpixels for training; (ii) using the mean superpixel velocity as a consistent motion feature across Hex-GFP and Lifeact-GFP for classification; (iii) construction of the velocity features to classify each superpixel using the mean velocity of itself and its surrounding eight neighbours; and (iv) training a binary machine learning classifier to classify each superpixel given the velocity features as either DVE=0 (not DVE) or DVE=1 (DVE). **C.** Overview of the key steps to automatically stage DVE migration from superpixel-based motion tracking including: (i) applying the trained DVE classifier to identify DVE-associated tracks in polar-geodesic projection and (ii) identifying a continuous time window when DVE associated superpixels exhibit significantly greater directional motion persistence than non-DVE associated superpixels (Migration phase, green) to classify the remainder time into Pre-migration (red) and Post-boundary (blue) phases. **D.** Detection of DVE migration as the mean consensus angle between the mean velocity direction of the ‘core’ persistent subset of DVE cells in polar and Cartesian-geodesic 2D projections (white dashed lines) given relative to 0°-180° line (black dashed lines). The DVE migration angle is used to spatially align the projections from multiple embryos with respect to the future anterior-posterior and left-right axes. **E.** Training of a LSTM-Convolution neural network to automatically segment individual VE cells from 2D projection images in each time point, integrating the temporal information from times −1 and +1. **F.** Schematic of the 32 sector and 8 major sector anatomical region-of-interest (ROI) spatial partitioning of each embryo relative to the anterior-posterior axis, given by the consensus DVE migration angle found in D). This consistent subdivision enables integrative analysis of statistics across multiple embryos.

**Figure S4.**
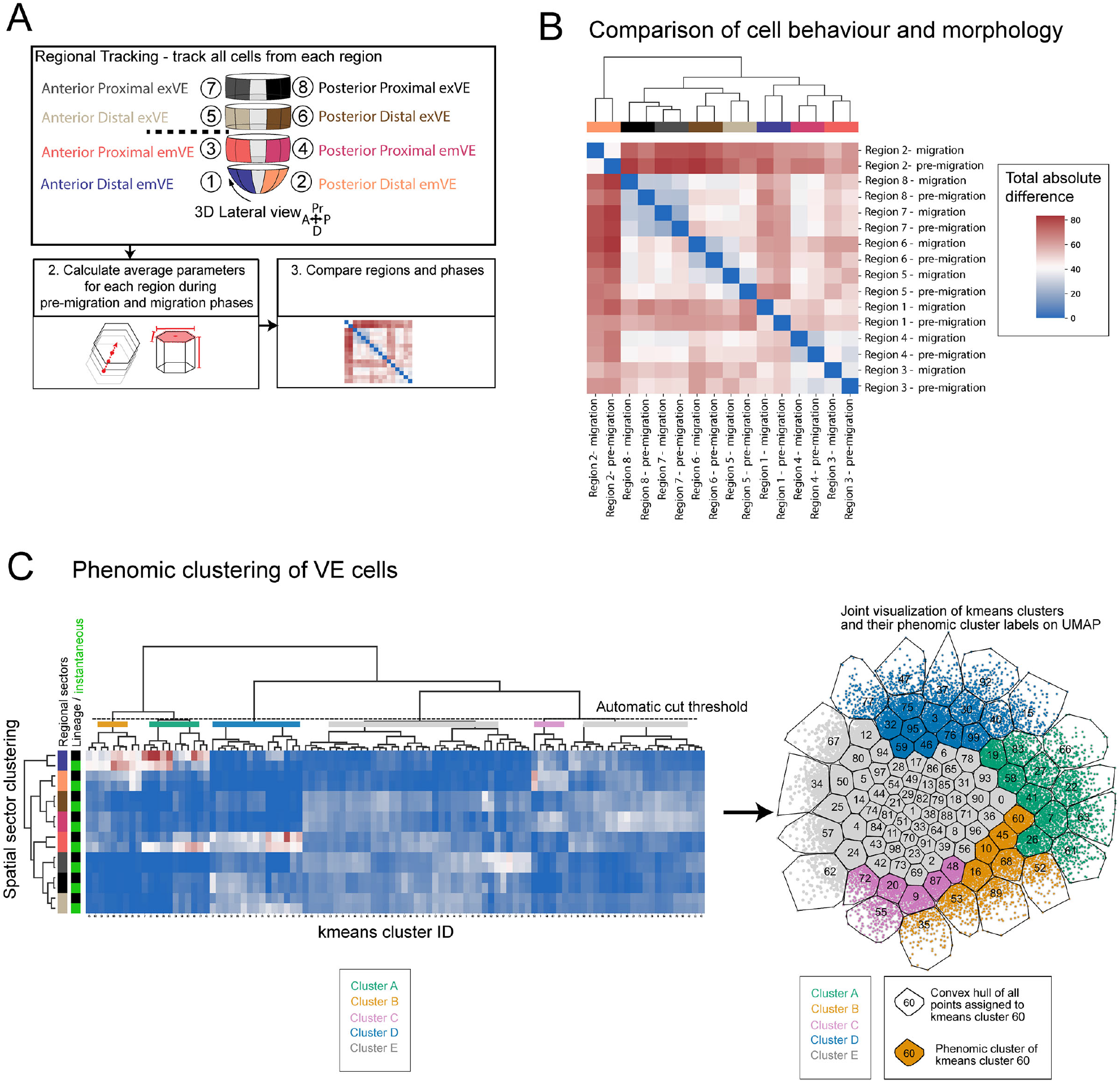
Single-cell phenomic analysis and hierarchical clustering. **A.** Overview of the analytical steps to compare phenomic behaviour across regions and phases based on the similarity of the 14 parameters. **B.** Hierarchical clustering of regions and phases based on the similarity across the 14 parameters, given by the absolute difference in the partial regression coefficients matrix amongst parameters (see Methods). All regions show the same behaviour between pre-migration and migration. exVE regions clustered together. emVE regions also clustered together except the posterior distal emVE (Region 2) which appears distinct. **C.** Hierarchical clustering of cell instances into phenomic clusters with the number of phenomic clusters determined by an automatic cut threshold (left, see Methods). Cell instances were first coarsely grouped into 100 coarser k-means clusters based on their UMAP coordinate (which is also a non-linear average of the 14 parameters) to minimise the effect of individual cell heterogeneity. Hierarchical clustering was then applied to the k-means clusters based on the anatomical origin distribution given by tracking (lineage) and instantaneously. The clustering checks individual phenomic clusters for consistency in anatomical origin which is used as a measure of uncertainty and marks all clusters with high variability as an average VE cell, cluster E (grey). The resultant phenomic clusters, overlaid on the k-means clusters (numbered and demarcated by black outlines) are thus determined based on phenomic and anatomical consistency (right).

**Figure S5.**
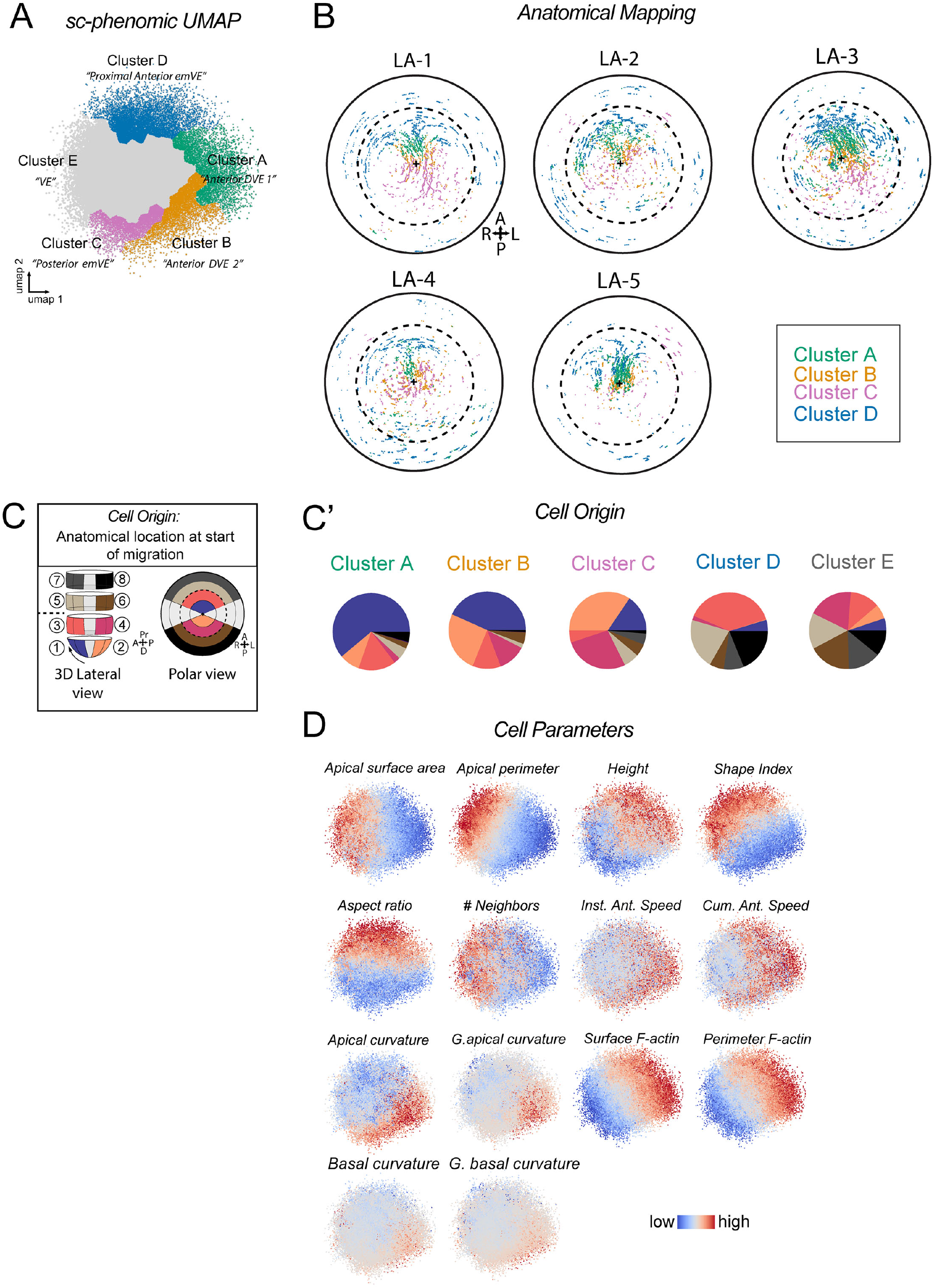
VE Single-cell phenomic UMAP. **A.** VE single-cell phenomic UMAP comprising of all instances of all VE cells from five Lifeact-GFP embryos, showing 5 phenomic clusters. **B.** Anatomical mapping of all instances of all cells in phenomic clusters A–D onto the respective polar projection of each of the embryos that they were measured from. **C.** Diagram showing each of the 8 anatomical sub-divisions of the VE used in this study. **C’.** Charts showing the proportion of instances of all VE cells from each anatomical location (from C) in each of the UMAP clusters (from A). **D.** VE single-cell phenomic UMAP coloured by intensity (red high, blue low) for each of the 14 cell parameters quantified.

**Figure S6.**
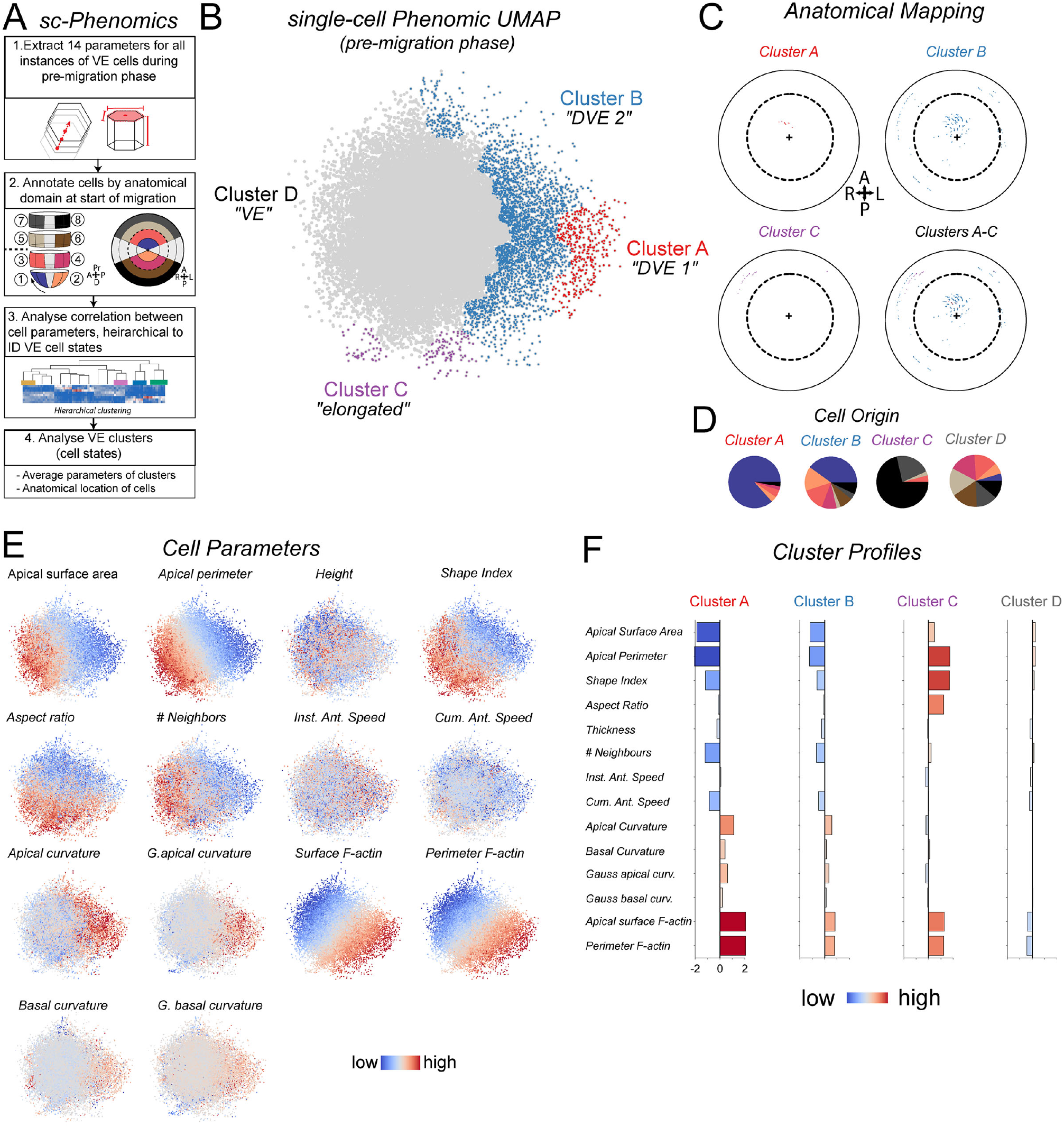
Pre-Migration phase single-cell phenomic UMAP. **A.** Summary of the single-cell phenomic analysis method applied to the pre-migration phase. **B.** VE single-cell phenomic UMAP comprising of all instances of all VE cells from the pre-migration phase only, of embryos based on the 14 measured cell parameters. **C.** Example of the anatomical mapping of one embryo in the dataset showing all instances of clusters A-C on a 2D polar projection. **D.** Charts showing the proportion of instances of all VE cells from each anatomical location (A) in each of the UMAP clusters (B). **E.** VE single-cell pre-migration UMAPs showing each of 14 cell measurements highlighted. F. Mean profiles of each pre-migration cluster for each of the 14 cell measurements.

**Figure S7.**
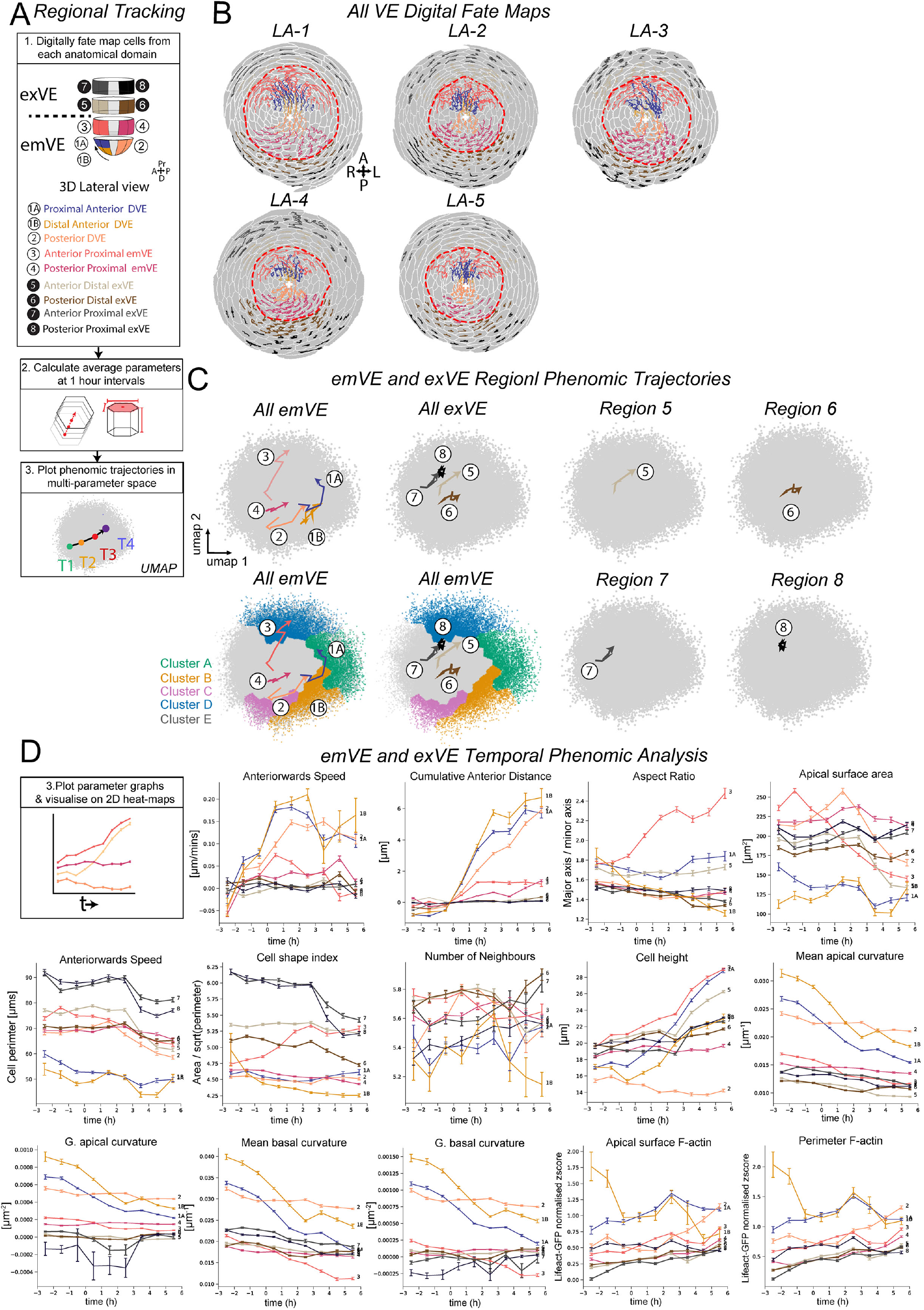
Tracking and phenomic trajectory analysis of emVE and exVE sub-regions during DVE migration. **A.** Summary of the analytical steps for all VE sub-regions in digital fate-mapping, phenomic trajectories and temporal analysis. **B.** Continuous tracking of cells from five sub-regions of the emVE and four sub-regions of the exVE during the migration phase of all embryos analysed. Tracks towards the periphery are exaggerated in length due to the distortion inherent in the projection. **C.** Average UMAP position of VE cells starting at each emVE and exVE subregion at 1-hour intervals in the multi-parameter phenomic UMAP. **D.** Temporal plots of of mean±s.e.m for 14 cell parameters in all five emVE and four exVE sub-regions.

**Figure S8.**
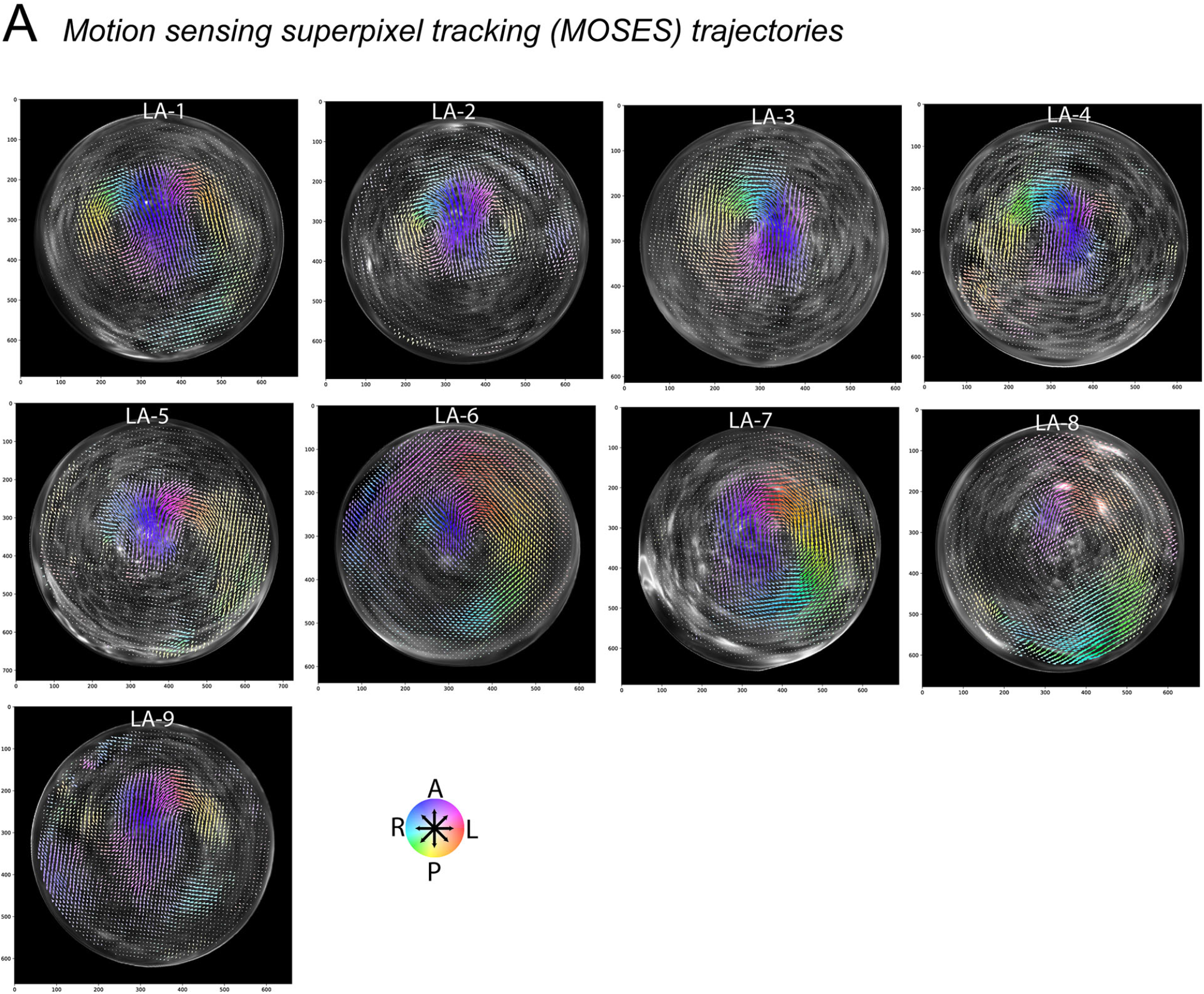
Motion Superpixel tracking analysis of Lifeact-GFP embryos. **A.** Polar projections of nine Lifeact-GFP embryos tracked by motion superpixel tracking, showing the mean velocity vector of each superpixel over the migration phase. The mean velocity vector is coloured by angle with colour intensity indicative of magnitude.

**Figure S9.**
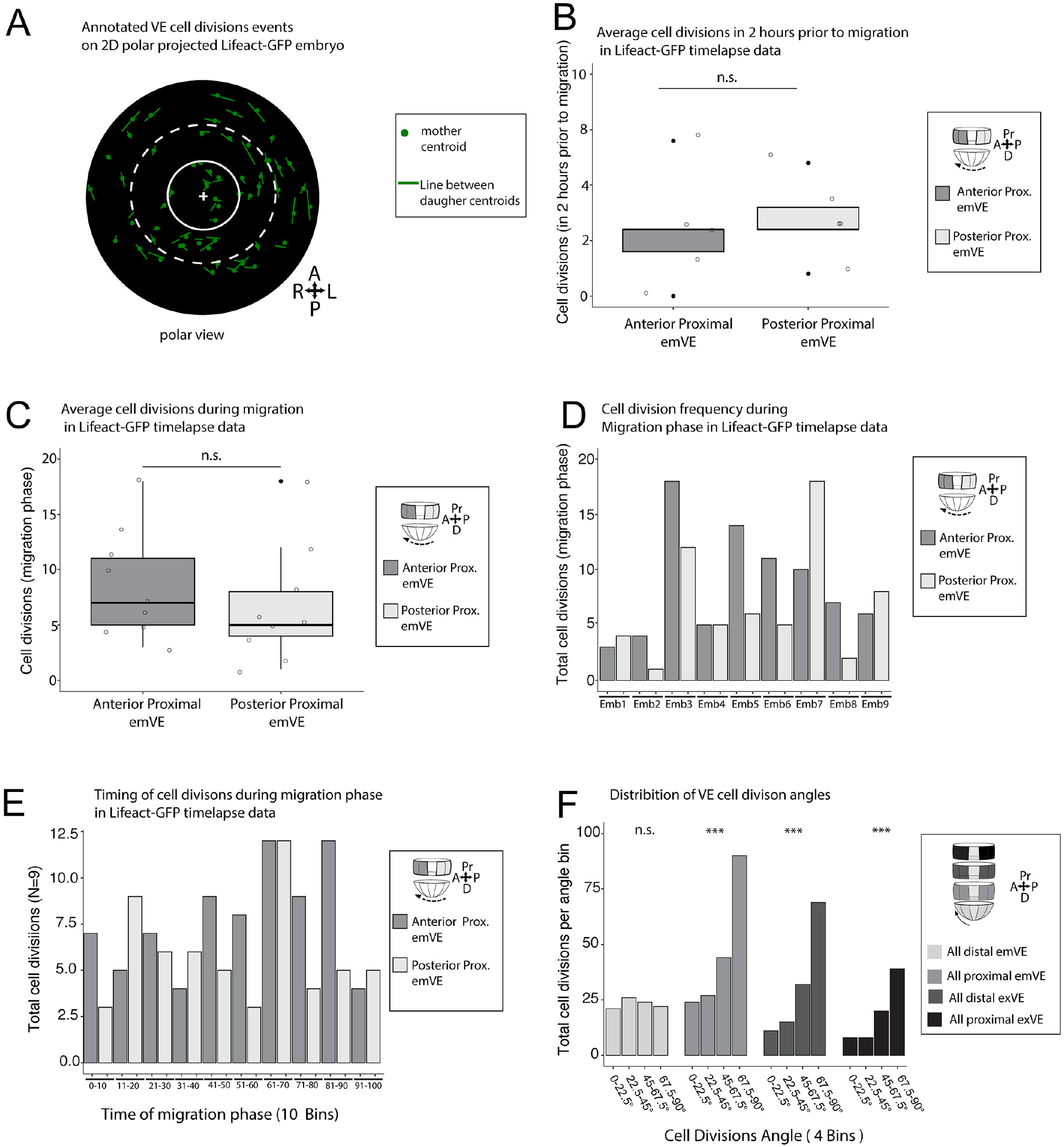
VE Cell Division Events in time-lapse imaged Lifeact-GFP embryos. **A.** Representative Lifeact-GFP embryo as 2D polar projection with all cell divisions events annotated. **B, C, D, E, F.** Cell division events from live imaged Lifeact-GFP embryos were quantified. **B.** Cell divisions in anterior- and posterior-proximal emVE during two hour period prior to migration (N=5 embryos). There was no significant difference in the average number of cell divisions between anterior proximal emVE (3.0 ± 2.6) and posterior proximal emVE (3.4 ± 1.8) (Student’s t-test, p=>0.05). **C.** Cell divisions in anterior- and posterior-proximal emVE during DVE migration were analysed (N=9 embryos). There was no significant difference in the average number of cell divisions between anterior proximal emVE (8.7 ± 5) and posterior proximal emVE (6.7 ±5.3) (Student’s t-test, p=>0.05). **D.** Cell division events in anterior- and posterior-proximal emVE during DVE migration per embryo. There were more posterior divisions than anterior in three embryos, while in the remaining six embryos there were either an equal number, or more divisions anteriorly than posteriorly. **E.** Cell divisions events in anterior- and posterior-proximal emVE during migration phase divided into ten bins show no difference in temporal ordering of divisions (N=9 embryos). **F.** Distribution of cell division angles in four regions of the VE during DVE migration. A 0° division angle means cells divide parallel to the proximal-distal axis a 90° angle = parallel to the radial axis. Distal emVE showed a distribution that was not significantly different than random (/2 test for expected probabilities p=>0.05), but all remaining regions showed a bias in cell division events along the radial axis (proximal emVE: *χ*^2^ for expected probabilities p=<0.001, distal exVE: *χ*^2^ for expected probabilities p=<0.001, proximal exVE: *χ*^2^ for expected probabilities p=<0.001).

**Figure S10.**
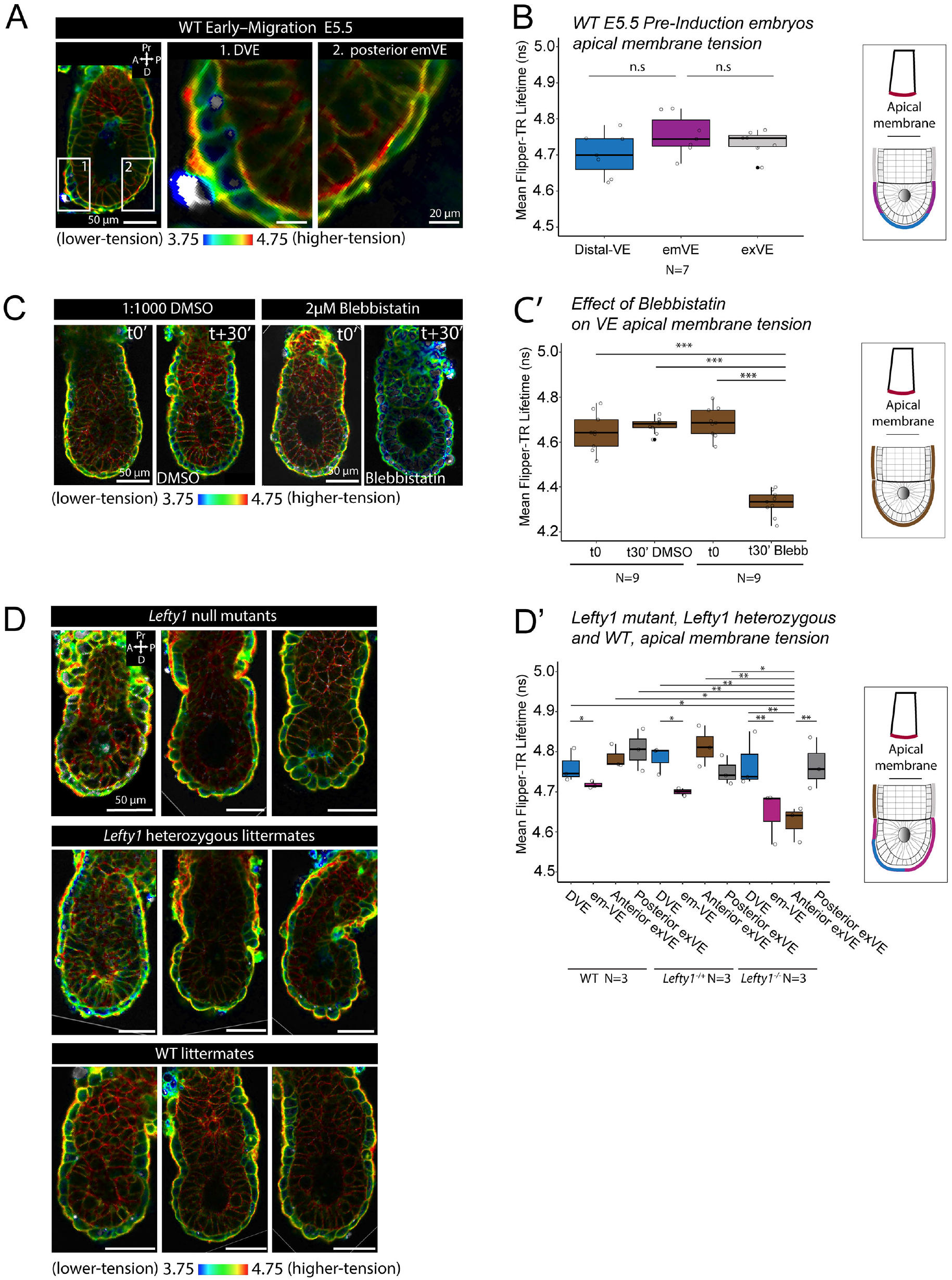
Fluorescence lifetime imaging. **A, B, C, D.** Fluorescence lifetime imaging microscopy (FLIM) and quantitation of FLIPPER-TR membrane tension reporter in mid-sagittal optical sections of wild type (WT) and *Lefty1* mutant embryos. **A.** Example image of an early migration stage embryo with DVE shifted to one-side. **B.** Apical lifetime in distal VE cells, emVE cells not belonging to the DVE and exVE in E5.5 pre-induction stage embryos (N=7) showed no significant differences between regions. **C.** E5.5 embryos were imaged to acquire a baseline (t0’) reading prior to treatment with an inhibitor of myosin (2 μM blebbistatin) or the solvent as control (1:1000 DMSO) and then re-imaged 30 minutes post-treatment (t+30’). **C’.** There was no significant difference in apical VE lifetime in the control DMSO treatment group, but the blebbistatin treated embryos showed a significant decrease in lifetime (one-way ANOVA, p=<0.001), Tukey’s HSD Test (p = < 0.001), showing that VE tension is actomyosin dependent. **D, D’.** Apical membrane lifetime of mid-migration *Lefty1*^−/−^ null (N=3), *Lefty1*^+/−^ heterozygous (N=3) and wild type (N=3) embryos. There were significant differences in tension based on genotype and region of the embryo, one-way ANOVA, p=<0.001, followed by Tukey’s HSD Test for specific comparisons. There was no difference in lifetime between anterior exVE, posterior exVE, and migrated DVE in WT or *Lefty1*^+/−^ heterozygous embryos (Tukey’s HSD Test, p=>0.05 for all comparisons). However in *Lefty1*^−/−^ mutant embryos, anterior exVE had a significantly lower lifetime than the DVE (Tukey’s HSD Test, p=<0.01), but not the posterior exVE (Tukey’s HSD Test, p=>0.05). Furthermore, the anterior exVE from *Lefty1*^−/−^ mutant embryos had significantly reduced tension when compared with the DVE, anterior exVE and posterior exVE of wild type and *Lefty1^-/+^* heterozygous embryos (Tukey’s HSD Test on *Lefty1*^−/−^ anterior exVE vs. wild type DVE (p=<0.05), wild type anterior exVE (p=<0.05), wild-type posterior exVE (p=<0.001), *Lefty1*^+/−^ heterozygous DVE (p=<0.01), *Lefty1*^+/−^ heterozygous anterior exVE (p=<0.001) and *Lefty1*^+/−^ heterozygous posterior exVE (p=<0.05). In all backgrounds, DVE had higher tension than emVE (Tukey’s HSD Test, p=>0.05).

**Table S1.**
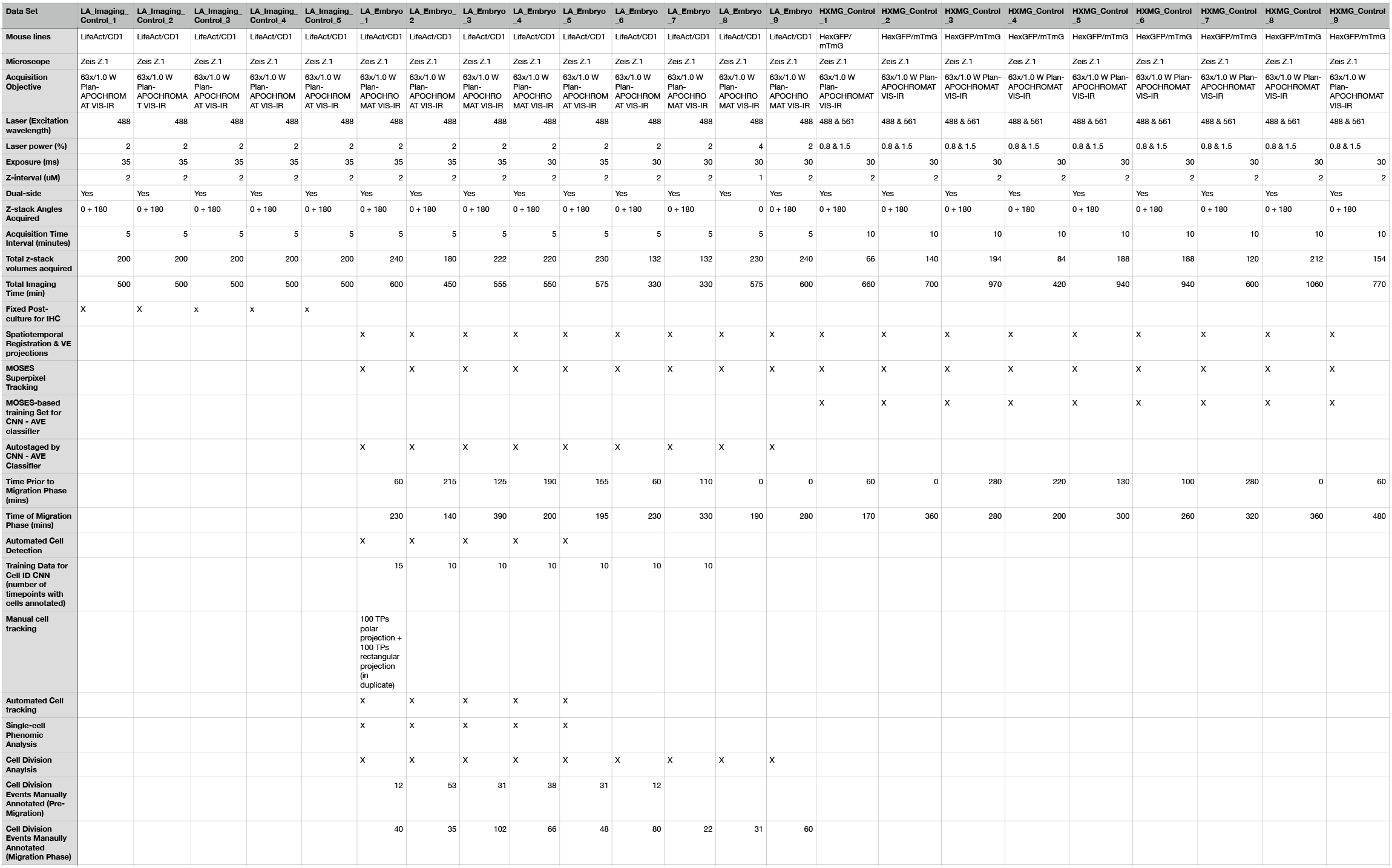
List of imaging data and analysis

**Table S2.** List of single-cell parameters for UMAP analysis

| Measure | Definition | Units | Analysis Level |
| --- | --- | --- | --- |
| **Area, A** | VE apical surface area | $\mu\text{m}^2$ | Cell |
| **Perimeter, P** | VE apical cell perimeter | $\mu\text{m}$ | Cell |
| **Shape Index, S** | $P/\sqrt{A}$ | - | Cell |
| **Aspect ratio** | Major axis length/minor axis length | - | Cell |
| **Cell height** | Mean distance between the VE apical surface and basal surface within the cell area | $\mu\text{m}$ | Cell |
| **# Cell neighbours** | Number of cells in immediate contact with the cell | - | Cell |
| **Instantaneous Anterior speed (VE), VVE** | Mean curvilinear speed of VE cell towards anterior | $\mu\text{m}/\text{min}$ | Cell |
| **Cumulative anterior distance** | Cumulative distance moved in the anterior migration direction relative to spatial position at start of migration stage | $\mu\text{m}$ | Cell |
| **Gauss surface curvature, KVE** | Mean 3D Gaussian curvature of VE layer surface within the cell area | $\mu\text{m}^{-2}$ | Local Tissue |
| **surface curvature, HVE** | Mean 3D Mean curvature of the apical surface of VE layer within the cell area | $\mu\text{m}^{-1}$ | Local Tissue |
| **Gauss basal surface curvature, KBasal** | Mean 3D Gaussian curvature of basal VE layer surface within the cell area | $\mu\text{m}^{-2}$ | Local Tissue |
| **Basal surface curvature, HBasal** | Mean 3D Mean curvature of the basal surface of VE layer within the cell area | $\mu\text{m}^{-1}$ | Local Tissue |
| **Apical area actin intensity** | Mean frame-normalised (zscore) LifeAct intensity within the cell area | - | Cell |
| **Apical perimeter actin intensity** | Mean frame-normalised (zscore) LifeAct intensity on the cell perimeter | - | Cell |

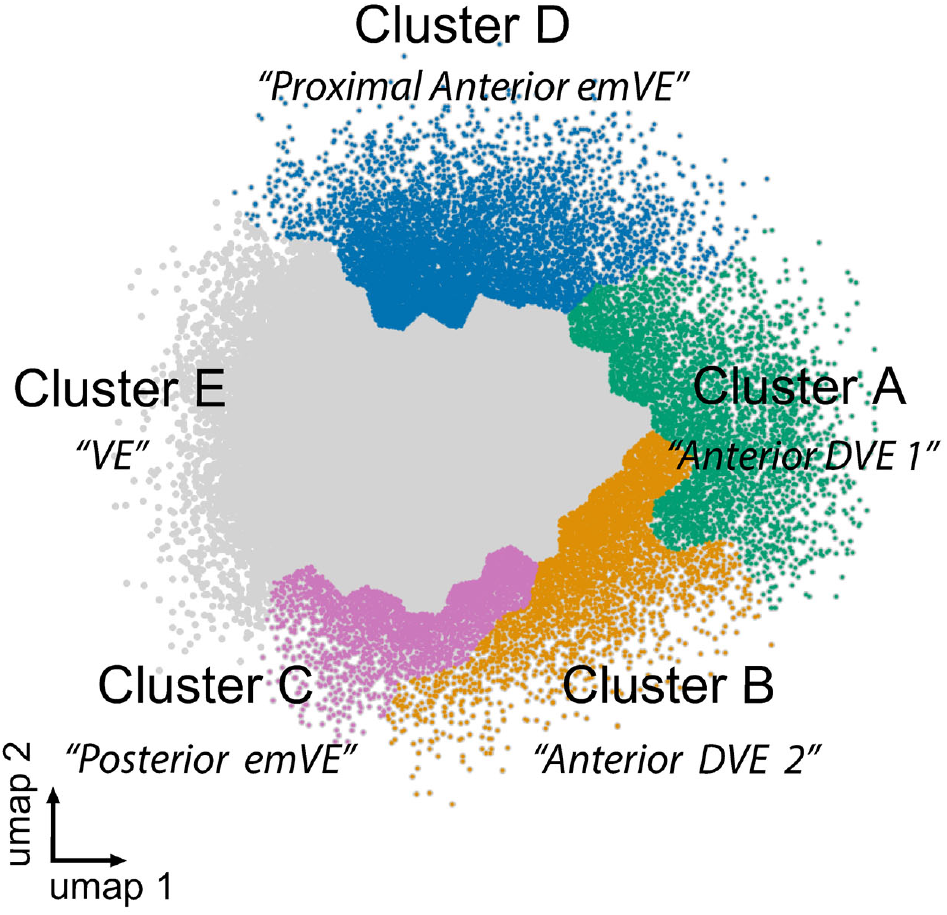

**Table S3.** UMAP cluster cell-statistics

| UMAP_cluster | A | B | C | D | E |
| --- | --- | --- | --- | --- | --- |
| #uniq_cells(tracks) | 490 | 604 | 503 | 599 | 2028 |
| #cell_instances(points) | 4939 | 4668 | 3773 | 8434 | 70087 |
| mean_Area | 80.52346857 | 116.6191976 | 207.9843496 | 189.0047956 | 211.9836127 |
| std_Area | 34.86962487 | 40.73001707 | 74.14555621 | 71.48139029 | 79.10606223 |
| mean_Perimeter | 41.74987103 | 46.51516565 | 63.29230273 | 88.41735758 | 75.959465 |
| std_Perimeter | 10.80886274 | 7.81106195 | 10.87329918 | 22.26876227 | 18.74232654 |
| mean_Shape_Index | 4.623896676 | 4.285127455 | 4.347583469 | 6.287724275 | 5.12587973 |
| std_Shape_Index | 0.51769817 | 0.224406052 | 0.308586131 | 0.999005134 | 0.775179415 |
| mean_Eccentricity | 1.653645121 | 1.29261722 | 1.222860135 | 2.461113106 | 1.526654629 |
| std_Eccentricity | 0.457465217 | 0.20631402 | 0.151629021 | 0.856780185 | 0.364529915 |
| mean_Thickness | 22.46898971 | 17.90128155 | 15.52420732 | 23.08539708 | 19.81309955 |
| std_Thickness | 4.382465709 | 3.984125083 | 3.852794722 | 4.130163375 | 3.957381623 |
| mean_Num_Neighbors | 4.754606196 | 5.051842331 | 5.636628677 | 5.515532369 | 5.743433162 |
| std_Num_Neighbors | 0.909398711 | 0.830731262 | 0.845752694 | 1.077089394 | 0.955248865 |
| mean_A_P_speed_VE | 0.130708615 | 0.115459743 | 0.065148348 | 0.016153396 | 0.013766578 |
| std_A_P_speed_VE | 0.207569044 | 0.198095563 | 0.159571964 | 0.14251743 | 0.126601846 |
| mean_cum_A_P_VE_speed | 2.676548187 | 1.3748286 | 0.670067231 | 0.76593717 | 0.183659292 |
| std_cum_A_P_VE_speed | 3.381420368 | 2.45493328 | 1.744130637 | 1.959203188 | 1.117753818 |
| mean_Mean_Curvature_VE | 0.018608583 | 0.02127803 | 0.018636241 | 0.011732662 | 0.014163141 |
| std_Mean_Curvature_VE | 0.007213856 | 0.006425515 | 0.005077823 | 0.005097735 | 0.004703488 |
| mean_Mean_Curvature_Epi | 0.023223832 | 0.026877937 | 0.024867422 | 0.016437137 | 0.020007063 |
| std_Mean_Curvature_Epi | 0.008518081 | 0.007893641 | 0.00641074 | 0.010816232 | 0.007950077 |
| mean_Gauss_Curvature_VE | 0.000192541 | 0.00045322 | 0.000328515 | -0.000722758 | 2.99E-07 |
| std_Gauss_Curvature_VE | 0.004523865 | 0.000312234 | 0.000230596 | 0.008839429 | 0.002662359 |
| mean_Gauss_Curvature_Epi | 0.000420801 | 0.000683199 | 0.000549453 | -0.000408465 | 7.13E-05 |
| std_Gauss_Curvature_Epi | 0.001366855 | 0.000531844 | 0.000430972 | 0.004632281 | 0.001473701 |
| mean_norm_apical_actin | 1.753306628 | 0.757132514 | 0.053330851 | 0.855594778 | 0.34379363 |
| std_norm_apical_actin | 0.998529035 | 0.627348763 | 0.333609662 | 0.535353585 | 0.443457129 |
| mean_norm_perimeter_actin | 1.757957566 | 0.930002378 | 0.276976641 | 0.950897818 | 0.518107186 |
| std_norm_perimeter_actin | 0.913554768 | 0.630632239 | 0.359617951 | 0.524461312 | 0.450450051 |

## Notes

### Competing Interest Statement

The authors have declared no competing interest.

